# Virus-free CRISPR knock-in of a chimeric antigen receptor into *KLRC1* generates potent GD2-specific natural killer cells

**DOI:** 10.1101/2024.02.14.580371

**Authors:** Keerthana Shankar, Isabella Zingler-Hoslet, Lei Shi, Varun Katta, Brittany E. Russell, Shengdar Q. Tsai, Christian M. Capitini, Krishanu Saha

## Abstract

Natural killer (NK) cells are an appealing off-the-shelf, allogeneic cellular therapy due to their cytotoxic profile. However, their activity against solid tumors remains suboptimal in part due to the upregulation of NK-inhibitory ligands, such as HLA-E, within the tumor microenvironment. Here, we utilize CRISPR-Cas9 to disrupt the *KLRC1* gene (encoding the HLA-E-binding NKG2A receptor) and perform non-viral insertion of a GD2-targeting chimeric antigen receptor (CAR) within NK cells isolated from human peripheral blood. Genome editing with CRISPR/Cas9 ribonucleoprotein complexes yields efficient genomic disruption of the *KLRC1* gene with 98% knockout efficiency and specific knock-in of the GD2 CAR transgene as high as 23%, with minimal off-target activity as shown by CHANGE-Seq, in-out PCR, and next generation sequencing. *KLRC1*-GD2 CAR NK cells display high viability and proliferation, as well as precise cellular targeting and potency against GD2^+^ human melanoma cells. Notably, *KLRC1*-GD2 CAR NK cells overcome HLA-E-based inhibition by HLA-E-expressing, GD2^+^ melanoma cells. Using a single-step, virus-free genome editing workflow, this study demonstrates the feasibility of precisely disrupting inhibitory signaling within NK cells via CRISPR/Cas9 while expressing a CAR to generate potent allogeneic cell therapies against HLA-E^+^ solid tumors.

## Introduction

Chimeric antigen receptor (CAR) T cells, which involve genetic engineering by lentiviral or retroviral vectors of autologous peripheral blood mononuclear cells with synthetic receptors to CD19 or BCMA, have revolutionized treatment of blood cancers.^1^ Despite high response rates ranging from 50-90%, CAR T cells have been shown to induce life threatening toxicities like cytokine release syndrome (CRS) and immune effector cell-associated neurotoxicity syndrome (ICANS) upon infusion.^2–5^ As an alternative approach, CAR natural killer (NK) cells have displayed outstanding results in the clinic for treating lymphoid malignancies, with 30% of patients achieving a complete response with no instances of CRS or ICANS.^6,7^ CAR NK cells have failed to produce comparable results in solid tumors in part due to the upregulation of NK inhibitory checkpoints by the tumor.^8^ One of these ligands, the nonclassical human leukocyte antigen (HLA) class I molecule HLA-E, is upregulated across several tumors and ligates the inhibitory NKG2A receptor as a heterodimer with CD94, limiting NK effector function.^9,10^ NKG2A effectively serves as an NK immune checkpoint molecule that abrogates effector functions, such as cytotoxicity, in part by disrupting actin-NKG2D synapses.^11^ Blockade of NKG2A by monoclonal antibodies, or ablation of the receptor through genomic engineering, has been shown to improve NK effector function *in vivo* as well as in clinical trials.^10,12–14^ Combinatorial therapies utilizing blockade of NKG2A with CAR modification present a large potential for enhancing NK cell responses against solid tumors but have been unexplored to date.

As a component of the innate immune system, NK cells have high sensitivity to exogenous DNA — making genomic engineering of NK cells with viral vectors challenging, often resulting in low transduction efficiencies (<1-20%) and poor cell viability.^15,16^ In addition, viral vectors integrate randomly, leading to variability of transgene expression, with some risk of insertional mutagenesis.^17,18^ Electroporation-based methods can deliver mRNA payloads in a non-viral manner quite efficiently, but only result in transient gene expression of the mRNA. Alternatively, delivery of transposable vector elements into NK cells results in multiple potential integration sites, also raising insertional mutagenesis risks.^19,20^ None of these approaches allow for a genome edit in which a specific gene can be targeted for knock-out or the precise knock-in of a transgene. Non-viral genome editing of NK cells with CRISPR genome editors has been achieved with high efficiency for knockouts, but with low efficiency for knock-in of large transgene insertions. For example, efficiencies range from 3-17% for fluorescent proteins like GFP (1.5 kb insertion) and mCherry (2.1 kb insertion) transgenes, and less than 10% for a CAR (1.4 kb insertion).^21–24^

Here, we describe a robust workflow using nonviral CRISPR-Cas9 ribonucleoproteins (RNP) to generate CAR NK cells against the diasialoganglioside GD2, an antigen expressed on a variety of solid tumors such as neuroblastoma, osteosarcoma and melanoma.^25^ We identify several parameters influencing editing efficiencies to ensure efficient knock-out of the *KLRC1* gene, encoding the inhibitory NKG2A receptor, and knock-in of an anti-GD2 CAR transgene at this locus. *KLRC1*-CAR NK cells are highly viable and proliferative, with minimal off-target editing. In a GD2^+^ melanoma model, the *KLRC1*-CAR NK cells display improved cytotoxicity and secretion of IFNγ *in vitro* when compared to unedited NK cells. This streamlined, virus-free engineering strategy has high potential to generate allogeneic CAR NK cells for the treatment of HLA-E^+^ solid tumors.

## Results

### CRISPR editing of the NKG2A receptor

Our editing strategy to improve NK potency against solid tumors involves knocking out the NKG2A receptor, which inhibits NK cell cytotoxicity upon engagement with the HLA-E ligand on the tumor (**Figure 1A**), while simultaneously knocking-in a CAR conferring specificity to the GD2 antigen. To knock out (KO) the NKG2A receptor, we designed three single-guide RNAs (sgRNAs) against several common coding exons in the human *KLRC1* gene (**Figure 1B**). Ribonucleoproteins (RNPs) were generated upon the complexation of these sgRNAs with *Sp*Cas9 to generate double-stranded DNA breaks at these targets within *KLRC1*. As a proof of concept, NK-92 cells, an IL-2 dependent human NK cell line were nucleofected with RNPs containing one of these sgRNAs and assessed for insertions or deletions (indels) around the targeted site after two weeks of cell culture. Next-generation sequencing (NGS) of PCR amplicons derived from genomic DNA showed a high level of indel formation (98%) in the RNP-treated cells for our top-performing guide RNA (gRNA1) targeting exon 3 of *KLRC1*. These indels were absent in the untransfected (UTF) control sample (**Figure 1C**). Moreover, the indels were centered exactly at the predicted site of DNA double-strand cleavage by *Sp*Cas9 (**Figure 1D**). While UTF NK-92 cells are highly cytotoxic and have been used as an allogeneic cell therapy for cancer for over 20 years, including CAR-modified NK-92s, their cancerous origin requires them to be irradiated before infusion to prevent proliferation, limiting their *in vivo* persistence.^26–30^ Additionally, their lack of CD16 expression renders them incapable of antibody dependent cellular cytotoxicity (ADCC).^31^ For these reasons, we then elected to use *ex vivo* expanded primary NK cells isolated from human peripheral blood for downstream development.

**Figure 1.**
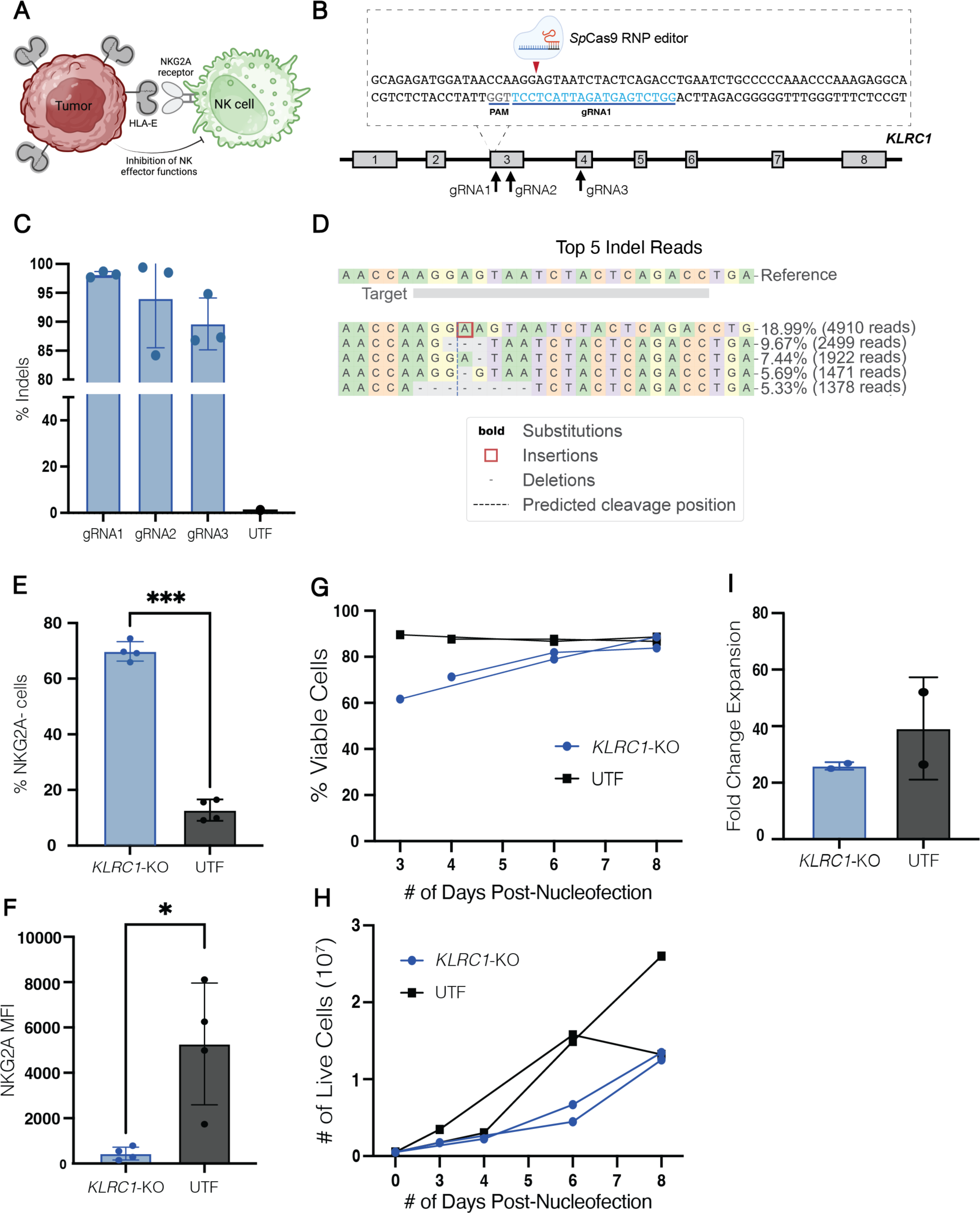
Cas9 ribonucleoprotein-based strategy yields high *KLRC1* knock-out efficiency. **(A)** Schematic showing the interaction between NK cells and the tumor. The HLA-E ligand is often upregulated on the surface of tumors and inhibits NK effector functions by binding the inhibitory NKG2A receptor. **(B)** Schematic depicting knock-out strategy. sgRNA1 (in blue), in complex with the spCas9 editor, targets the third exon of the *KLRC1* gene. The locations of the three gRNA sequences are indicated by black arrows in sequential order. **(C)** Percent of indel formations for each of the three sgRNAs in NK-92 cells at the *KLRC1* gene. Data is shown for three replicates for one donor (n = 1). (**D)** Quantification of the top five allelic editing outcomes of gRNA1 as analyzed by CRISPResso2.^46^ **(E)** Primary PB-NK cells were nucleofected with RNP complexes targeting *KLRC1.* Percent of NKG2A^−^ cells is shown in comparison to the UTF population, as measured by flow cytometry. Data is shown as the average of two replicates across four donors (n = 4). **(F)** MFI of the NKG2A receptor on the cell surface of *KLRC1*-KO and UTF cells. Data is shown as the average of two replicates across four donors (n = 4). **(G and H)** Cell viability and proliferation of *KLRC1-*KO and UTF cells after nucleofection. Data is shown as the average of two replicates across two donors (n = 2). **(I)** Calculated fold expansion of *KLRC1-*KO and UTF cells eight days after nucleofection. A two-tailed paired *t* test and one-way ANOVA followed by Tukey’s multiple comparisons test were used to test for statistical significance. *, p ≤ 0.05; ***, p ≤ 0.001; ns, p ≥ 0.05. PB, peripheral blood; RNP, ribonucleoprotein; UTF, untransfected; MFI, mean fluorescence intensity.

NK cells were isolated from the whole blood of healthy donors using density-based negative selection and expanded with irradiated K562-mb15-41BBL feeder cells supplemented with IL-2 and IL-15. After four days of culture, NK cells were nucleofected with an RNP complex targeting the *KLRC1* gene and expanded for one week. The edited cells were then analyzed for NKG2A expression using flow cytometry alongside a donor matched UTF control population. Within the RNP-treated cells, an average of 70% NK cells were negative for NKG2A on their cell surface, compared to 13% of NK cells within the UTF control (**Figure 1E**). The loss of surface NKG2A protein was further corroborated by a decrease in the mean fluorescence intensity (MFI) signal in the RNP-treated cells when compared to the UTF cells (**Figure 1F**). *KLRC1*-KO NK cells maintained high viability throughout expansion and proliferated consistently (**Figures 1G, 1H**). The fold expansion of *KLRC1*-KO NK cells was not significantly different than that of their UTF counterparts (**Figure 1I**).

### Non-viral transgene knock-in into *KLCR1*

To assess whether this editing strategy could also be used to insert long transgenes in primary NK cells, we adapted a virus-free editing technique developed for primary T cells that harnesses homology-directed repair (HDR) at double-stranded DNA breaks.^32^ We nucleofected NK cells with an RNP complex together with a linear, double stranded DNA (dsDNA) template encoding a long transgene (2709 bp): the signaling-inert mCherry gene (*KLRC1*-No CAR) (**Figure 2A**). Since HDR is most likely to occur when cells are actively proliferating and entering the S- or G2-phases of the cell cycle,^33^ we first determined the optimal time to nucleofect the cells to ensure the highest knock-in efficiencies. We nucleofected NK cells on days 4-7 of expansion and assessed the level of NKG2A knockout and mCherry expression by flow cytometry. mCherry integration was highest when cells were nucleofected on day 4 with an average of 4% cells exhibiting mCherry expression and 68% cells demonstrating NKG2A knockout (**Figures 2B, 2C**). However, minimal cell growth was observed for the first six days after nucleofection (**Figure 2D**). On days 5, 6, and 7, the percentage of mCherry integration was significantly decreased, to roughly 1% (**Figure S1A**). In contrast, these later timepoints increased the percentage of NKG2A^−^ cells, with knockout rates approaching 80% (**Figure S1B**).

**Figure 2.**
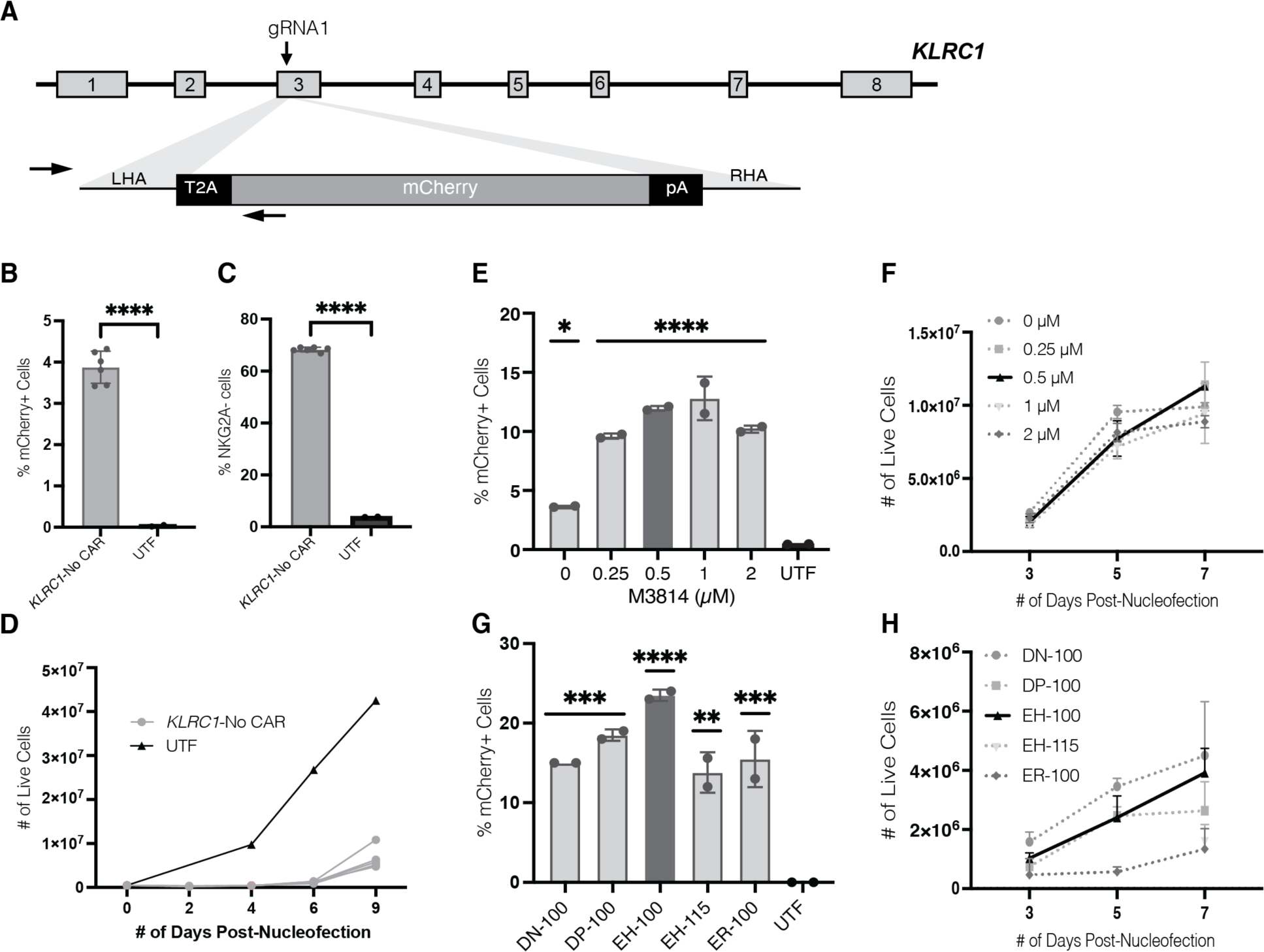
Non-viral transgene knock-in into primary NK cells. See also Figure S1. **(A)** Schematic depicting virus-free insertion of the mCherry fluorescence gene at the third exon of the *KLRC1* locus. **(B and C)** NK cells were nucleofected on day 4 of expansion. Percentage of mCherry^+^ cells and NKG2A^−^ cells is shown, as measured by flow cytometry, one week after nucleofection. Data is shown for six replicates for one donor (n = 1). **(D)** Cell proliferation of *KLRC1*-No CAR cells from B. **(E)** NK cells were incubated with varying concentrations of M3814 for 24 hours after nucleofection. The percent of mCherry^+^ cells was measured by flow cytometry one week after nucleofection. Data is shown for two replicates for one donor (n = 1). **(F)** Expansion of mCherry^+^ cells after a 24-hour incubation with M3814. **(G)** NK cells were nucleofected on day 4 using different pulse programs, followed by M3814 treatment. Data is shown for two replicates for one donor (n = 1). **(H)** Expansion of NK cells following nucleofection with different pulse programs. Statistical significance was calculated using a two-tailed unpaired *t* test (B and C) and by ordinary one-way ANOVA. Dunnett’s multiple comparison test was used as the post-test (E and G). *, p ≤ 0.05; **, p ≤ 0.01; ***, p ≤ 0.001; ****, p ≤ 0.0001. ANOVA, analysis of variance.

To increase HDR, the DNA-dependent protein kinase (DNA-PK) inhibitor, M3814, was added to the cell culture medium, as others had previously shown it to improve knock-in rates of primary human T cells by inhibiting non-homologous end joining (NHEJ) DNA repair events.^22^ NK cells were nucleofected on day 4 with the RNP complex and mCherry dsDNA, and incubated with M3814 at various concentrations. Nucleofected cells were expanded for one week and assayed by flow cytometry for NKG2A knock-out and transgene expression. Incubation with M3814 had no effect on NKG2A knock-out efficiencies, with the percent of NKG2A^−^ cells remaining around 75% (**Figure S1C**). In contrast, knock-in efficiencies significantly improved when nucleofected cells were treated with any concentration of M3814 compared to no M3814 treatment (**Figure 2E**). Cells treated with either 0.5 μM or 1 μM of M3814 had the highest percentage of transgene expression at 11.9% and 12.8%, respectively. Moreover, treatment with M3814 did not affect cell expansion after nucleofection, with M3814-treated cells growing comparably to untreated cells (**Figure 2F**). For subsequent studies, 0.5 μM M3814 was chosen due to the increased editing efficiency and high cell yield at one week after nucleofection.

As our strategy depends on electroporation for the efficient delivery of payload into NK cells, we next optimized the pulse codes to strike a balance between knock-in efficiency and cell viability. For the Lonza 4D nucleofection system, we selected two programs (DN-100 and EH-100) that have resulted in high RNP based knock-out efficiencies and plasmid-based GFP knock-in efficiencies while maintaining cell viability in primary NK cells.^21^ Additionally, we included two pulse codes (DP-100 and ER-100) that were recommended by the manufacturer as being similar to DN-100 but potentially yielding higher efficiencies. As a control, we also tested the EH-115 program used to nucleofect primary human T cells.^32,34^ Consistent with the M3814 optimization, there were no significant differences in NKG2A knock-out efficiencies across the different programs, with the percent of NKG2A^−^ cells remaining between 76-85% (**Figure S1D**). The EH-100 program yielded the highest percentage of transgene-positive NK cells at 23.5% (**Figure 2G**). Finally, we assessed cell viability and growth throughout expansion for each of the pulse code programs to maximize cell yield. Throughout expansion, the DN-100 program resulted in higher NK cell yields, while the ER-100 program yielded the lowest number of viable NK cells. The EH-100 pulse code was chosen for subsequent experiments due to the significant improvement in transgene editing and sufficient expansion of the edited NK cells (**Figure 2H**).

### Characterization of *KLRC1-*CAR NK cells

Having developed an efficient non-viral *KLRC1* gene editing protocol for NK cells (**Figure 3A**), we applied our strategy to generate GD2-targeting CAR NK cells (*KLRC1-*CAR), with an anti-GD2 CAR composed of the following: an anti-GD2 single chain variable fragment (scFv; 14g2a); CD28 transmembrane and endodomain; 41BB costimulatory domain and CD3z cytotoxicity domain (**Figure 3B**). Though the CAR transgene (3180 bp) was approximately 500 bp larger than the mCherry transgene (2709 bp), the CAR knock-in efficiency (average of 20.0%) was similar to that of mCherry (average of 20.1%) (**Figures 3C, 3D**). The MFI for the CAR transgene also significantly increased compared to the donor matched UTF control (**Figure 3E**). In line with previous outcomes, the NKG2A knock-out efficiencies remained largely unchanged when NK cells received the CAR transgene but were increased marginally with the delivery of the mCherry transgene (**Figure S2A**). The NKG2A MFI levels, however, had no significant differences across the three groups that were edited via CRISPR (**Figure S2B**). After profiling for canonical CD56 and CD16 NK cell surface markers in the cell products, the fractions of CD56^+^CD16^+^ and CD56^+^CD16^−^ cells among the edited and untransfected groups remained unchanged through the editing and expansion workflows (**Figure 3F).**

**Figure 3.**
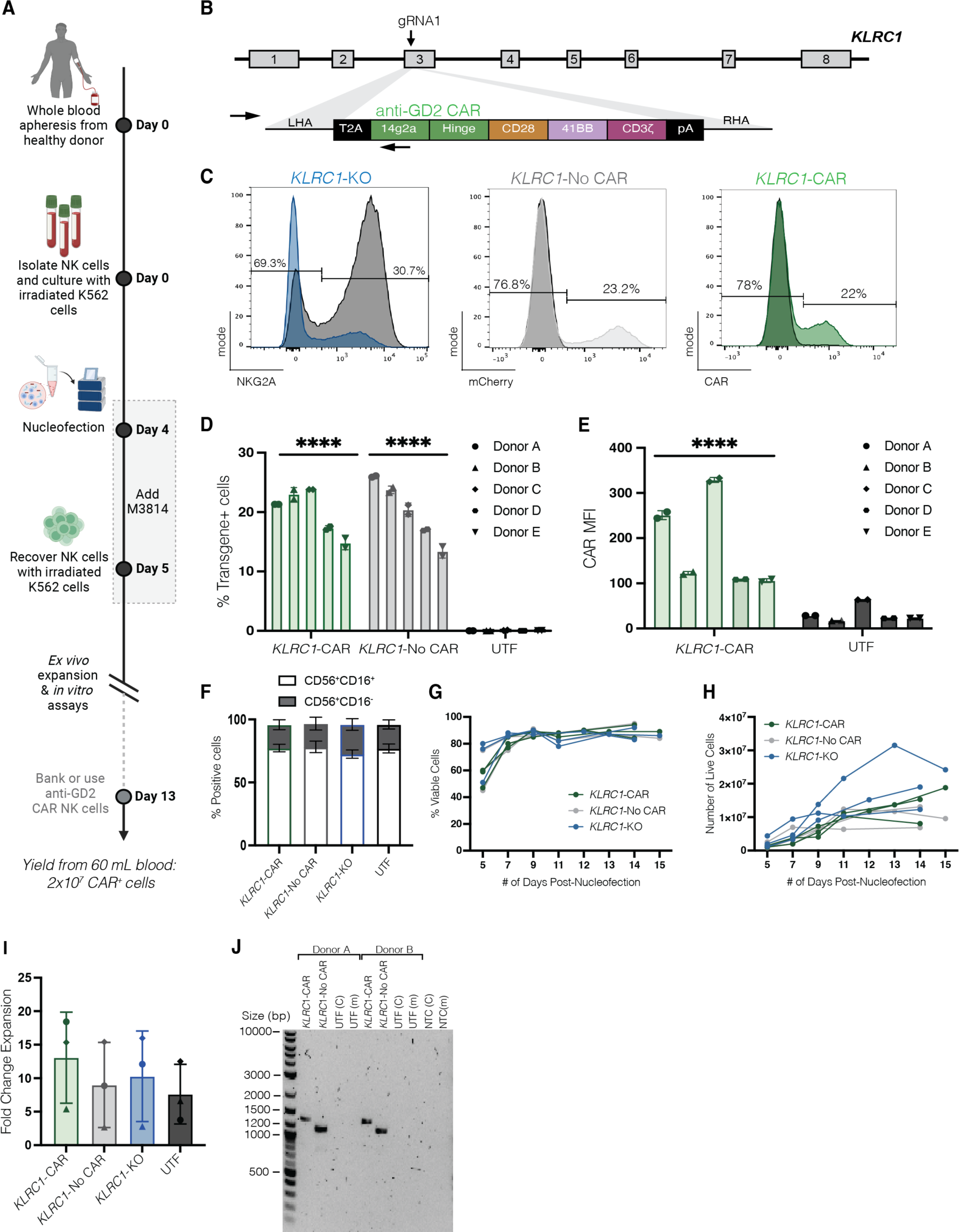
Characterization of non-viral *KLRC1-*CAR NK cells. See also Figure S2. **(A)** Proposed manufacturing timeline for *KLRC1*-CAR NK cells. NK cells are isolated from whole blood of healthy donors using negative selection and activated with irradiated K562-mb15-41BBL cells (1 NK: 2 K562) along with IL-2 and IL-15. On day 4, NK cells are nucleofected with Cas9 RNP and dsDNA HDRT using program EH-100, followed by a 24-hour treatment of 0.5uM M3814. Nucleofected NK cells are recovered with addition of irradiated K562-mb15-41BBL cells (1:1), supplemented with IL-2 and IL-15. *KLRC1*-CAR cells are expanded *ex vivo* for downstream analysis and cell banking. The calculated cell yield reflects the estimated number of *KLRC1-*CAR^+^ NK cells derived from 60 mL of blood on Day 13 of expansion. **(B)** Schematic depicting virus-free insertion of an anti-GD2 CAR at the third exon of the *KLRC1* locus. **(C)** Representative histogram plots showing knockout of NKG2A and knock-in of CAR or mCherry transgenes one week after nucleofection. X-axis describes protein expression levels. Y-axis describes frequency after normalization to the mode. UTF samples are shown as black histograms in all plots. The percent shown for each gate represents protein levels in the *KLRC1*-KO (left), *KLRC1-*No CAR (middle), and *KLRC1-*CAR (right) edited samples. **(D and E)** CAR and mCherry knock-in efficiencies, and MFI levels are shown. Data is shown as an average of two replicates across five donors (n = 5). **(F)** The percent of CD56^+^CD16^+^ and CD56^+^CD16^−^ cells in each cell population is shown. Data is shown as an average of two replicates across two donors (n = 2). **(G and H)** Cell viability and cell expansion of *KLRC1-*CAR, *KLRC1-*No CAR, and *KLRC1-*KO NK cells after nucleofection are shown. During expansion, multiple nucleofection reactions were pooled to form one sample per group. Data is normalized to the number of reactions for each of the three donors (n = 3). **(I)** Calculated fold change expansion of *KLRC1*-CAR, *KLRC1-*No CAR, and *KLRC1-*KO NK cells from H is shown in comparison to donor-matched UTF cells, with each donor indicated by a different shape (circle, triangle, diamond). **(J)** In-out PCR showing on-target integration of the CAR and mCherry transgenes across two donors. The UTF samples were amplified using the CAR (UTF (C)) and mCherry (UTF (m)) primer pairs, as controls. NTC refers to the PCR reaction performed without the genomic DNA. The primers are indicated by the black arrows on each template in the schematics of Figures 2A and 3B. Statistical significance was calculated using an ordinary one-way ANOVA. Post-tests were performed using the uncorrected Fisher’s LSD (D), Sidak’s multiple comparisons test (E), and Dunnett’s multiple comparisons test (I). *, p ≤ 0.05; ****, p ≤ 0.0001; ns, p ≥ 0.05. RNP, ribonucleoprotein; dsDNA, double stranded DNA; HDRT, homology directed repair template; CAR, chimeric antigen receptor; MFI, mean fluorescence intensity; UTF, untransfected; PCR, polymerase chain reaction; NTC, non-template control; ANOVA, analysis of variance.

We next confirmed that insertion of the CAR into the *KLRC1* locus did not affect NK cell viability and proliferation by tracking cell expansion after nucleofection. When comparing the various cell products immediately after nucleofection, poor cell expansion correlated with the length of the exogenous dsDNA template within the first week, with *KLRC1-*CAR NK cells having the lowest fold expansion, followed by *KLRC1-*No CAR, and *KLRC1-*KO NK cells (**Figures S2C, S2D, S2E)**. However, after recovery from electroporation, *KLRC1-*CAR NK cells were highly viable and proliferated at a similar rate to *KLRC1-*No CAR and *KLRC1-*KO NK cells in the log growth phase (**Figures 3G, 3H)**. The fold change expansion for all edited groups beginning on day 5 was comparable to that of donor matched UTF NK cells with no significant differences (**Figure 3I**).

We next characterized the genomic outcomes within the *KLRC1*-CAR NK cell product by checking for on- and off-target modifications, and integration of our transgenes throughout the genome. First, an “in-out” PCR for the intended transgene integration at the *KLRC1* on-target site was performed on genomic DNA extracted from edited NK cells. Using primers that were specific to the transgene and the *KLRC1* locus, we showed site-specific amplification of our GD2 CAR and mCherry transgenes, that were not present in the UTF cells (**Figure 3J)**. The size of the bands are approximately 1300 bases (CAR) and 1100 bases (mCherry), exactly as predicted from the proper on-target integration of each transgene (**Table S1**).

We also examined the genome for off-target activity from the *KLRC1* gRNA via CHANGE-seq. This method profiles the entire human genome for double-stranded DNA breaks induced by a Cas9 RNP. Moderate off-target activity in the CHANGE-seq assay was observed largely within intergenic or intron regions of the genome, with 10 sites having off-target nuclease activity within exons and 8 sites having off-target activity near a transcription start site (TSS) (**Figure S3, Table S2**). The CHANGE-seq specificity ratio, calculated by dividing the number of on-target reads by the sum of total reads, for our sgRNA is 0.038. As expected, small mismatches distal to the protospacer-adjacent motif (PAM) of Cas9 were present in the top nominated off-target sites (**Figure 4A**).

**Figure 4.**
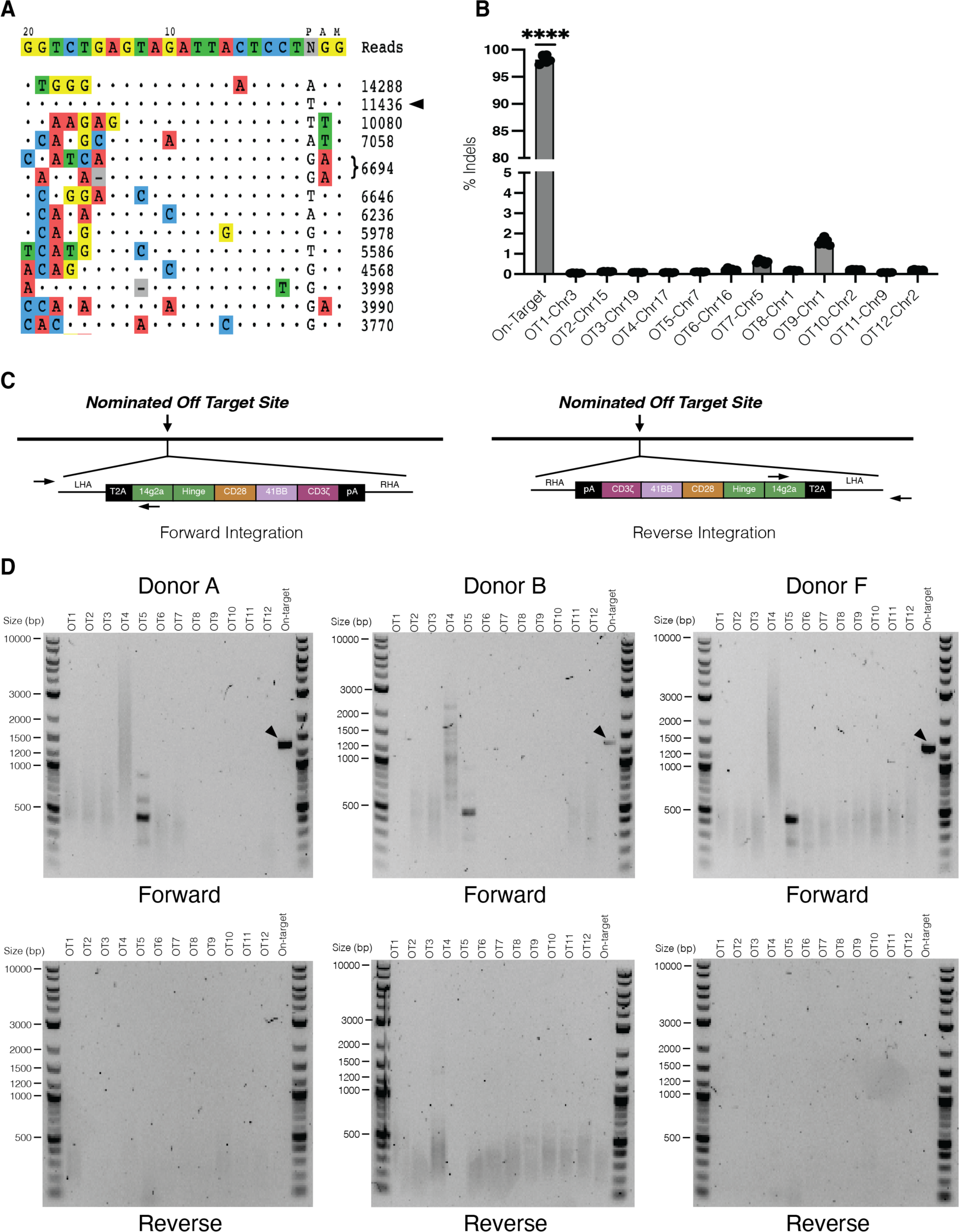
Genome-wide off-target analysis. See also Figure S3. (**A**) Nominated off-target sites detected by CHANGE-Seq are organized by number of read counts. The on-target site is the second line on the list, indicated by a black arrow. Base pair mismatches between the sgRNA and the off-target sites are shown as colored nucleotides (n = 1). (**B**) Editing across the top 12 nominated off-target sites was assayed via deep sequencing using the rhAmpSeq system. Data is shown as mean (SD) across three donors, with two replicates per donor (n = 3). (**C**) Schematic depicting locations of the forward and reverse primers for in-out PCRs for the CAR gene at an off-target site. The left schematic represents CAR integration in the 5’ – 3’ orientation, while the right represents integration in the 3’ – 5’ orientation, with the primers indicated by black arrows. (**D**) In-out PCRs were performed on genomic DNA extracted from *KLRC1*-CAR NK cells to assess for off-target integration in the forward (top) and reverse (bottom) orientations across three donors. The on-target CAR integration is indicated by the black arrow. PCR, polymerase chain reaction; CAR, chimeric antigen receptor.

To further evaluate potential off-target genomic modifications, we performed deep sequencing around each of the top 12 off-target sites nominated by CHANGE-seq (>3500 reads). We completed NGS via the rhAmpSeq analysis system to multiplex the genomic analysis for on- and off-target sites from the same genomic DNA sample. By rhAmpSeq, we observed high indel formation only at the on-target site with minimal editing across all 12 off-target sites (**Figure 4B**). A small number of indel-containing reads (<2%) had a broad single-nucleotide mutation spectrum around each of the nominated off-target sites, reflecting the background level of PCR-based or sequencing-based errors in the assay and the limit of detection of this assay.

Finally, to evaluate the potential off-target integration of our CAR, we performed in-out PCR on each of the top 12 off-target sites across three donors to check for unintended CAR integration, in either the 5’-3’ or 3’-5’ orientation (**Figure 4C)**. No bands of the expected size were observed (**Table S1**) in any of the off-target sites, indicating that there is no off-target integration of the CAR within the limit of detection for this assay (**Figure 4D**). The 500 bp band at site 5 (**Figure 4D**, top, lane 6) is from non-specific binding of the primers as the expected amplicon size from off-target integration at this site would have been 1931 bp. As a positive control, the on-target site was included, and a band of approximately 1300 bp was observed (**Figure 4D**, top, lane 14), which corresponds to the expected amplicon size of 1349 bp, confirming successful on-target integration of the CAR within *KLRC1*.

### *KLRC1*-CAR NK cells are highly cytotoxic against GD2^+^ tumor cells

The specificity of *KLRC1*-CAR NK cell function was evaluated against GD2^+^ M21 melanoma cells (**Figure 5A**). We isolated pure populations of NKG2A^−^, CAR^+^, or mCherry^+^ cells using florescence activated cell sorting (FACS) to determine the effect of each gene modification on NK cytotoxicity. NK cells were co-cultured with M21 cells at a 5:1 effector-target ratio for 24 hours (**Figure 5B**). NK cytotoxicity was 6-10% higher in the *KLRC1*-CAR NK cells compared to *KLRC1*-No CAR, *KLRC1*-KO and UTF cells (**Figure 5C, left**). To demonstrate that the boost in cytotoxicity function arose from CAR binding, we added a monoclonal antibody to label the CAR target antigen, GD2, to the co-culture so that NK cell recognition of the tumor could occur independent of the CAR via CD16, triggering ADCC. Upon addition of the hu14.18K322A anti-GD2 monoclonal antibody, all NK cell products were highly cytotoxic (**Figure 5C, right**) and the differences in NK cell cytotoxicity between *KLRC1*-CAR, *KLRC1*-No CAR, *KLRC1*-KO NK cells were erased in ADCC promoting conditions. Upon comparing non-ADCC and ADCC conditions, we attribute that the boost in the cytotoxic activity of the *KLRC1*-CAR NK cells compared to other cells is specific to the GD2 antigen. The precise knock-in of the CAR increases the cytotoxicity of the *KLRC1-*CAR NK cells to the maximal level in this co-culture assay.

**Figure 5.**
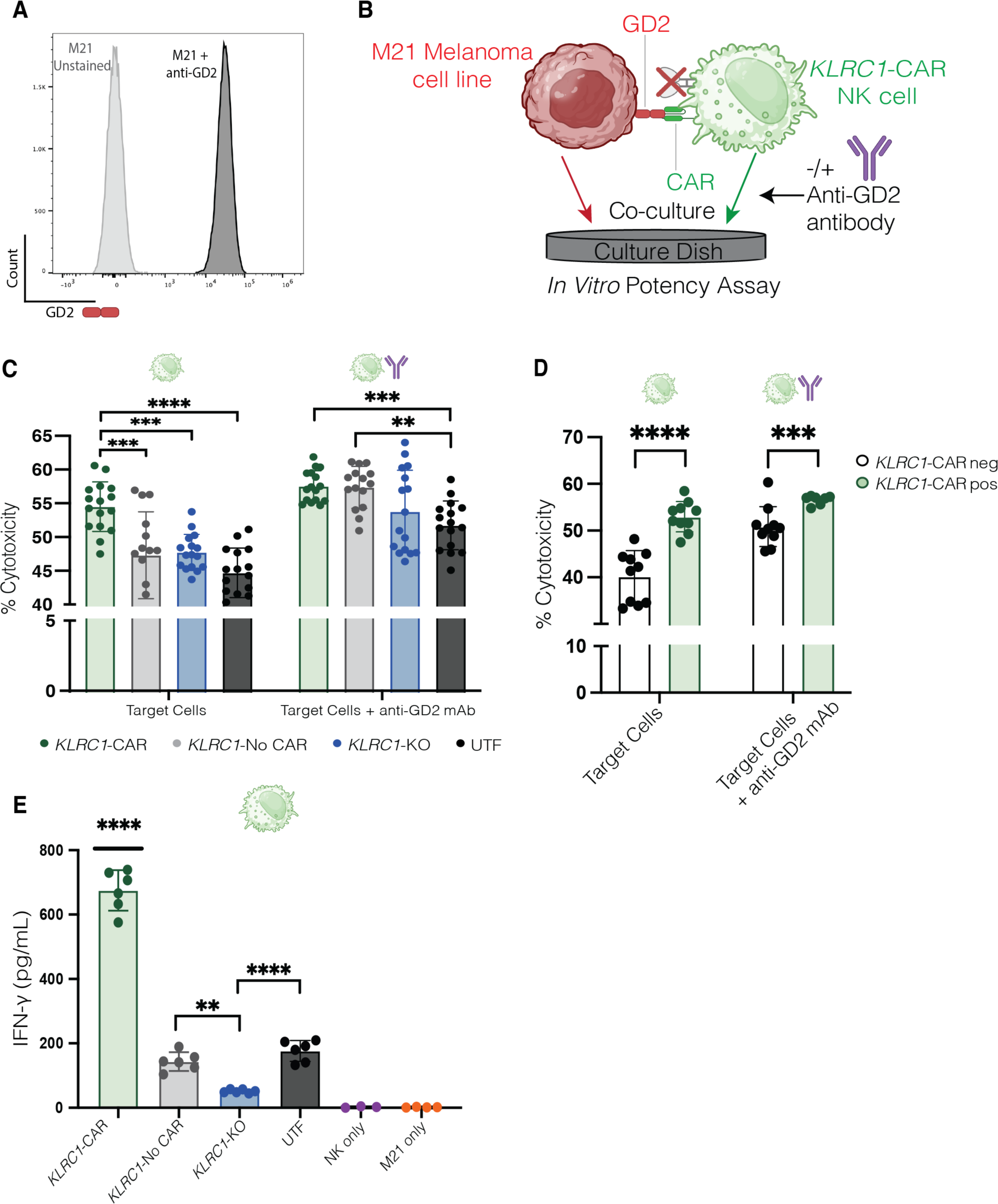
*KLRC1-*CAR cells display improved cytotoxicity against melanoma cells *in vitro*. **(A)** Flow cytometry histogram plot depicting GD2 expression on the surface of M21 melanoma cells. **(B)** Schematic depicting the *in vitro* potency assay used to assess NK cell cytotoxicity. **(C)** Following FACS, NK cells were co-cultured with M21 cells at a 5:1 E:T ratio for 24 hours. For ADCC conditions, post-FACS NK cells were co-cultured with M21 cells in the presence of 500 ng/mL hu14.18K322A anti-GD2 monoclonal antibody. The non-ADCC conditions are indicated with a single green NK cell icon above the graph, while the ADCC conditions are indicated by cell and antibody icons. Cytotoxicity was measured by quantifying secreted LDH with the CyQuant LDH Cytotoxicity Assay kit. Data is shown as the mean (SD) across three donors, with 5-6 replicates per donor (n = 3). **(D)** Cytotoxicity was measured by the same LDH release assay as in C, but for CAR^+^ and CAR^−^ fractions within the same sample. Data is shown as the mean (SD) across three donors, with 5-6 replicates per donor (n = 3). **(E)** NK cell secreted IFNγ was measured using an ELISA on media supernatant taken from the 24-hour co-culture with M21 cells. Data is shown as mean (SD) for one donor with 6 replicates (n = 1). Statistical significance was calculated using an ordinary two-way ANOVA, with the Dunnett’s multiple comparison test (C), and the Sidak’s multiple comparison post-test (D), and a one-way ANOVA with the Tukey’s multiple comparison post-test (E). *, P ≤ 0.05; **, P ≤ 0.01; ***, P ≤ 0.001; ****, P ≤ 0.0001; ns, P ≥ 0.05. LDH, lactate dehydrogenase; ADCC, antibody dependent cellular cytotoxicity; CAR, chimeric antigen receptor; ANOVA, analysis of variance.

To further verify that the increase in cytotoxicity is due to the CAR, we isolated CAR^+^ and CAR^−^ cell fractions within a sample and measured NK cytotoxicity. The CAR^+^ fractions exhibited 13% greater cytotoxicity than their CAR^−^ counterparts (**Figure 5D**). Again, this difference between the two populations was completely erased upon addition of the anti-GD2 monoclonal antibody under ADCC conditions.

Analysis of media supernatant indicated that the *KLRC1-*CAR NK cells secreted the greatest IFNγ upon antigen exposure (**Figure 5E**). Though there were no differences in the cytotoxicity between the *KLRC1-*No CAR and *KLRC1-*KO groups, we observed greater IFNγ production by the *KLRC1-* No CAR cells, indicating some antigen-independent activation of NK cells. The *KLRC1*-KO cells produced the least amount of IFNγ, even when compared to their UTF counterparts. Robust IFNγ secretion by the *KLRC1*-CAR NK cells reflects the high levels of cytotoxicity against melanoma cells.

### *KLRC1*-CAR NK cells overcome HLA-E based inhibition

To interrogate the combinatorial effect of the NKG2A knock-out and CAR insertion, we assessed the potency of the *KLRC1-*CAR NK cells against HLA-E expressing M21 cells. Following transduction with an HLA-E vector containing an HLA-G leader peptide,^10^ 44% of M21 cells expressed HLA-E on the cell surface, with expression increasing to 100% after a 24-hour treatment with 100 ng/mL of IFNγ (**Figure 6A**). Pure populations of NKG2A^−^, CAR^+^, or mCherry^+^ cells were isolated using FACS and co-cultured with IFNγ-treated M21-HLA-E cells at various effector-target ratios for 24 hours alongside donor-matched UTF controls **(Figures 6B, 6C)**. At the lowest ratio (0.1:1), there were no differences in NK-mediated cytotoxicity between the groups. When cultured at the 0.5:1 ratio, *KLRC1-*CAR NK cells displayed superior cytotoxicity compared to *KLRC1-*No CAR, *KLRC1-*KO, and UTF cells. At the highest ratio, *KLRC1-*CAR NK cells continued to exhibit improved cytotoxicity when compared to *KLRC1-*No CAR and UTF cells by 9% and 27% respectively, but no significant differences were observed relative to the *KLRC1-*KO NK cells. Again, in ADCC conditions, the differences in cytotoxicity were nullified between the edited groups for the 0.1:1 and 1:1 effector-target ratios, mirroring the trends seen against the wild-type M21 cells **(Figure 6D)**. Aside from a minor difference between the *KLRC1-*CAR and *KLRC1-* No CAR cells, the same results were observed at the 0.5:1 effector-target ratio in ADCC promoting conditions.

**Figure 6.**
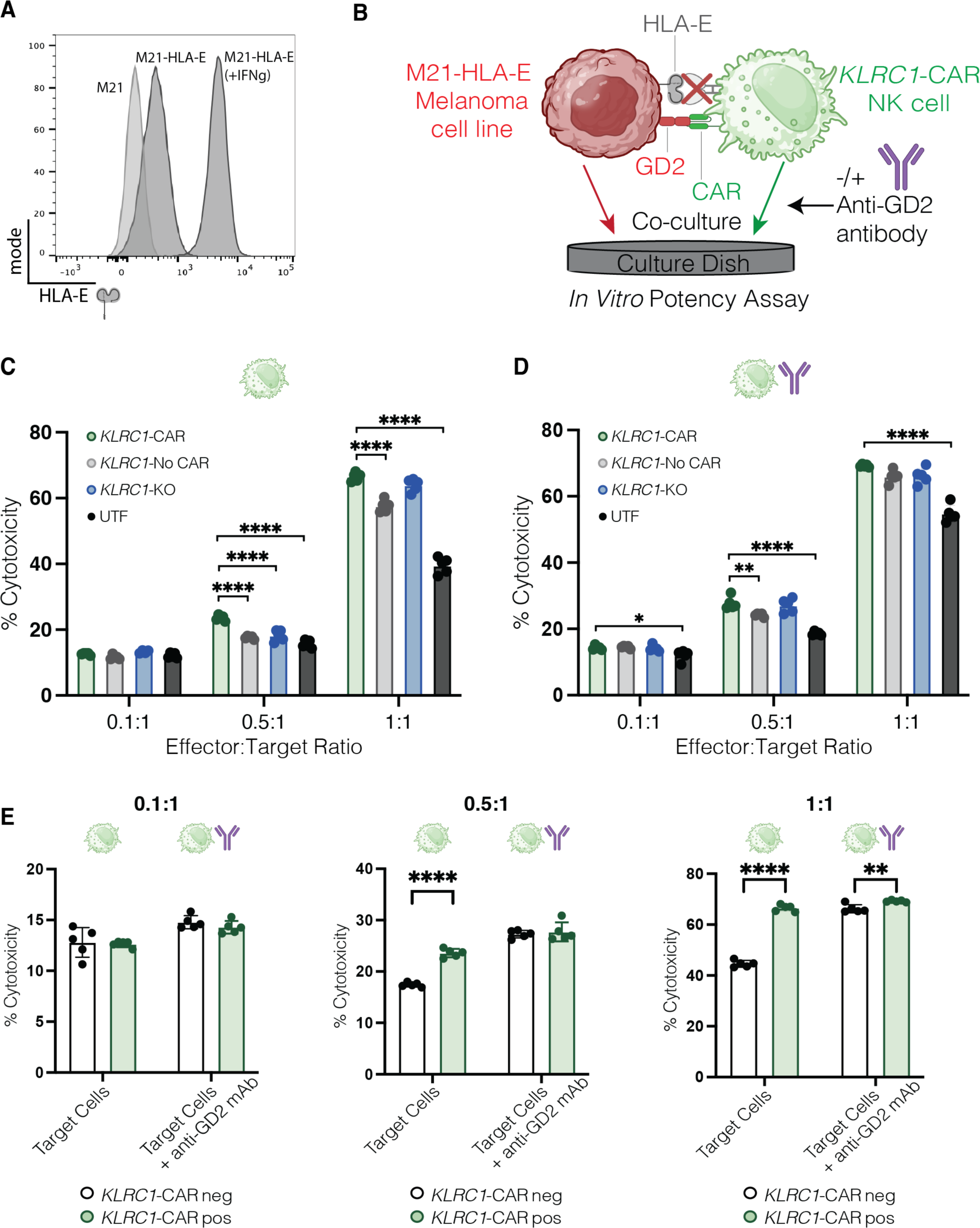
Overcoming the NKG2A-HLA-E based checkpoint. **(A)** Flow cytometry histogram plot showing HLA-E expression on wild-type M21 cells, HLA-E transduced M21 cells, and HLA-E transduced M21 cells treated with 100 ng/mL of IFNγ for 24 hours. **(B)** Schematic depicting the *in vitro* potency assay used to assess NK cell cytotoxicity against the M21-HLA-E cells. **(C)** After FACS, NK cells were co-cultured with M21-HLA-E cells at 0.1:1, 0.5:1, and 1:1 effector-target ratios for 24 hours. Cytotoxicity was measured by quantifying secreted LDH with the CyQuant LDH Cytotoxicity Assay kit. Data is shown as the mean (SD) of five replicates for one donor (n = 1). **(D)** Sorted NK cells were co-cultured with M21-HLA-E cells at the same three ratios as in C with 500 ng/mL of hu14.18K322A anti-GD2 monoclonal antibody to promote ADCC. Data is shown as the mean (SD) of five replicates for one donor (n = 1). **(E)** *KLRC1-*CAR NK samples were sorted into CAR^−^ and CAR^+^ fractions and were cultured with M21-HLA-E cells under the same conditions as in C and D. The secreted LDH was used to quantify NK cytotoxicity after 24 hours. Data is shown as the mean (SD) of five replicates for one donor (n = 1). Statistical significance was calculated using an ordinary one-way ANOVA, with the Dunnett’s multiple comparison test (C and D), and a two-way ANOVA with the Sidak’s multiple comparison post-test (E). *, P ≤ 0.05; **, P ≤ 0.01; ***, P ≤ 0.001; ****, P ≤ 0.0001; ns, P ≥ 0.05. LDH, lactate dehydrogenase; ADCC, antibody dependent cellular cytotoxicity; CAR, chimeric antigen receptor; ANOVA, analysis of variance.

Finally, we confirmed that the improvement in NK potency against HLA-E expressing cancer cells was still driven by CAR activity by analyzing the cytotoxicity of CAR^+^ and CAR^−^ fractions from within the same sample **(Figure 6E)**. Similar to earlier outcomes, the CAR^+^ fractions displayed improved cytotoxicity at 0.5:1 and 1:1 effector-target ratios by 7% and 22% respectively, with the differences minimized under ADCC conditions. At the lowest effector-target ratio, no differences in NK cytotoxicity were observed.

## Discussion

NK cells often fail to control solid tumors due to the immunosuppressive conditions of the tumor microenvironment, which can include the expression of ligands (e.g., HLA-E) that engage inhibitory receptors on NK cells like NKG2A.^35^ Though NK cells have previously been genetically modified to successfully improve cytotoxicity and persistence using viral vectors, transposons, mRNA transfection, and CRISPR+AAV technologies, these methods elicit concerns of safety, lack the machinery for stable editing, or have knock-in limitations on transgene size.^19,20,36–39^ Here, we demonstrate an optimized CRISPR-Cas9-enabled approach that generates stable and precisely edited primary NK cells without the use of viral vectors. Our data show that the CRISPR platform can be leveraged for efficient knock-out of the inhibitory NKG2A receptor and virus-free knock-in of an anti-GD2 CAR. We illustrate a high knock-in efficiency within primary NK cells for large transgenes (∼3.1 kb) with little to no off-target editing. Moreover, we also validate that *KLRC1*-CAR NK cells are viable, proliferative and capable of inducing tumor lysis and cytokine secretion *in vitro.* Taken together, our work establishes that *ex vivo* expanded NK cells from peripheral blood can be effectively modified to express a functional CAR, while also deleting an immune checkpoint molecule using CRISPR genome editing.

Despite nominating many off-target sites for the sgRNA from the highly sensitive CHANGE-seq assay, we saw minimal indel formation across the off-target sites. One explanation is that the RNP complex is only present in the NK cells transiently, resulting in lower exposure to nuclease activity when compared to the *in vitro* assay conditions used in CHANGE-Seq. Furthermore, M3814 may inhibit NHEJ events in cells that experienced an off-target double-stranded break resulting in either cell death or precise HDR. Notably, these findings are similar to that of the recently-approved exagamglogene autotemcel (exa-cel, Casgevy^TM^) Cas9-edited cell therapy, in which none of the nominated sites were seen upon validation in the cell product.^40,41^ Importantly, no off-target integration was observed, indicating that the endogenous *KLRC1* promoter is the sole driver of CAR expression of our promoter-less CAR transgene.

Our RNP-based knock-out strategy yielded high numbers of *KLRC1^−/−^*cells that were viable and proliferative, at efficiencies higher than those previously reported.^12,13,21^ Bexte et al. reported a knock-out efficiency of 43.5%, while Donald et al. reported an efficiency of 41% for sgRNAs targeting the third exon. Interestingly, Bexte et al. reported a lower indel frequency for the sgRNA used in this study (∼58-80%) in primary T cells. However, protein expression was not reported in that study, which makes it difficult to conclude how the editing translates to knock-out efficiencies in their system. In contrast, Huang et al., reported high indel frequencies (>80%) for the sgRNA used in this study, but showed less than 25% NKG2A knock-out. Though Donald et al. delivers a greater amount of nuclease, their system uses the Cas12a enzyme, highlighting another difference between the studies.

The dsDNA template strategy used in this study for transgene integration has been utilized in primary NK cells before, but for smaller sequences. The studies from the Lin and Marson groups report knock-in efficiencies ranging from 3.09% - 7.5% for a dsDNA GFP template (∼1.5 kb) across several loci.^21–23^ One study by Kath et. al reports knock-in outcomes of less than 10% for a truncated CAR of approximately 1.4 kb in size.^24^ Despite using transgenes that are larger than those published previously, this study showed similar knock-in rates with our system (without HDR enhancers), suggesting that templates of particular size ranges (i.e., 1.5 kb to 3 kb) may produce similar editing outcomes. Since HDR is more likely to occur within cells in the S- or -G2 phases,^33^ the best knock-in percentages occurred when NK cells were expanded with K562-mb15-41BBL feeder cells, IL-2, and IL-15 for the first four days. As the K562 cells are eliminated from the culture beyond day 4, the NK cells may enter a less activated state even with cytokine stimulation, which could explain the gradual decrease in editing on days 5-7. Two major differences between this study and those of the Lin and Marson groups is the length and type of activation prior to nucleofection. Regardless of whether cytokines, CD2/CD335 coated beads, or K562 feeder cells are used to activate NK cells, for two or four days, this study observes comparable knock-in rates across all three systems. Collectively, these findings suggest that *ex vivo* expanded NK cells are most amenable to genome engineering 2-4 days after isolation/thaw, with day 4 being an optimal time point if feeder cells are employed to ensure a pure, feeder-free NK population.

We found using an NHEJ inhibitor and fine-tuning of the pulse code in the nucleofector to be pivotal in improving our knock-in outcomes. The DNA-PK inhibitor, M3814, improved HDR outcomes by two-fold in our system. Interestingly, these results contrast with that of the Lin group where other HDR enhancers and DNA-PK inhibitors had no effect on editing efficiencies within NK cells.^21^ Despite being in the same class of drugs, M3814 and KU00060648 elicit a different effect on cell viability and editing, highlighting the need for further research on the use of pharmacological enhancers for this cell type. Our increase in editing is further compounded by using a stronger pulse code (EH-100) that likely increases the permeabilization of the cells, allowing for better cargo delivery without affecting cell viability. Surprisingly, the EH-115 program resulted in the lowest percent of mCherry^+^ cells at 13.8%, which is in contrast to the results reported by the Lin group.^21^ This discrepancy may be due to the fact that our system uses a linear, dsDNA template for transgene insertion instead of a plasmid. The DN-100 program resulted in knock-in efficiencies of 15%, which is approximately double what was published by the Lin group for the same dsDNA template strategy.^21^

NK cells are known to be challenging to genetically engineer due to their poor recovery post-editing.^21^ Their innate lineage equips them with pattern recognition receptors (PRR) and Toll-Like receptors (TLR) that can recognize cytosolic viral and dsDNA molecules, thus increasing dsDNA-associated toxicities.^15,16,42^ We observe slower proliferation of the *KLRC1*-CAR NK and *KLRC1*-No CAR NK cells in the first five days following electroporation, compared to the *KLRC1*-KO and UTF NK cells. The co-delivery of a dsDNA template along with the RNP complex is likely toxic to the cells and may cause activation of cGAS-STING, AIM2, or Type I-IFN signaling, ultimately suppressing cell proliferation and inducing apoptosis. Given that our templates for PCR for dsDNA template production are transformed and grown in *E. coli,* and purified in-house, it is also possible that our templates contain trace amounts of endotoxins that may further exacerbate the dsDNA-associated toxicity. Despite these challenges, our study demonstrates that this toxicity can be overcome through careful testing of parameters with highly purified reagents.

Our work provides compelling evidence that the CRISPR platform can be utilized to generate functional CAR NK cells using a primary cell source. When exposed to both wild-type and HLA-E expressing M21 cells, *KLRC1-*CAR NK cells display greater target lysis and IFNγ secretion compared to all other groups. In the HLA-E expressing model, the difference between *KLRC1-* CAR and UTF NK cells is further widened even when cultured at a lower effector-target ratio (9% for 5:1 E:T versus 27% for 1:1 E:T), underscoring the importance of disrupting the NKG2A-HLA-E checkpoint axis. Our data also suggests that NKG2A knock-out and CAR expression have an additive effect on NK potency against solid tumors. However, in ADCC conditions, we observe no differences in killing between the *KLRC1*-CAR, *KLRC1-*No CAR, and *KLRC1*-KO cells, highlighting the impact of CAR engagement and signaling in improving NK cytotoxicity. The increase in cytotoxicity of all groups except the *KLRC1-*CAR NK cells in ADCC conditions also indicates that the CAR NK cells already display maximal cytotoxic capacity on their own, while the *KLRC1-*No CAR, *KLRC1-*KO, and UTF NK cells require the addition of a monoclonal antibody to achieve comparable activity.

Clinical translation of this cell therapeutic strategy is likely viable with further research along two principal directions. First, additional analysis of *KLRC1-*CAR NK cells within the M21-HLA-E system, such as cytokine profiles and activation signatures, would further characterize the potency profile of the cell product. Second, the use of feeder cells could be eliminated through the use of feeder-free NK expansion protocols.^21,43,44^ Finally, the potency of these cells upon cryopreservation and thawing will also need to be evaluated to truly enable an off-the-shelf strategy. Altogether, we demonstrated the feasibility of using the CRISPR system to genetically modify primary NK cells, and of driving CAR expression and activity via the *KLRC1* locus. This CRISPR-based platform can enable virus-free manufacturing of off-the-shelf, genetically engineered NK cells to remove inhibitory immune checkpoints while simultaneously inserting CARs with the potential for enhanced potency against solid tumors.

## Materials and Methods

### Cell Lines

M21 human melanoma cells were a gift from Dr. Paul Sondel (University of Wisconsin-Madison), and K562-mb15-41BBL cells were a gift from Dr. Dario Campana (National University of Singapore). Both cell lines were maintained in RPMI 1640 (Gibco, Waltham, MA) supplemented with 10% fetal bovine serum (FBS) (WiCell, Madison, WI), 1% penicillin-streptomycin (Gibco, Waltham, MA), and 1% L-glutamine (Gibco, Waltham, MA). NK-92 cells (ATCC, Manassas, VA) were cultured in X-VIVO 10 (Lonza, Basel, Switzerland) supplemented with 5% FBS (WiCell, Madison, WI), 5% horse serum (ThermoFisher, Waltham, MA), 199.82 μM myo-inositol (Sigma Aldrich, St. Louis, MO), and 19.94 μΜ folic acid (Sigma Aldrich, St. Louis, MO). The M21-HLA-E transduced cell line was maintained in the same medium with the addition of 600 μg/mL of G418 Geneticin. Cell authentication was performed using short tandem repeat analysis (Idexx BioAnalytics, Westbrook, Maine) and per ATCC guidelines using cell morphology, growth curves, and *Mycoplasma* testing within 6 months with the MycoStrip Mycoplasma Detection Kit (Invitrogen, Waltham, MA). Cell lines were maintained in culture at 37°C in 5% CO_2_.

### Transduction of M21 cell line

The human HLA-E gene insert (**Table S1**), incorporating the HLA-G leader peptide, and *HindIII* and *EcoRI* restriction sites, was generated synthetically by IDT (IDT, Newark, NJ). The pcDNA™3.1(+) plasmid (ThermoFisher, Waltham, MA) and the synthesized HLA-E gene were digested separately using *HindIII* and *EcoRI* enzymes (ThermoFisher, Waltham, MA) following the manufacturer’s instructions. The digested products (backbone of pcDNA^™^3.1 and HLA-E gene product) were ligated using the T4 DNA ligase enzyme (ThermoFisher, Waltham, MA) according to the manufacturer’s instructions. The ligated product was transformed into JM109 competent *E*. *coli*. (Promega, Madison, WI) using a heat shock protocol. To confirm successful transformation of clones, DNA sequencing was performed using primers specific to the pcDNA3.1 CMV (forward) and BGH (reverse) sequences.

24 hours before transfection, M21 cells (1×10^6^ cells per well) were plated in a 6 well plate, with cells reaching 80-90% confluency at the time of transfection. Cell transfection was performed using the TransIT-Lenti transfection reagent (Mirus bio, Madison, WI) per the manufacturer’s instructions. After 24 hours of transfection, the cells were treated with 600 μg/mL of G418 Geneticin (ThermoFisher, Waltham, MA) to select for transfected cells. Monoclonal cell populations were generated through a serial dilution technique, and flow cytometry was used to select the clone with the highest HLA-E expression.

### Isolation and expansion of NK cells

Whole blood was collected from the median cubital vein of healthy donors using 10mL lithium heparin-coated vacutainers (158 USP units, BD Biosciences, Franklin Lakes, NJ). Peripheral blood (PB) NK cells were isolated from whole blood using a density based negative selection kit according to the manufacturer’s instructions (RosetteSep Human NK Enrichment Cocktail, STEMCELL Technologies, Vancouver, Canada). Isolated PB-NK cells were expanded with irradiated K562-mb15-41BBL feeder cells (1-3 days before NK cell isolation, 100 Gy) along with 50 IU/mL IL-2 (Peprotech, Cranbury, NJ) and 2 ng/mL IL-15 (Peprotech, Cranbury, NJ) at a ratio of 1:2 in NK MACS media (Miltenyi Biotec, Gaithersburg, MD) supplemented with 5% human AB serum (Millipore Sigma, Burlington, MA) and 1% MACS supplement (Miltenyi Biotec, Gaithersburg, MD). Cells were cultured until day 4, after which they were nucleofected for experiments. After day 5, unedited and nucleofected NK cells were cultured in complete NK MACS medium (Miltenyi Biotec, Gaithersburg, MD) with 100 IU/mL IL-2 and 2 ng/mL IL-15. Cells were maintained at a density of 1e6 cells/mL and were passaged every two days. Feeder cells were added to support NK expansion on days 0 and 5.

### *sp*Cas9 RNP complexing

Purified sNLS-SpCas9-sNLS Cas9 protein (10 mg/mL, Aldevron, South Fargo, ND) and sgRNA (60 pmoles, IDT, Newark, NJ) with the crRNA and tracrRNA pre-complexed were aliquoted into individual PCR tubes for each nucleofection replicate. Sterile-filtered 15–50 kDa poly(L-glutamic acid) (PGA) (Sigma Aldrich, St. Louis, MO) was added to the sgRNA at 0.8:1 volume ratio as previously described.^22^ 50 pmoles of Cas9 was added to the sgRNA + PGA mixture for a final molar ratio of 1:1.2 sgRNA:Cas9. The complexes were incubated at 37° C for 15 minutes prior to use.

### Preparation of dsDNA templates

PCR amplicons of the CAR and mCherry transgenes were generated as described previously.^34^ PCR products were pooled into 600 μL and were purified using a singular round of Solid Phase Reversible Immobilization (SPRI) cleanup using AMPure XP beads (Beckman Coulter, Brea, CA) according to the manufacturer’s instructions, at a 1:1 ratio. Bead incubation and elution times were increased to 5 minutes and 15 minutes, respectively. Each pooled sample was eluted in 30 μL of nuclease-free water (IDT, Newark, NJ), which were then combined to form one sample. 3M pH5.5 Sodium Acetate (NaOAc) (Sigma Aldrich, St. Louis, MO) was added to the pooled sample at 0.1x of the total volume. The sample was ethanol precipitated by adding 2.5x of 100% ethanol, followed by a 30-minute incubation at −80° C. The precipitate was purified using an ethanol wash (70-80%) and eluted in 20-30 μL of nuclease free water. DNA template concentration was measured and adjusted as previously described.^34^ Primers and DNA sequences are listed in Table S1.

### Nucleofection

NK cells were nucleofected on day 4, unless otherwise described for optimization experiments. Post-nucleofection recovery medium consisting of Complete NK MACS medium (Miltenyi Biotec, Gaithersburg, MD) with 100 IU/mL IL-2 and 2 ng/mL IL-15 +/− 0.6uM M3814 (Selleck Chemicals, Houston, TX) was prepared, with M3814 added at 1.2X the working concentration to account for future dilution. For M3814 related optimization experiments, multiple aliquots of post-recovery medium were prepared with a corresponding concentration of M3814. Next, 150uL of M3814 recovery medium was plated into individual wells of a 96-well round bottom plate for each replicate and set aside in the incubator. PB-NK cells were harvested and counted using the Countess II FL Automated Cell Counter (ThermoFisher, Waltham, MA) with 0.4% Trypan Blue viability stain (ThermoFisher, Waltham, MA). 5e5 cells were aliquoted into individual 1.5 mL Eppendorf tubes for each nucleofection replicate. While the RNP complexes incubated, the cells were pelleted at 100g for 10 minutes, and 3μg (1.5uL) of dsDNA templates were aliquoted into PCR tubes for each replicate. Following incubation, the RNP complexes were added to the dsDNA templates and were allowed to incubate for 3-5 minutes at room temperature. Cell pellets were resuspended in P3 buffer with 22% supplement (Lonza, Basel, Switzerland), and combined with the RNP+dsDNA mixture, for a total volume of 24 μL. Samples were added to the 24 μL reaction cuvettes (Lonza, Basel, Switzerland) and were nucleofected using the EH-100 program (Lonza Amaxa 4D Nucleofector, Basel, Switzerland), unless specified otherwise for optimization experiments. Immediately after nucleofection, 100 μL of pre-warmed recovery medium (no M3814) was added to each of the cuvette wells, and samples were rested at 37° C for 15 minutes. Finally, samples were transferred from the cuvette to the 96-well plate containing pre-warmed recovery medium with M3814. After 24 hours, nucleofected NK cells were combined with irradiated K562-mb15-41BBL (100 Gy) at a 1:1 ratio in complete NK MACS medium with 100 IU/mL IL-2 and 2 ng/mL IL-15.

### Flow cytometry and cell sorting

1e5 cells were plated per condition and stained in 96-well round bottom plates. Cells were washed with 1X PBS, centrifuged at 1200g for 1 minute, and stained with 0.1 μL/mL Zombie Aqua viability stain (Biolegend, San Diego, CA) for 20 minutes. Next, cells were washed twice with 2% FACS buffer (1X PBS and 2% BSA) centrifuged and stained with human TruStain FcX (Biolegend, San Diego, CA) for 20 minutes. Cells were washed twice more with 2% FACS buffer, pelleted, and stained with NKG2A (Clone S19004C, Biolegend, San Diego, CA) and anti-14G2a (Clone 1A7, National Cancer Institute, Biological Resources Branch, Rockville MD) antibodies for detection of NKG2A and CAR scFv, respectively. For phenotyping experiments, NK cells were also stained with CD56 (Clone HCD56, Biolegend, San Diego, CA) and CD16 (Clone 3G8, Biolegend, San Diego, CA). Flow cytometry was performed on an Attune NxT flow cytometer (ThermoFisher, Waltham, MA), and data was analyzed using FlowJo.

For sorting, NK cells were either stained for NKG2A or GD2 CAR and were sorted using a 130 μm nozzle on a FACS Aria II sorter (BD Biosciences, Franklin Lakes, NJ). Sorted cells were recovered in Complete NK MACS medium (Miltenyi Biotec, Gaithersburg, MD) with 100 IU/mL IL-2 and 2 ng/mL IL-15 for four days prior to cytotoxicity assays. NK cells were assayed by flow cytometry or sorted seven days after nucleofection (Day 11). For flow analysis of the M21 melanoma cell line, cells were detached using 0.05% Trypsin (ThermoFisher, Waltham, MA) for 5 minutes prior to filtering with a 70 μm strainer, counting and plating for staining. M21 cells were stained as described above, but using an anti-GD2 antibody (Clone 14G2a, Biolegend, San Diego, CA).

### LDH release cytotoxicity assay

First, 1e4 M21 cells were plated per well in a 96-well, flat bottom plate in culture medium with or without 100 ng/mL of IFNγ. 24 hours later, sorted NK cells were added to the well at the corresponding effector: target ratios in complete NK MACS medium supplemented with 100 IU/mL IL-2 and 2 ng/mL IL-15 for a total volume of 100 μL. For ADCC conditions, 500 ng/mL of anti-GD2 hu14.18K322A (a gift from Paul Sondel) was added to each of the wells. After 24 hours of co-culture, the plate was centrifuged at 300g for 5 minutes to pellet the cells. 50 μL of supernatant was harvested and assayed for secreted LDH using the CyQuant LDH Cytotoxicity Assay kit according to the manufacturer’s instructions (ThermoFisher, Waltham, MA). The remaining supernatant was collected and frozen at −80° C for subsequent ELISA.

### IFNγ ELISA

Quantification of secreted IFNγ was carried out using an IFNγ sandwich ELISA kit (ThermoFisher, Waltham, MA) according to the manufacturer’s instructions on media supernatants (undiluted) taken from the 24-hour LDH co-culture assay.

### In-Out PCR

Genomic DNA was extracted from 1e6 cells as described previously.^34^ One primer was designed to bind upstream of the left homology arm or downstream of the right homology arm, depending on the orientation of transgene integration. The second primer was designed to bind within the scFv region of the CAR, or within the protein coding region of the mCherry transgene. PCRs were carried out using the Q5 Hot Start 2x Master Mix (NEB, Ipswich, MA) according to the manufacturer’s protocol for 30 cycles. Primer sequences are listed in Table S1.

### On-target editing analysis via Next Generation Sequencing

Genomic DNA was extracted and indel formation was analyzed via next generation sequencing (Illumina, San Diego, CA) as previously described.^34^ Briefly, edited amplicons were amplified and purified using genomic PCR and SPRI cleanup, respectively. A second PCR was performed with indexing primers to attach unique dual identifiers (Illumina, San Diego, CA) to the amplicons. After a second SPRI cleanup, samples were combined and run on the Illumina MiniSeq per the manufacturer’s instructions and analyzed using the CRISPR RGEN software. Primer sequences are listed in Table S1.

### Off-target nomination via genome wide CHANGE-seq

The Gentra Puregene kit (Qiagen, Germantown, MD) was used to extract genomic DNA from unedited primary NK cells per the manufacturer’s instructions. Genomic analysis was performed by CHANGE-Seq as previously described.^45^ In short, genomic DNA underwent tagmentation with a custom Tn5-transposome, and was gap repaired with HiFi HotStart Uracil+ Ready Mix (Kapa Biosystems, Wilmington, MA). USER enzyme (NEB, Ipswich, MA) and T4 polyncucleotide was then used to treat the gap-repaired DNA. DNA circularization was performed using T4 DNA ligase (NEB, Ipswich, MA), followed by treatment with a mixture of exonucleases to remove residual linear DNA. *In vitro* cleavage reactions were performed on the circularized DNA using *Sp*Cas9 protein (NEB, Ipswich, MA) and *KLRC1* sgRNA. Cleaved products were incubated with proteinase K (NEB, Ipswich, MA), A-tailed, ligated with a hairpin adapter (NEB, Ipswich, MA), and treated with USER enzyme (NEB, Ipswich, MA). Products were amplified by PCR with the Kapa HiFi polymerase (Kapa Biosystems, Wilmington, MA), and libraries were quantified by qPCR (Kapa Biosystems, Wilmington, MA). Libraries were sequenced with 150 bp paired-end reads on the Illumina NextSeq2000 instrument. CHANGE-seq data analyses were performed as described previously.^45^

### Off-target analysis at nominated sites via Next Generation Sequencing

Indel frequency at off-target sites was assayed by the rhAmpSeq system (IDT, Newark, NJ). Genomic DNA was extracted from *KLRC1*-CAR NK cells. Pools of forward and reverse primers were designed using IDT’s rhAmpSeq design tool to amplify the on-target and top 12 off-target sites detected by CHANGE-Seq. PCRs and library preparation were performed according to IDT’s instructions. Pooled libraries were diluted and sequenced on the Illumina MiniSeq instrument with 150 bp paired-end reads. Data was analyzed using IDT’s rhAmpSeq analysis tool.

### Statistical Analysis

All analyses were performed using GraphPad Prism (V10.1.0). For experiments with two groups, a two-tailed *t* test was used. For optimization experiments with more than two groups, a one-way ANOVA was used, with the Prism recommended post-test. For editing and cytotoxicity experiments with multiple donors, a one-way or two-way ANOVA was used, followed by the Prism recommended post-test. Post-tests are listed in the figure legend for each experiment. Error bars represent mean (SD). ns, p ≥ 0.05; *, p ≤ 0.05; **, p ≤ 0.01; ***, p ≤ 0.001; ****, p ≤ 0.0001.

## Data Availability Statement

The datasets used and/or analyzed during the current study are available from the corresponding author on reasonable request.

## Acknowledgements

We acknowledge funding from the NIH/NCI R01 CA278051, St. Baldrick’s Foundation Empowering Immunotherapies for Childhood Cancer (EPICC) research grant, the NSF Engineering Research Center (ERC) for Cell Manufacturing Technologies (CMaT) NSF-EEC 1648035, Hyundai Hope on Wheels, the Grainger Institute for Engineering at UW-Madison (CMC and KS), the MACC Fund (CMC), and NIH R35 GM119644-01 (KS), and NIH U01AI176471 (SQT). We also acknowledge the University of Wisconsin Carbone Cancer Center Flow Cytometry Laboratory, the Small Animal Imaging and Radiotherapy Facility, and the TSB-BioBank, supported by NCI P30 CA014520 and 1S10RR025483-01 (BD FACS Aria II BSL-2 Cell Sorter), for use of its facilities and services. We thank the NCI Biological Resources Branch for 1A7 anti-14G2a antibody for detection of CAR scFV, Paul Sondel for the hu14.18K322A monoclonal antibody and M21 human melanoma cells, and Dario Campana for the K562-mb15-41BBL cells. We also thank Matthew Forsberg and the members of the Saha lab for helpful input on experimental design and manuscript preparation. The contents of this article do not necessarily reflect the views or policies of the Department of Health and Human Services, nor does mention of trade names, commercial products, or organizations imply endorsement by the US Government.

## Author Contributions

KSh and BER designed the sgRNA sequences, and KSh designed the mCherry and CAR sequences. KSh and BER designed, performed, and analyzed knock-out experiments and NGS studies in NK-92 cells. KSh collected whole blood, isolated NK cells, and established the cell expansion and gene editing protocol. KSh designed, performed, and analyzed all gene editing optimization studies and co-culture experiments. IZH assisted with DNA template production, cell culture, and extraction of genomic DNA. LS generated the M21-HLA-E line and helped perform characterization experiments. VK performed and analyzed the CHANGE-Seq assay, and KSh performed PCRs, carried out NGS experiments, and ran analysis for subsequent off-target experiments. KSh wrote the manuscript with input from all authors. KSa, CMC, and SQT supervised the research. All authors read and approved the final manuscript.

## Ethics Approval

All human studies were conducted using a University of Wisconsin-Madison IRB-approved protocol (2018-0103, approved 08/31/23)

## Supplemental Tables

**Table S1.** Nucleotide sequences of DNA templates, gRNAs, and primers used in this study.

**Table S2.** Complete list of potential off-target sites detected by CHANGE-Seq for the *KLRC1* gRNA used in this study.

