## Supplemental Figures for "Virus-free CRISPR knock-in of a chimeric antigen receptor into *KLRC1* generates potent GD2-specific natural killer cells"

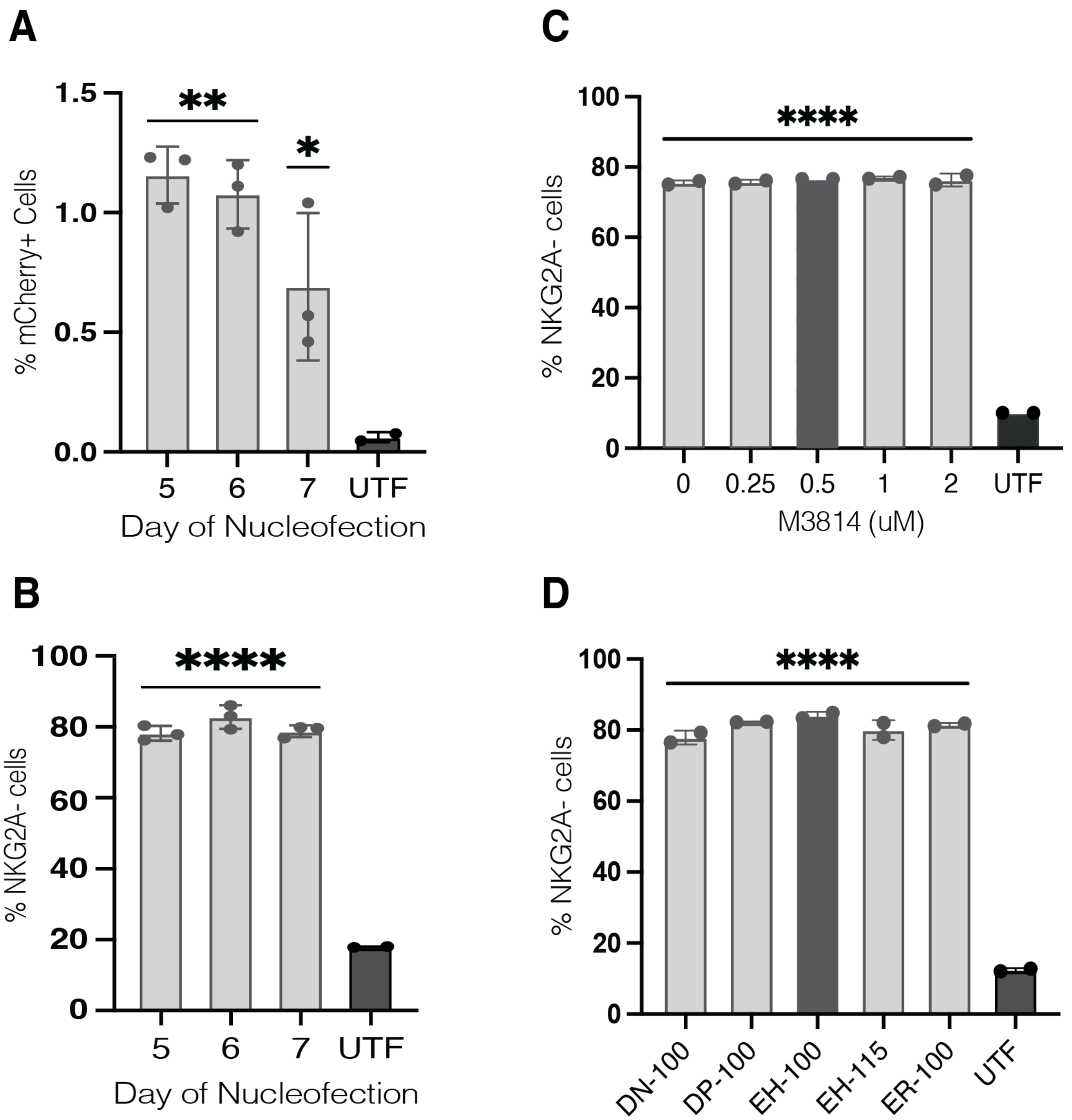


**Figure S1. Temporal optimization of transgene delivery and the effect of M3814 and pulse programs on NKG2A knock-out efficiency, related to Figure 2. (A and B)** NK cells were nucleofected on days 5, 6, or 7 of expansion. Percent of mCherry$^{+}$ cells and NKG2A$^{-}$ cells were measured by flow cytometry. Data is shown for one donor with three replicates (n = 1). **(C)** Percent of NKG2A$^{-}$ cells after 24-hour incubation with M3814 at varying concentrations, assayed by flow cytometry one week after nucleofection. Data is shown for two replicates for one donor (n = 1). **(D)** Percent of NKG2A$^{-}$ cells after nucleofection with varying pulse programs, measured by flow cytometry one week after nucleofection. Data is shown for two replicates for one donor (n = 1). Significance was calculated using an ordinary one-way ANOVA, with the Dunnett's multiple comparisons post-test. *, P $\leq$ 0.05; **, P $\leq$ 0.01; ****, P $\leq$ 0.0001. ANOVA, analysis of variance.

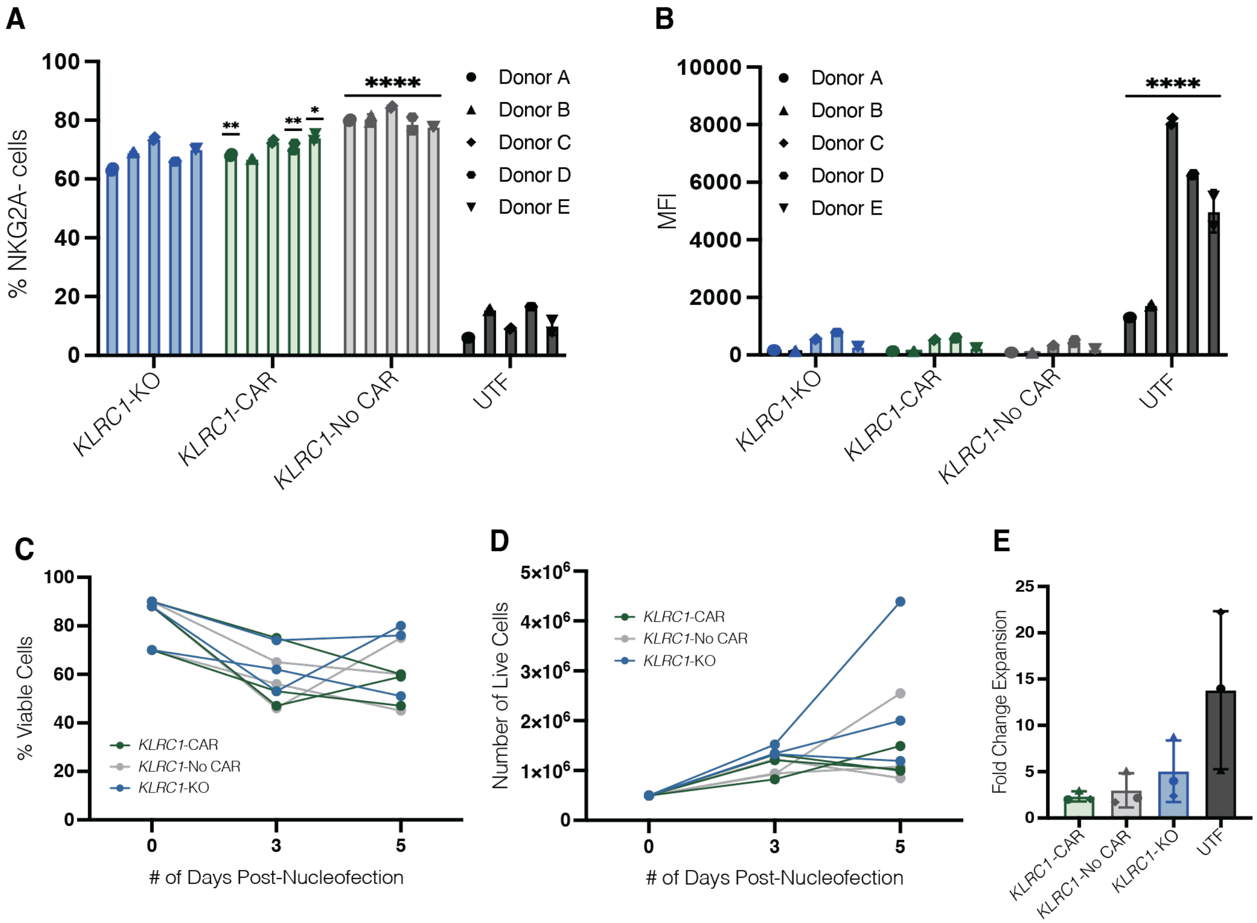


**Figure S2. Impact of transgene knock-in on NKG2A knock-out efficiency and cell expansion, related to Figure 3. (A and B)** NK cells were nucleofected on day 4 of expansion with either a Cas9 RNP complex, or a Cas9 RNP complex with a dsDNA template encoding the CAR or mCherry gene. The percent of NKG2A$^{-}$ cells was measured by flow cytometry one week after nucleofection, along with NKG2A MFI levels on the cell surface. Data is shown as an average of two replicates across five donors (n = 5). Significance tests were performed by comparing % NKG2A expression on *KLRC1*-KO cells versus *KLRC1*-CAR, and *KLRC1*-No CAR NK cells. **(C and D)** Cell viability and cell expansion of *KLRC1*-CAR, *KLRC1*-No CAR, *KLRC1*-KO NK cells are shown for five days after nucleofection. Data is an average of two replicates for each of the three donors (n = 3). **(E)** Calculated fold change expansion of *KLRC1*-CAR, *KLRC1*-No CAR, and *KLRC1*-KO NK cells from D is shown in comparison to donor-matched UTF cells, with each donor indicated by a different shape (circle, triangle, diamond). An ordinary one-way ANOVA was used to test for statistical significance. The Dunnett's multiple comparisons test (A and B) and Tukey's multiple comparisons test (E) were used as the post-test. *, $P \leq 0.05$; **, $P \leq 0.01$; ****, $P \leq 0.0001$; ns, $P \geq 0.05$. RNP, ribonucleoprotein; dsDNA, double stranded DNA; CAR, chimeric antigen receptor; MFI, mean fluorescence intensity; ANOVA, analysis of variance.

**A**

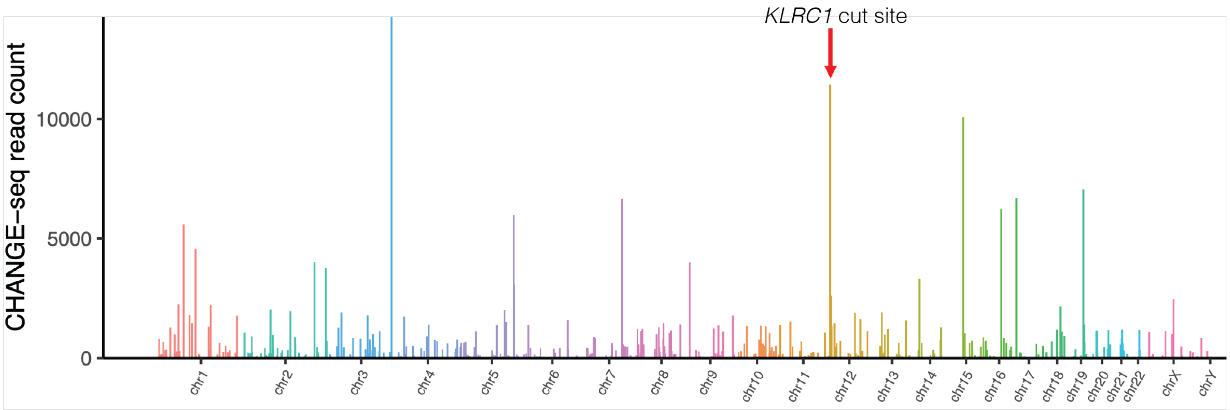


**Figure S3. Genome wide off-target analysis of the Cas9 editor via CHANGE-Seq, related to Figure 4. (A)** Manhattan plot describing off-target activity of the KLRC1 gRNA across the genome, as detected by CHANGE-Seq. The intended on-target site is indicated by the red arrow. X-axis describes the chromosomal location, and the y-axis describes the CHANGE-Seq read counts (n = 1).
