## Supplemental Table 1 for "Virus-free CRISPR knock-in of a chimeric antigen receptor into *KLRC1* generates potent GD2-specific natural killer cells"

| DNA Sequences (5' -> 3') | |
|---|---|
| **gRNA1 (PAM)** | GGTCTGAGTAGATTACTCCT(TGG) |
| **gRNA2 (PAM)** | TGAACAGGAAATAACCTATG(CGG) |
| **gRNA3 (PAM)** | TCTTATCTTAATGGCCTCTG(TGG) |
| **Left homology arm** | AACTTCTTTATTCCTTAGTAATTTATTGCTTCTCATTGCCCCAGCAATAATATTTTGTCAAATGCAGAAAAtttatcttttttttttgagtcggagtctcactctgtcgcccaggctggagtgcagtggcacaatctccgctcactgcaacctctgccgcccatgttcaagagattctcctgcctcagcctcccgagtagctaggactacaggcgcctgacatcatgcccgactactttttgtatttttagtagagacggagtttcaccgtgttagccaggatggtctcgatctcctgacctcgtgatcggcatgcctcggcctcccaaagtgctgggattacaggcgtgagccaccgcgcccggccTAAAAATCTTTTTTTAAAACAAATATTCATAAGAAACGTGTTTAGGCTTGAAGAAAATCAGAGAAAGAACTTTAGATTATTTAATGCAAAATGAGCTCCAATACTCGTTCTCCACCTCACCCTTTTAATTGCACTAGGGAATCCTGTATATAAACCATTTATTAACTTCTTAACTACTGTTATTATAGAGTACAGTCCCTGACATCACACACTGCAGAGATGGATAACCAAGG |
| **Right homology arm** | AGTAATCTACTCAGACCTGAATCTGCCCCCAAACCCAAAGAGGCAGCAACGAAAACCTAAAGGCAATAAAAACTCCATTTTAGCAACTGAACAGGAAATAACCTATGCGGAATTAAACCTTCAAAAAGCTTCTCAGGATTTTCAAGGGAATGACAAAACCTATCACTGCAAAGGTAAAGCATTTAAAAGATCCTCAATATAACAGTCTAGGATGTGCAGCTTGGGGTACAGGAATGTGGGGAAAGAGAAGGGAGTGCTCATATATCTTCTATTTGCAAAGATCAGAATTCCAAGTTGAGATATGCTATTTCAATGTAAAGTATGAAGACTGATTGAACTCATTGTTGAAGTTTGTAGTCTTTGTCAAATAATTCATGGAGCATTATTTTTCCTGAAAATTCAATGGTATATTATTCTGAGAAAAAGATTACAATGGGAGATGAGGGTTTGGGGTCCAAGTTTCTCTGTATGATTCCTGTGCATTCAGGTTCTCTTGTCTGTGAATCTTCTAAACGACTGTATCCACCTCTCCTTTCGCACTGTTCCCATTTCTCTCCCTGCAGATTTACCATCAGCTCCAGAGAAGCTCATTGTTGGGAT |
| **CAR** | AggcagcggagagggcagaggaagtcttctaacatgcggtgacgtggaggagaatcccggccctATGGAGTTTGGGCTGAGCTGGCTTTTTCTTGTGGCTATTTTAAAAGGTGTCCAGTGCTCTAGAGATATTTTGCTGACCCAAACTCCACTCTCCCTGCCTGTCAGTCTTGGAGATCAAGCCTCCATCTCTTGCAGATCTAGTCAGAGTCTTGTACACCGTAATGGAAACACCTATTTACATTGGTACCTGCAGAAGCCAGGCCAGTCTCCAAAGCTCCTGATTCACAAAGTTTCCAACCGATTTTCTGGGGTCCCAGACAGGTTCAGTGGCAGTGGATCAGGGACAGATTTCACACTCAAGATCAGCAGAGTGGAGGCTGAGGATCTGGGAGTTTATTTCTGTTCTCAAAGTACACATGTTCCTCCGCTCACGTTCGGTGCTGGGACCAAGCTGGAGCTGAAACGGGCTGATGCTGCACCAACTGTATCCATCTTCCCAGGCTCGGGCGGTGGTGGGTCGGGTGGCGAGGTGAAGCTTCAGCAGTCTGGACCTAGCCTGGTGGAGCCTGGCGCTTCAGTGATGATATCCTGCAAGGCTTCTGGTTCCTCATTCACTGGCTACAACATGAACTGGGTGAGGCAGAACATTGGAAAGAGCCTTGAATGGATTGGAGCTATTGATCCTTACTATGGTGGAACTAGCTACAACCAGAAGTTCAAGGGCAGGGCCACATTGACTGTAGACAAATCGTCCAGCACAGCCTACATGCACCTCAAGAGCCTGACATCTGAGGACTCTGCAGTCTATTACTGTGTAAGCGGAATGGAGTACTGGGGTCAAGGAACCTCAGTCACCGTCTCCTCAGCCAAAACGACACCCCCATCAGTCTATGGAAGGGTCACCGTCTCTTCAGCGGAGCCCAAATCTTGTGACAAAACTCACACATGCCCACCGTGCCCGGATCCCAAATTTTGGGTGCTGGTGGTGGTTGGTGGAGTCCTGGCTTGCTATAGCTTGCTAGTAACAGTGGCCTTTATTATTTTCTGGGTGAGGAGTAAGAGGAGCAGGCTCCTGCACAGTGACTACATGAACATGACTCCCCGCCGCCCCGGGCCCACCCGCAAGCATTACCAGCCCTATGCCCCACCACGCGACTTCGCAGCCTATCGCTCCAAACGGGGCAGAAAGAAACTCCTGTATATATTCAAACAACCATTTATGAGACCAGTACAAACTACTCAAGAGGAAGATGGCTGTAGCTGCCGATTTCCAGAAGAAGAAGAAGGAGGATGTGAACTGAGAGTGAAGTTCAGCAGGAGCGCAGACGCCCCCGCGTACCAGCAGGGCCAGAACCAGCTCTATAACGAGCTCAATCTAGGACGAAGAGAGGAGTACGATGTTTTGGACAAGAGACGTGGCCGGGACCCTGAGATGGGGGGAAAGCCGAGAAGGAAGAACCCTCAGGAAGGCCTGTACAATGAACTGCAGAAAGATAAGATGGCGGAGGCCTACAGTGAGATTGGGATGAAAGGCGAGCGCCGGAGGGGCAAGGGGCACGATGGCCTTTACCAGGGTCTCAGTACAGCCACCAAGGACACCTACGACGCCCTTCACATGCAGGCCCTGCCCCCTcgcTAATTCACTCCTCAGGTGCAGGCTGCCTATCAGAAGGTGGTGGCTGGTGTGGCCAATGCCCTGGCTCACAAATACCACTGAGATCTTTTTCCCTCTGCCAAAAATTATGGGGACATCATGAAGCCCCTTGAGCATCTGACTTCTGGCTAATAAAGGAAATTTATTTTCATTGCAATAGTGTGTTGGAATTTTTTGTGTCTCTCACTCGGAAGGACATATGGGAGGGCAAATCATTTAAAACATCAGAATGAGTATTTGGTTTAGAGTTTGGCAACATATGCCCATATGCTGGCTGCCATGAACAAAGGTTGGCTATAAAGAGGTCATCAGTATATGAAACAGCCCCCTGCTGTCCATTCCTTATTCCATAGAAAAGCCTTGACTTGAGGTTAGATTTTTTTTATATTTTGTTTTGTGTTATTTTTTTCTTTAACATCCCTAAAATTTTCCTTACATGTTTTACTAGCCAGATTTTTCCTCCTCTCCTGACTACTCCCAGTCATAGCTGTCCCTCTTCTCTTATGGAGATCCCTCGACCTGCAGCCCAAGCTTGGCGTAATCATGGTCATAGCTGT |
| **mCherry** | AggcagcggagagggcagaggaagtcttctaacatgcggtgacgtggaggagaatcccggccctATGCCTGAACCCTCTAAGTCTGCTCCAGCCCCTAAAAAGGGTTCTAAGAAGGCTATCACTAAGGCGCAGAAGAAGGATGGTAAGAAGCGTAAGCGCAGCCGCAAGGAGAGCTATTCTATCTATGTGTACAAGGTTCTGAAGCAGGTCCACCCCGACACCGGCATCTCATCCAAGGCCATGGGGATCATGAACTCCTTCGTCAACGACATCTTCGAGCGCATCGCGGGCGAGGCTTCTCGCCTGGCTCACTACAATAAGCGCTCGACCATCACCTCCAGGGAGATTCAGACGGCTGTGCGCCTGCTGCTGCCTGGGGAGCTGGCTAAGCATGCTGTGTCCGAGGGCACTAAGGCAGTTACCAAGTACACTAGCTCTAAGGATCCACCGGTCGCCACCATGGTGAGCAAGGGCGAGGAGGATAACATGGCCATCATCAAGGAGTTCATGCGCTTCAAGGTGCACATGGAGGGCTCCGTGAACGGCCACGAGTTCGAGATCGAGGGCGAGGGCGAGGGCCGCCCCTACGAGGGCACCCAGACCGCCAAGCTGAAGGTGACCAAGGGTGGCCCCCTGCCCTTCGCCTGGGACATCCTGTCCCCTCAGTTCATGTACGGCTCCAAGGCCTACGTGAAGCACCCCGCCGACATCCCCGACTACTTGAAGCTGTCCTTCCCCGAGGGCTTCAAGTGGGAGCGCGTGATGAACTTCGAGGACGGCGGCGTGGTGACCGTGACCCAGGACTCCTCCCTGCAGGACGGCGAGTTCATCTACAAGGTGAAGCTGCGCGGCACCAACTTCCCCTCCGACGGCCCCGTAATGCAGAAGAAGACCATGGGCTGGGAGGCCTCCTCCGAGCGGATGTACCCCGAGGACGGCGCCCTGAAGGGCGAGATCAAGCAGAGGCTGAAGCTGAAGGACGGCGGCCACTACGACGCTGAGGTCAAGACCACCTACAAGGCCAAGAAGCCCGTGCAGCTGCCCGGCGCCTACAACGTCAACATCAAGTTGGACATCACCTCCCACAACGAGGACTACACCATCGTGGAACAGTACGAACGCGCCGAGGGCCGCCACTCCACCGGCGGCATGGACGAGCTGTACAAGTAAttcactcctcaggtgcaggctgcctatcagaaggtggtggctggtgtggccaatgccctggctcacaaataccactgagatctttttccctctgccaaaaattatggggacatcatgaagccccttgagcatctgacttctggctaataaaggaaatttattttcattgcaatagtgtgttggaattttttgtgtctctcactcggaaggacatatgggagggcaaatcatttaaaacatcagaatgagtatttggtttagagtttggcaacatatgcccatatgctggctgccatgaacaaaggttggctataaagaggtcatcagtatatgaaacagccccctgctgtccattccttattccatagaaaagccttgacttgaggttagattttttttatattttgttttgtgttatttttttctttaacatccctaaaattttccttacatgttttactagccagatttttcctcctctcctgactactcccagtcatagctgtccctcttctcttatggagatccctcgacctgcagcccaagcttggcgtaatcatggtcatagctgt |
| **HLA-E** | AAGCTTATGGTGGTCATGGCGCCCCGAACCCTCTTCCTGCTGCTCTCGGGGGCCCTGACCCTGACCGAGACCTGGGCGGGCTCCCACTCCTTGAAGTATTTCCACACTTCCGTGTCCCGGCCCGGCCGCGGGGAGCCCCGCTTCATCTCTGTGGGCTACGTGGACGACACCCAGTTCGTGCGCTTCGACAACGACGCCCGCGAGTCCGAGGATGGTGCCGCGGGCGCCGTGGATGGAGCAGGAGGGGTCAGAGTATTGGGACCGGGAGACACGGAGCGCCAGGGACACCGCACAGATTTTCCGAGTGAACCTGCGGACGCTGCGCGGCTACTACAATCAGAGCGAGGCCGGGTCTCACACCCTGCAGTGGATGCATGGCTGCGAGCTGGGGCCCGACAGGCGCTTCCTCCGCGGGTATGAACAGTTCGCCTACGACGGCAAGGATTATCTCACCCTGAATGAGGACCTGCGCTCCTGGACCGCGGTGGACACGGCGGCTCAGATCTCCGAGCAAAAGTCAAATGATGCCTCTGAGGCGGAGCACCAGAGAGCCTACCTGGAAGACACATGCGTGGAGTGGCTCCACAAATACCTGGAGAAGGGGAAGGAGACGCTGCTTCACCTGGAGCCCCCAAAGACACACGTGACTCACCACCCCATCTCTGACCATGAGGCCACCCTGAGGTGCTGGGCCCTGGGCTTCTACCCTGCGGAGATCACACTGACCTGGCAGCAGGATGGGGAGGGCCATACCCAGGACACGGAGCTCGTGGAGACCAGGCCTGCAGGGGATGGAACCTTCCAGAAGTGGGCAGCTGTGGTGGTGCCTTCTGGAGAGGAGCAGAGATACACGTGCCATGTGCAGCATGAGGGGCTACCCGAGCCCGTCACCCTGAGATGGAAGCCGGCTTCCCAGCCCACCATCCCCATCGTGGGCATCATTGCTGGCCTGGTTCTCCTTGGATCTGTGGTCTCTGGAGCTGTGGTTGCTGCTGTGATATGGAGGAAGAAGAGCTCAGGTGGAAAAGGAGGGAGCTACTCTAAGGCTGAGTGGAGCGACAGTGCCCAGGGGTCTGAGTCTCACAGCTTGTAAGAATTC |

| Primers for dsDNA template preparation (5' -> 3') | |
|---|---|
| Forward | cggagtctcactctgtcgccca |
| Reverse | TGGACCCCAAACCCTCATCTCCC |

| On-Target Next Generation Sequencing Primers | |
|---|---|
| Forward (5' -> 3') | TCCCTGACATCACACACTGC |
| Reverse (5' -> 3') | TGCCTTTAGGTTTTCGTTGC |

**Template Integration PCR Primer Sequences**

| On Target | | | | |
|---|---|---|---|---|
| Locus | Transgene | Forward (5' -> 3') | Reverse (5' -> 3') | Expected size (bp) |
| KLRC1 | CAR | TGTGCAGACCACATAGTCTTAACCA | aggaaccagaagccttgcagga | 1349 |
| KLRC1 | mCherry | TGTGCAGACCACATAGTCTTAACCA | TGCCTTAGTGCCCTCGGACACA | 1156 |

| Off-Target (Forward orientation) | | | | | |
|---|---|---|---|---|---|
| Off Target Site | Chromosome | Locus | Forward (5' -> 3') | Reverse (5' -> 3') | Expected size (bp) |
| 1 | 3 | RTP4 | AGATTTGGGGGAAGCACTGAAGTC | GGGGGTGTCGTTTTGGCTGAGG | 1831 |
| 2 | 15 | CDAN1 | CTGGCTTTTGTGTTACCTGCTTGAC | GGGGGTGTCGTTTTGGCTGAGG | 1632 |
| 3 | 19 | ZNF146 | TATGCAGGGAGCAGGGAGGATATAG | GGGGGTGTCGTTTTGGCTGAGG | 1457 |
| 4 | 17 | RABEP1 | tgggaggctgaggcaggagaac | ACAGTTGGTGCAGCATCAGCCC | 1611 |
| 5 | 7 | ST7-OT3 | AATATTGGCAGGAAGGCAGGACAG | ACAGTTGGTGCAGCATCAGCCC | 1931 |
| 6 | 16 | SALL1 | CAAAGGGACCCTGCCCACC | GGGGGTGTCGTTTTGGCTGAGG | 1810 |
| 7 | 5 | HAND1 | CACGGGCAATCCCCGCA | GGGGGTGTCGTTTTGGCTGAGG | 1738 |
| 8 | 1 | LRRIQ3 | TGGTCATGCTTGGAAGACCCTTG | GGGGGTGTCGTTTTGGCTGAGG | 1776 |
| 9 | 1 | AKNAD1 | TCCAAGACTGCTTTTCCTGTGGG | GGGGGTGTCGTTTTGGCTGAGG | 1922 |
| 10 | 2 | NRP2 | GCGCAGAAGAGTGTAAGAAGGTCA | GGGGGTGTCGTTTTGGCTGAGG | 1805 |
| 11 | 9 | LOC105375972 | CTCCATGGCAGCATTTCTCATTTGG | GGGGGTGTCGTTTTGGCTGAGG | 2206 |
| 12 | 2 | ASB1 | caccccatgcacacacctccac | GGGGGTGTCGTTTTGGCTGAGG | 1677 |

| Off-Target (Reverse orientation) | | | | | |
|---|---|---|---|---|---|
| Off Target Site | Chromosome | Locus | Forward (5' -> 3') | Reverse (5' -> 3') | Expected size (bp) |
| 1 | 3 | RTP4 | GGGGGTGTCGTTTTGGCTGAGG | TGCCAAGACAAGTCAGGTTTGATGA | 1723 |
| 2 | 15 | CDAN1 | GGGGGTGTCGTTTTGGCTGAGG | GCCCCATTCTCCGTACCTGC | 1829 |
| 3 | 19 | ZNF146 | GGGGGTGTCGTTTTGGCTGAGG | TAGTGCTTGACCCATGCAGAACC | 1759 |
| 4 | 17 | RABEP1 | GGGGGTGTCGTTTTGGCTGAGG | CTGGCTGAAGTGGGCGTCCTTG | 2011 |
| 5 | 7 | ST7-OT3 | GGGGGTGTCGTTTTGGCTGAGG | GAATTCTGTCTGGCAGGTCCCC | 1980 |
| 6 | 16 | SALL1 | GGGGGTGTCGTTTTGGCTGAGG | TTCTTCCTCTCTGTCCCCTGAGTC | 1540 |
| 7 | 5 | HAND1 | GGGGGTGTCGTTTTGGCTGAGG | AATCAGGGGCTACCGTTGCG | 1696 |
| 8 | 1 | LRRIQ3 | GGGGGTGTCGTTTTGGCTGAGG | TGCAGAACCCATGACCAATCTCTG | 1625 |
| 9 | 1 | AKNAD1 | GGGGGTGTCGTTTTGGCTGAGG | AACAGCCCATCTCAAGGCACC | 2362 |
| 10 | 2 | NRP2 | GGGGGTGTCGTTTTGGCTGAGG | GCCTCTTCCCGTTTGAGGTTTCT | 1830 |
| 11 | 9 | LOC105375972 | GGGGGTGTCGTTTTGGCTGAGG | CCCTCCCCTCTGGGATAGAGTTTT | 1761 |
| 12 | 2 | ASB1 | GGGGGTGTCGTTTTGGCTGAGG | CTGACATTTCACCAAGGGAGCCA | 2066 |
