## Supplemental Table 2 for "Virus-free CRISPR knock-in of a chimeric antigen receptor into *KLRC1* generates potent GD2-specific natural killer cells"

| Chr | Start | End | Strand | Peak Score | Focus Ratio/Region Size | Annotation | Detailed Annotation | Distance to TSS | Nearest PromoterID | Entrez ID | Nearest Unigene | Nearest Refseq | Nearest Ensembl | Gene Name | Gene Type |
|---|---|---|---|---|---|---|---|---|---|---|---|---|---|---|---|
| chr3 | 187358041 | 187358063 | + | 14288 | NA | Intergenic | Intergenic | -10328 | NM_022147 | 64108 | Hs.43388 | NM_022147 | ENSG00000136514 | RTP4 | protein-coding |
| chr12 | 10451126 | 10451148 | + | 11436 | NA | exon (NM_213658, exon 3 of 8) | exon (NM_213658, exon 3 of 8) | 2243 | NM_007328 | 3821 | Hs.512576 | NM_002259 | ENSG00000134545 | KLRC1 | protein-coding |
| chr15 | 42665152 | 42665174 | - | 10080 | NA | intron (NM_020759, intron 13 of 32) | intron (NM_020759, intron 13 of 32) | 71965 | NM_138477 | 146059 | Hs.599232 | NM_138477 | ENSG00000140326 | CDAN1 | protein-coding |
| chr19 | 36212768 | 36212790 | + | 7058 | NA | intron (NM_001042474, intron 1 of 4) | intron (NM_001042474, intron 1 of 4) | -1823 | NM_007145 | 7705 | Hs.643436 | NM_007145 | ENSG00000167635 | ZNF146 | protein-coding |
| chr17 | 5322688 | 5322710 | + | 6694 | NA | intron (NM_001083585, intron 2 of 16) | L1MA4\|LINE\|L1 | 40436 | NM_004703 | 9135 | Hs.584784 | NM_004703 | ENSG00000029725 | RABEP1 | protein-coding |
| chr7 | 117240145 | 117240167 | + | 6646 | NA | Intergenic | Intergenic | 57475 | NR_002332 | 93655 | Hs.368131 | NR_002332 | ENSG00000004866 | ST7-OT3 | ncRNA |
| chr16 | 51260180 | 51260202 | - | 6236 | NA | intron (NR_039672, intron 6 of 6) | intron (NR_039672, intron 6 of 6) | -108919 | NM_002968 | 6299 | Hs.135787 | NM_002968 | ENSG00000103449 | SALL1 | protein-coding |
| chr5 | 154483000 | 154483022 | + | 5978 | NA | Intergenic | Intergenic | -4747 | NM_004821 | 9421 | Hs.152531 | NM_004821 | ENSG00000113196 | HAND1 | protein-coding |
| chr1 | 73938555 | 73938577 | + | 5586 | NA | Intergenic | Intergenic | 259621 | NM_001105659 | 127255 | Hs.644625 | NM_145258 | ENSG00000162620 | LRRIQ3 | protein-coding |
| chr1 | 108835341 | 108835363 | - | 4568 | NA | intron (NM_152763, intron 7 of 15) | intron (NM_152763, intron 7 of 15) | 21752 | NM_152763 | 254268 | Hs.729441 | NM_152763 | ENSG00000162641 | AKNAD1 | protein-coding |
| chr2 | 205650340 | 205650361 | + | 3998 | NA | Intergenic | Intergenic | -32150 | NM_201267 | 8828 | Hs.471200 | NM_003872 | ENSG00000118257 | NRP2 | protein-coding |
| chr9 | 9535157 | 9535179 | - | 3990 | NA | intron (NM_002839, intron 8 of 45) | intron (NM_002839, intron 8 of 45) | -264224 | NR_135135 | 105375972 | Hs.738242 | NR_135135 | ENSG00000230920 | LOC105375972 | ncRNA |
| chr2 | 238487345 | 238487367 | + | 3770 | NA | Intergenic | Intergenic | 60428 | NM_001040445 | 51665 | Hs.516788 | NM_016114 | ENSG00000065802 | ASB1 | protein-coding |
| chr14 | 22389328 | 22389350 | - | 3320 | NA | intron (NR_148361, intron 2 of 4) | intron (NR_148361, intron 2 of 4) | 43414 | NR_148364 | 105370401 | Hs.435144 | NR_148361 | | LOC105370401 | ncRNA |
| chr5 | 155438094 | 155438116 | + | 3056 | NA | Intergenic | MLT1A0-int\|LTR\|ERVL-MaLR | 424350 | NM_001099293 | 285643 | Hs.567824 | NM_001099293 | ENSG00000226650 | KIF4B | protein-coding |
| chr12 | 12508435 | 12508457 | + | 2612 | NA | intron (NM_030640, intron 3 of 6) | intron (NM_030640, intron 3 of 6) | 54068 | NM_030640 | 80824 | Hs.536535 | NM_030640 | ENSG00000111266 | DUSP16 | protein-coding |
| chrX | 77858667 | 77858689 | + | 2458 | NA | intron (NM_032121, intron 3 of 9) | intron (NM_032121, intron 3 of 9) | 36890 | NM_032121 | 84061 | Hs.323562 | NM_032121 | ENSG00000102158 | MAGT1 | protein-coding |
| chr1 | 59058607 | 59058629 | + | 2248 | NA | intron (NR_110626, intron 2 of 2) | intron (NR_110626, intron 2 of 2) | 38142 | NR_110626 | 101926925 | | NR_110626 | | LINC01358 | ncRNA |
| chr1 | 152759401 | 152759422 | + | 2218 | NA | intron (NM_001025231, intron 1 of 1) | intron (NM_001025231, intron 1 of 1) | 1381 | NM_001025231 | 448834 | Hs.149386 | NM_001025231 | ENSG00000203786 | KPRP | protein-coding |
| chr18 | 50010494 | 50010515 | + | 2152 | NA | intron (NM_001080467, intron 4 of 39) | MIR3\|SINE\|MIR | 116059 | NR_036204 | 100422865 | | NR_036204 | ENSG00000283343 | MIR4320 | ncRNA |
| chr2 | 78158047 | 78158069 | + | 2014 | NA | intron (NR_110288, intron 1 of 3) | intron (NR_110288, intron 1 of 3) | 69328 | NR_110287 | 101927948 | Hs.639404 | NR_110287 | ENSG00000229494 | LOC101927948 | ncRNA |
| chr5 | 127654701 | 127654723 | + | 2002 | NA | intron (NM_001048252, intron 2 of 2) | intron (NM_001048252, intron 2 of 2) | 1697 | NM_001127385 | 613212 | Hs.66194 | NM_001048252 | ENSG00000205279 | CTXN3 | protein-coding |
| chr2 | 135351226 | 135351248 | - | 1966 | NA | intron (NM_001286568, intron 4 of 20) | intron (NM_001286568, intron 4 of 20) | 179493 | NM_001286569 | 84083 | Hs.658422 | NM_032143 | ENSG00000121988 | ZRANB3 | protein-coding |
| chr13 | 27458837 | 27458859 | + | 1906 | NA | Intergenic | Intergenic | -8246 | NM_152912 | 219402 | Hs.534582 | NM_152912 | ENSG00000122033 | MTIF3 | protein-coding |
| chr3 | 42058142 | 42058164 | + | 1902 | NA | Intergenic | Intergenic | -29047 | NR_146089 | 22906 | Hs.535711 | NM_014965 | ENSG00000182606 | TRAK1 | protein-coding |
| chr12 | 82600333 | 82600355 | + | 1894 | NA | Intergenic | Intergenic | -86811 | NM_001320322 | 160335 | Hs.577775 | NM_152588 | ENSG00000179104 | TMTC2 | protein-coding |
| chr1 | 91315548 | 91315570 | - | 1806 | NA | intron (NM_001017975, intron 28 of 38) | intron (NM_001017975, intron 28 of 38) | 89310 | NM_001017975 | 164045 | Hs.454818 | NM_001017975 | ENSG00000162669 | HFM1 | protein-coding |
| chr3 | 118188972 | 118188993 | + | 1788 | NA | Intergenic | Tigger3\|DNA\|TcMar-Tigger | -452786 | NR_135573 | 101926968 | Hs.518185 | NR_135573 | | LOC101926968 | ncRNA |
| chr9 | 134994009 | 134994031 | - | 1774 | NA | Intergenic | Intergenic | -76017 | NM_002003 | 2219 | Hs.440898 | NM_002003 | ENSG00000085265 | FCN1 | protein-coding |
| chr1 | 229184410 | 229184432 | + | 1766 | NA | Intergenic | Intergenic | -9620 | NR_039659 | 100616234 | | NR_039659 | ENSG00000283468 | MIR4454 | ncRNA |
| chr4 | 26056467 | 26056489 | - | 1728 | NA | Intergenic | LTR33A\|LTR\|ERVL | 142286 | NM_001145432 | 389203 | Hs.479386 | NM_001145432 | ENSG00000250317 | SMIM20 | protein-coding |
| chr12 | 98738716 | 98738738 | + | 1624 | NA | intron (NM_001352223, intron 10 of 10) | intron (NM_001352223, intron 10 of 10) | 93427 | NM_181868 | 317 | Hs.552567 | NM_001160 | ENSG00000120868 | APAF1 | protein-coding |
| chr6 | 129911823 | 129911845 | + | 1580 | NA | Intergenic | Intergenic | -50563 | NM_001010876 | 253582 | | NM_001010876 | ENSG00000203756 | TMEM244 | protein-coding |
| chr13 | 97555665 | 97555687 | - | 1568 | NA | Intergenic | L2c\|LINE\|L2 | 121455 | NM_021033 | 5911 | Hs.508480 | NM_021033 | ENSG00000125249 | RAP2A | protein-coding |
| chr11 | 30002747 | 30002769 | - | 1510 | NA | Intergenic | Intergenic | 14272 | NM_002233 | 3739 | Hs.592002 | NM_002233 | ENSG00000182255 | KCNA4 | protein-coding |
| chr5 | 132437251 | 132437273 | + | 1498 | NA | intron (NM_001207001, intron 2 of 3) | intron (NM_001207001, intron 2 of 3) | 26489 | NM_001207001 | | | | | | |
| chr1 | 98486462 | 98486484 | + | 1466 | NA | Intergenic | MER135\|DNA\|DNA | -175207 | NR_033716 | 51375 | Hs.197015 | NM_015976 | ENSG00000162627 | SNX7 | protein-coding |
| chr8 | 78572053 | 78572075 | + | 1464 | NA | intron (NM_181839, intron 1 of 2) | intron (NM_181839, intron 1 of 2) | -13561 | NR_125389 | 101927003 | Hs.614849 | NR_125389 | ENSG00000254266 | PKIA-AS1 | ncRNA |
| chr12 | 23220510 | 23220531 | - | 1458 | NA | Intergenic | Intergenic | 44884 | NR_120471 | 101928441 | Hs.129111 | NR_120471 | ENSG00000256321 | LOC101928441 | ncRNA |
| chr8 | 127158177 | 127158199 | + | 1416 | NA | Intergenic | Intergenic | 39439 | NR_120364 | 103021165 | Hs.573407 | NR_120364 | | CASC19 | ncRNA |
| chr19 | 38702872 | 38702894 | - | 1412 | NA | intron (NM_001322033, intron 3 of 20) | intron (NM_001322033, intron 3 of 20) | 41591 | NM_144691 | 147968 | Hs.731775 | NM_144691 | ENSG00000182472 | CAPN12 | protein-coding |
| chr4 | 97411318 | 97411340 | + | 1394 | NA | intron (NR_102713, intron 1 of 5) | intron (NR_102713, intron 1 of 5) | 44403 | NR_102713 | 101410545 | Hs.628520 | NR_102713 | ENSG00000251620 | STPG2-AS1 | ncRNA |
| chr11 | 298839 | 298860 | + | 1384 | NA | intron (NM_001025295, intron 1 of 1) | intron (NM_001025295, intron 1 of 1) | 677 | NM_001025295 | 387733 | Hs.443469 | NM_001025295 | ENSG00000206013 | IFITM5 | protein-coding |
| chr6 | 15718035 | 15718056 | + | 1380 | NA | Intergenic | L1M5\|LINE\|L1 | -54987 | NM_001271669 | 84062 | Hs.571148 | NM_032122 | ENSG00000047579 | DTNBP1 | protein-coding |
| chr5 | 105349315 | 105349336 | - | 1378 | NA | Intergenic | Intergenic | 249851 | NR_000039 | 9366 | Hs.158296 | NR_000039 | ENSG00000232159 | RAB9BP1 | pseudo |
| chr9 | 93178002 | 93178024 | + | 1372 | NA | Intergenic | Intergenic | -6917 | NM_006648 | 65268 | Hs.654856 | NM_006648 | ENSG00000165238 | WNK2 | protein-coding |
| chr10 | 78082323 | 78082345 | - | 1354 | NA | Intergenic | Intergenic | 48574 | NM_001142284 | 6229 | Hs.280130 | NM_001026 | ENSG00000138326 | RPS24 | protein-coding |
| chr12 | 14591814 | 14591836 | - | 1346 | NA | intron (NR_120465, intron 2 of 5) | intron (NR_120465, intron 2 of 5) | -23968 | NM_024829 | 79887 | Hs.131933 | NM_024829 | ENSG00000121316 | PLBD1 | protein-coding |
| chr10 | 91545437 | 91545459 | - | 1344 | NA | intron (NR_024467, intron 1 of 4) | intron (NR_024467, intron 1 of 4) | 66012 | NR_024467 | 100188947 | Hs.535293 | NR_024467 | | HECTD2-AS1 | ncRNA |
| chr10 | 37145525 | 37145547 | + | 1342 | NA | intron (NM_052997, intron 8 of 35) | intron (NM_052997, intron 8 of 35) | 19679 | NM_052997 | 91074 | Hs.373787 | NM_052997 | ENSG00000148513 | ANKRD30A | protein-coding |
| chr1 | 147111576 | 147111597 | + | 1316 | NA | intron (NR_103466, intron 2 of 5) | intron (NR_103466, intron 2 of 5) | 2760 | NR_103466 | 644861 | Hs.451804 | NR_103466 | | NBPF13P | pseudo |
| chr10 | 92412122 | 92412144 | + | 1300 | NA | Intergenic | MER21-int\|LTR\|ERVL | -6528 | NR_038243 | 100507674 | Hs.568772 | NR_038243 | | MARK2P9 | pseudo |
| chr8 | 64974377 | 64974399 | + | 1294 | NA | Intergenic | L1MA9\|LINE\|L1 | -175597 | NM_001324112 | 9420 | Hs.657330 | NM_004820 | ENSG00000172817 | CYP7B1 | protein-coding |
| chr14 | 85375203 | 85375225 | + | 1292 | NA | Intergenic | LTR16C\|LTR\|ERVL | -12061 | NR_135155 | 105370605 | | NR_135155 | ENSG00000258814 | LINC02329 | ncRNA |
| chr1 | 35193490 | 35193512 | - | 1282 | NA | promoter-TSS (NR_136702) | promoter-TSS (NR_136702) | -356 | NR_136702 | 6421 | Hs.355934 | NM_005066 | ENSG00000116560 | SFPQ | protein-coding |
| chr3 | 32958686 | 32958708 | - | 1270 | NA | Intergenic | Intergenic | 7123 | NM_005508 | 1233 | Hs.184926 | NM_005508 | ENSG00000183813 | CCR4 | protein-coding |
| chr9 | 79400521 | 79400543 | + | 1230 | NA | Intergenic | Intergenic | -171241 | NM_001351564 | 7091 | Hs.444213 | NM_007005 | ENSG00000106829 | TLE4 | protein-coding |
| chr8 | 16985841 | 16985863 | - | 1214 | NA | Intergenic | MLT1E1A-int\|LTR\|ERVL-MaLR | 16313 | NM_019851 | 26281 | Hs.199905 | NM_019851 | ENSG00000078579 | FGF20 | protein-coding |
| chr8 | 2528173 | 2528195 | - | 1212 | NA | Intergenic | Intergenic | 95181 | NR_125423 | 101927815 | Hs.571467 | NR_125423 | ENSG00000254319 | LOC101927815 | ncRNA |
| chr13 | 44974928 | 44974950 | - | 1206 | NA | intron (NM_012345, intron 5 of 9) | intron (NM_012345, intron 5 of 9) | 14539 | NM_012345 | 26747 | Hs.525006 | NM_012345 | ENSG00000083635 | NUFIP1 | protein-coding |
| chr21 | 26117898 | 26117920 | - | 1188 | NA | intron (NM_001136131, intron 1 of 15) | intron (NM_001136131, intron 1 of 15) | 22481 | NM_001136016 | 351 | Hs.434980 | NM_000484 | ENSG00000142192 | APP | protein-coding |
| chr18 | 39366350 | 39366372 | + | 1186 | NA | intron (NR_024391, intron 3 of 3) | intron (NR_024391, intron 3 of 3) | 255814 | NR_030628 | 100126323 | | NR_030628 | ENSG00000283615 | MIR924 | ncRNA |
| chr22 | 29706370 | 29706392 | - | 1174 | NA | Intergenic | L2b\|LINE\|L2 | -13974 | NM_182527 | 164633 | Hs.643608 | NM_182527 | ENSG00000100314 | CABP7 | protein-coding |
| chr20 | 50417001 | 50417023 | - | 1168 | NA | Intergenic | MIR\|SINE\|MIR | -93309 | NM_001278618 | 5770 | Hs.417549 | NM_002827 | ENSG00000196396 | PTPN1 | protein-coding |
| chr8 | 99789561 | 99789583 | - | 1152 | NA | intron (NM_017890, intron 43 of 61) | L1ME4a\|LINE\|L1 | 104442 | NM_004374 | 1345 | Hs.351875 | NM_004374 | ENSG00000164919 | COX6C | protein-coding |
| chr20 | 17751444 | 17751466 | + | 1146 | NA | Intergenic | L1ME3A\|LINE\|L1 | 51507 | NM_001014977 | 140836 | Hs.115366 | NM_178477 | ENSG00000125888 | BANF2 | protein-coding |
| chrX | 54134362 | 54134384 | - | 1124 | NA | intron (NM_001300788, intron 7 of 13) | intron (NM_001300788, intron 7 of 13) | 48885 | NM_017848 | 54954 | Hs.86045 | NM_017848 | ENSG00000184083 | FAM120C | protein-coding |
| chr20 | 15593102 | 15593123 | - | 1122 | NA | intron (NM_001033087, intron 4 of 13) | intron (NM_001033087, intron 4 of 13) | 330202 | NR_130925 | 613266 | Hs.309149 | NR_130924 | | LOC613266 | ncRNA |
| chr12 | 118969844 | 118969866 | + | 1110 | NA | Intergenic | Intergenic | -11640 | NM_194286 | 84530 | Hs.744964 | NM_194286 | ENSG00000139767 | SRRM4 | protein-coding |
| chr3 | 153065647 | 153065669 | + | 1110 | NA | Intergenic | L1ME3A\|LINE\|L1 | -96554 | NM_002886 | 5912 | Hs.98643 | NM_002886 | ENSG00000181467 | RAP2B | protein-coding |
| chr8 | 13215843 | 13215865 | - | 1108 | NA | intron (NM_182643, intron 5 of 17) | intron (NM_182643, intron 5 of 17) | 60694 | NM_001316668 | 10395 | Hs.134296 | NM_006094 | ENSG00000164741 | DLC1 | protein-coding |
| chr5 | 44324957 | 44324979 | + | 1102 | NA | intron (NM_004465, intron 1 of 2) | intron (NM_004465, intron 1 of 2) | 63714 | NM_004465 | 2255 | Hs.248049 | NM_004465 | ENSG00000070193 | FGF10 | protein-coding |
| chr9 | 106603299 | 106603321 | + | 1096 | NA | Intergenic | Intergenic | 2382 | NR_107048 | 102465995 | | NR_107048 | ENSG00000277889 | MIR8081 | ncRNA |
| chr18 | 54299814 | 54299836 | - | 1094 | NA | intron (NR_147257, intron 9 of 9) | L1MEc\|LINE\|L1 | 30346 | NM_001351618 | 11201 | Hs.438533 | NM_007195 | ENSG00000101751 | POLI | protein-coding |
| chrX | 7188475 | 7188497 | + | 1076 | NA | intron (NM_001320751, intron 2 of 11) | MER20\|DNA\|hAT-Charlie | -30945 | NM_000351 | 412 | Hs.522578 | NM_000351 | ENSG00000101846 | STS | protein-coding |
| chr11 | 130482752 | 130482773 | + | 1060 | NA | Intergenic | Intergenic | 33788 | NM_139055 | 170689 | Hs.534221 | NM_139055 | ENSG00000166106 | ADAMTS15 | protein-coding |
| chr2 | 2193745 | 2193767 | + | 1054 | NA | intron (NM_001329849, intron 2 of 24) | AmnSINE1\|SINE\|5S-Deu-L2 | -125476 | NR_024468 | 730811 | Hs.629472 | NR_024468 | ENSG00000225619 | MYT1L-AS1 | ncRNA |
| chr10 | 103242519 | 103242540 | + | 1048 | NA | Intergenic | Intergenic | -3358 | NM_001143909 | 729020 | Hs.458437 | NM_001143909 | ENSG00000235376 | RPEL1 | protein-coding |
| chr8 | 96211180 | 96211201 | - | 1044 | NA | Intergenic | LTR82B\|LTR\|ERVL | 24444 | NM_001199975 | 7381 | Hs.131255 | NM_006294 | ENSG00000156467 | UQCRB | protein-coding |
| chr15 | 47980356 | 47980379 | + | 1034 | NA | Intergenic | Intergenic | -134131 | NR_120336 | 101928442 | Hs.242804 | NR_120336 | ENSG00000259572 | LINC01491 | ncRNA |
| chrX | 73606906 | 73606928 | + | 1008 | NA | intron (NM_001039840, intron 3 of 5) | L1P3\|LINE\|L1 | 43769 | NM_001039840 | 53344 | Hs.496323 | NM_001039840 | ENSG00000204116 | CHIC1 | protein-coding |
| chr3 | 134078537 | 134078559 | + | 1000 | NA | Intergenic | Charlie13a\|DNA\|hAT-Charlie | -20900 | NR_135574 | 100507210 | Hs.669052 | NR_135574 | ENSG00000243832 | LINC02000 | ncRNA |
| chr8 | 58479484 | 58479506 | - | 994 | NA | Intergenic | Intergenic | 20666 | NM_000780 | 1581 | Hs.1644 | NM_000780 | ENSG00000167910 | CYP7A1 | protein-coding |
| chr1 | 47986606 | 47986628 | + | 988 | NA | intron (NM_001194986, intron 2 of 6) | L2c\|LINE\|L2 | 10273 | NM_001194986 | 388630 | Hs.61504 | NM_001194986 | ENSG00000269113 | TRABD2B | protein-coding |
| chr13 | 36282127 | 36282149 | + | 982 | NA | intron (NM_001198910, intron 2 of 15) | intron (NM_001198910, intron 2 of 15) | 15717 | NM_001144984 | 728591 | Hs.656745 | NM_001144981 | ENSG00000242715 | CCDC169 | protein-coding |
| chr2 | 84793642 | 84793664 | + | 970 | NA | intron (NM_001370, intron 68 of 76) | intron (NM_001370, intron 68 of 76) | 87592 | NM_001277053 | 129293 | Hs.469134 | NM_001080824 | ENSG00000186854 | TRABD2A | protein-coding |
| chr18 | 61501492 | 61501514 | - | 928 | NA | intron (NM_031891, intron 4 of 11) | MIRc\|SINE\|MIR | 167921 | NM_031891 | 28316 | Hs.671510 | NM_031891 | ENSG00000101542 | CDH20 | protein-coding |
| chr4 | 92229762 | 92229784 | - | 912 | NA | Intergenic | Intergenic | 47529 | NR_125920 | 101929194 | Hs.626389 | NR_125920 | | LNCPRESS2 | ncRNA |
| chr2 | 23248540 | 23248562 | - | 910 | NA | Intergenic | Intergenic | -136876 | NM_052920 | 114818 | Hs.130593 | NM_025067 | ENSG00000119771 | KLHL29 | protein-coding |
| chr7 | 35229704 | 35229726 | - | 890 | NA | intron (NM_001077653, intron 6 of 7) | intron (NM_001077653, intron 6 of 7) | 24385 | NM_001077653 | 57057 | Hs.404167 | NM_020417 | ENSG00000164532 | TBX20 | protein-coding |
| chr2 | 148690997 | 148691019 | - | 878 | NA | intron (NM_015630, intron 2 of 13) | intron (NM_015630, intron 2 of 13) | 46017 | NM_015630 | 26122 | Hs.23270 | NM_015630 | ENSG00000135999 | EPC2 | protein-coding |
| chr2 | 77498938 | 77498960 | + | 860 | NA | intron (NR_146416, intron 1 of 1) | intron (NR_146416, intron 1 of 1) | 23484 | NM_001134745 | 80059 | Hs.285782 | NM_024993 | ENSG00000176204 | LRRTM4 | protein-coding |
| chr15 | 101227556 | 101227578 | + | 856 | NA | intron (NM_014918, intron 2 of 2) | L1M4\|LINE\|L1 | 24365 | NM_014918 | 22856 | Hs.110488 | NM_014918 | ENSG00000131873 | CHSY1 | protein-coding |
| chrY | 2885225 | 2885247 | + | 840 | NA | Intergenic | HERVK3-int\|LTR\|ERVK | 43654 | NM_001008 | 6192 | Hs.282376 | NM_001008 | ENSG00000129824 | RPS4Y1 | protein-coding |
| chr7 | 38065141 | 38065163 | - | 838 | NA | Intergenic | Intergenic | -113054 | NM_032016 | 83930 | Hs.309753 | NM_032016 | ENSG00000010270 | STARD3NL | protein-coding |
| chr3 | 75481897 | 75481919 | - | 838 | NA | intron (NR_151705, intron 2 of 2) | MLT1E1A\|LTR\|ERVL-MaLR | 46600 | NR_151705 | 107986100 | Hs.673812 | NR_151705 | ENSG00000272690 | LINC02018 | ncRNA |
| chr16 | 58514241 | 58514263 | + | 836 | NA | TTS (NR_040072) | TTS (NR_040072) | -1227 | NM_024860 | 79918 | Hs.731691 | NM_024860 | ENSG00000103037 | SETD6 | protein-coding |
| chr21 | 24965151 | 24965173 | + | 820 | NA | intron (NR_046198, intron 2 of 3) | L1MD\|LINE\|L1 | 124612 | NR_046198 | 339622 | Hs.616316 | NR_046198 | ENSG00000226983 | LINC01692 | ncRNA |
| chr8 | 77482768 | 77482790 | + | 812 | NA | Intergenic | L1MC4a\|LINE\|L1 | 32082 | NR_125830 | 102724874 | Hs.571511 | NR_125830 | | LOC102724874 | ncRNA |
| chr3 | 97138554 | 97138576 | - | 808 | NA | intron (NM_001080448, intron 3 of 17) | L1PA15-16\|LINE\|L1 | -301028 | NM_001278300 | 285220 | Hs.272208 | NM_173655 | ENSG00000080224 | EPHA6 | protein-coding |
| chr1 | 2509641 | 2509663 | - | 804 | NA | intron (NM_018216, intron 18 of 18) | intron (NM_018216, intron 18 of 18) | 16976 | NM_018216 | 55229 | Hs.26156 | NM_018216 | ENSG00000157881 | PANK4 | protein-coding |
| chr3 | 124364923 | 124364945 | + | 784 | NA | intron (NM_001024660, intron 10 of 59) | intron (NM_001024660, intron 10 of 59) | -9398 | NR_106731 | 102464832 | | NR_106731 | ENSG00000276656 | MIR6083 | ncRNA |
| chr4 | 180598421 | 180598443 | - | 778 | NA | Intergenic | MER117\|DNA\|hAT-Charlie | 560717 | NR_033918 | 728081 | Hs.535735 | NR_033918 | ENSG00000248197 | LINC00290 | ncRNA |
| chr10 | 73217988 | 73218010 | - | 776 | NA | intron (NM_173348, intron 7 of 13) | MLT1F2-int\|LTR\|ERVL-MaLR | 29332 | NM_015190 | 23234 | Hs.408577 | NM_015190 | ENSG00000213551 | DNAJC9 | protein-coding |
| chr14 | 83972018 | 83972040 | - | 772 | NA | Intergenic | LTR16C\|LTR\|ERVL | -1299903 | NR_145983 | 26843 | | NR_145983 | | SNORD3P3 | snoRNA |
| chr4 | 115309891 | 115309912 | + | 758 | NA | Intergenic | Intergenic | -196025 | NM_022569 | 64579 | Hs.591700 | NM_022569 | ENSG00000138653 | NDST4 | protein-coding |
| chr15 | 68363244 | 68363266 | - | 714 | NA | intron (NM_001004439, intron 4 of 29) | intron (NM_001004439, intron 4 of 29) | 68898 | NM_001004439 | 22801 | Hs.436416 | NM_012211 | ENSG00000137809 | ITGA11 | protein-coding |
| chr16 | 6024469 | 6024491 | - | 710 | NA | intron (NM_018723, intron 1 of 15) | intron (NM_018723, intron 1 of 15) | 4777 | NM_001308117 | 54715 | Hs.459842 | NM_018723 | ENSG00000078328 | RBFOX1 | protein-coding |
| chr2 | 241850514 | 241850536 | + | 706 | NA | 3' UTR (NM_005018, exon 5 of 5).2 | 3' UTR (NM_005018, exon 5 of 5).2 | 8381 | NM_005018 | 5133 | Hs.158297 | NM_005018 | ENSG00000188389 | PDCD1 | protein-coding |
| chr12 | 39905306 | 39905328 | - | 702 | NA | intron (NM_052885, intron 4 of 9) | intron (NM_052885, intron 4 of 9) | 200542 | NM_052885 | 114134 | Hs.558595 | NM_052885 | ENSG00000151229 | SLC2A13 | protein-coding |
| chr7 | 34730643 | 34730665 | - | 702 | NA | intron (NR_015356, intron 2 of 4) | LTR55\|LTR\|ERVL? | 27618 | NR_033665 | 404744 | Hs.487951 | NM_207283 | ENSG00000197085 | NPSR1-AS1 | ncRNA |
| chr4 | 121813911 | 121813933 | + | 700 | NA | exon (NM_005033, exon 10 of 12) | exon (NM_005033, exon 10 of 12) | 10079 | NM_001237 | 890 | Hs.58974 | NM_001237 | ENSG00000145386 | CCNA2 | protein-coding |
| chr18 | 24700709 | 24700731 | + | 688 | NA | Intergenic | L1ME4a\|LINE\|L1 | -25061 | NR_134604 | 105372028 | Hs.569901 | NR_134604 | ENSG00000266573 | LOC105372028 | ncRNA |
| chr11 | 61582799 | 61582821 | - | 684 | NA | Intergenic | Intergenic | -1938 | NM_001252065 | 9066 | Hs.131188 | NM_004200 | ENSG00000011347 | SYT7 | protein-coding |
| chr5 | 14483352 | 14483374 | + | 670 | NA | intron (NR_134469, intron 46 of 56) | intron (NR_134469, intron 46 of 56) | 19380 | NR_145815 | 109623458 | | NR_145815 | | SNORD170 | snoRNA |
| chr1 | 14689554 | 14689576 | + | 666 | NA | intron (NM_015209, intron 1 of 7) | intron (NM_015209, intron 1 of 7) | 90848 | NM_201628 | 23254 | Hs.368823 | NM_015209 | ENSG00000189337 | KAZN | protein-coding |
| chr8 | 98535997 | 98536019 | + | 662 | NA | intron (NM_001256312, intron 11 of 12) | intron (NM_001256312, intron 11 of 12) | 108986 | NM_020697 | 3788 | Hs.388045 | NM_020697 | ENSG00000156486 | KCNS2 | protein-coding |
| chr19 | 40279875 | 40279897 | + | 660 | NA | intron (NM_001243027, intron 1 of 13) | intron (NM_001243027, intron 1 of 13) | 2755 | NR_030371 | 693226 | | NR_030371 | ENSG00000207631 | MIR641 | ncRNA |
| chr13 | 76718569 | 76718591 | - | 638 | NA | Intergenic | Intergenic | 167825 | NM_138444 | 115207 | Hs.644125 | NM_138444 | ENSG00000178695 | KCTD12 | protein-coding |
| chr5 | 9429656 | 9429678 | - | 636 | NA | intron (NM_003966, intron 2 of 22) | L2a\|LINE\|L2 | -81666 | NR_109935 | 101929338 | Hs.675861 | NR_109935 | ENSG00000251370 | SEMA5A-AS1 | ncRNA |
| chr16 | 66092321 | 66092343 | + | 636 | NA | Intergenic | LTR16C\|LTR\|ERVL | -274275 | NM_001795 | 1003 | Hs.76206 | NM_001795 | ENSG00000179776 | CDH5 | protein-coding |
| chr1 | 178311702 | 178311724 | + | 634 | NA | intron (NM_170692, intron 3 of 17) | HAL1\|LINE\|L1 | -29758 | NM_004841 | 9462 | Hs.496139 | NM_004841 | ENSG00000075391 | RASAL2 | protein-coding |
| chr14 | 29040625 | 29040647 | + | 634 | NA | Intergenic | Intergenic | -254849 | NR_135255 | 105370424 | Hs.375809 | NR_135255 | ENSG00000257056 | LINC02282 | ncRNA |

<table>
<tr><td>chr5</td><td>1486864</td><td>1486886</td><td>-</td><td></td><td>634</td><td>NA</td><td>intron (NM_024830, intron 5 of 13)</td><td>intron (NM_024830, intron 5 of 13)</td><td>23981</td><td>NR_106723</td><td>102466103</td><td></td><td>NR_106723</td><td>ENSG00000273742</td><td>MIR6075</td><td>ncRNA</td></tr>
<tr><td>chr7</td><td>87835998</td><td>87836020</td><td>-</td><td></td><td>628</td><td>NA</td><td>3' UTR (NM_018843, exon 12 of 12)</td><td>3' UTR (NM_018843, exon 12 of 12)</td><td>-40220</td><td>NM_001318062</td><td>10926</td><td>Hs.485380</td><td>NM_006716</td><td>ENSG00000006634</td><td>DBF4</td><td>protein-coding</td></tr>
<tr><td>chr15</td><td>61485020</td><td>61485042</td><td>+</td><td></td><td>628</td><td>NA</td><td>Intergenic</td><td>Intergenic</td><td>-244887</td><td>NR_146448</td><td>100506530</td><td>Hs.42265</td><td>NR_146448</td><td>ENSG00000259284</td><td>LINC02349</td><td>ncRNA</td></tr>
<tr><td>chr12</td><td>29498933</td><td>29498955</td><td>+</td><td></td><td>624</td><td>NA</td><td>5' UTR (NM_001353179, exon 1 of 26)</td><td>5' UTR (NM_001353179, exon 1 of 26)</td><td>749</td><td>NM_001353179</td><td>341350</td><td>Hs.674588</td><td>NM_183378</td><td>ENSG00000187950</td><td>OVCH1</td><td>protein-coding</td></tr>
<tr><td>chr21</td><td>27479380</td><td>27479402</td><td>-</td><td></td><td>608</td><td>NA</td><td>Intergenic</td><td>MLT1A1-int|LTR|ERVL-MaLR</td><td>-242988</td><td>NR_024357</td><td>54088</td><td>Hs.570434</td><td>NR_024357</td><td>ENSG00000225298</td><td>LINC00113</td><td>ncRNA</td></tr>
<tr><td>chr4</td><td>152168981</td><td>152169003</td><td>-</td><td></td><td>608</td><td>NA</td><td>Intergenic</td><td>Intergenic</td><td>68174</td><td>NR_121627</td><td>100996286</td><td>Hs.322462</td><td>NR_121627</td><td>ENSG00000245954</td><td>LINC02273</td><td>ncRNA</td></tr>
<tr><td>chr10</td><td>83225966</td><td>83225987</td><td>+</td><td></td><td>596</td><td>NA</td><td>Intergenic</td><td>Tigger3a|DNA|TcMar-Tigger</td><td>451113</td><td>NR_134317</td><td>105378397</td><td>Hs.147725</td><td>NR_134317</td><td></td><td>LINC02650</td><td>ncRNA</td></tr>
<tr><td>chr10</td><td>29651694</td><td>29651716</td><td>+</td><td></td><td>596</td><td>NA</td><td>intron (NM_001323599, intron 3 of 38)</td><td>L2a|LINE|L2</td><td>-16733</td><td>NM_021738</td><td>6840</td><td>Hs.499209</td><td>NM_003174</td><td>ENSG00000197321</td><td>SVIL</td><td>protein-coding</td></tr>
<tr><td>chr17</td><td>63105148</td><td>63105170</td><td>-</td><td></td><td>578</td><td>NA</td><td>intron (NM_025185, intron 3 of 24)</td><td>intron (NM_025185, intron 3 of 24)</td><td>95622</td><td>NM_025185</td><td>26115</td><td>Hs.410889</td><td>NM_015623</td><td>ENSG00000170921</td><td>TANC2</td><td>protein-coding</td></tr>
<tr><td>chr8</td><td>5825789</td><td>5825811</td><td>-</td><td></td><td>560</td><td>NA</td><td>Intergenic</td><td>HERVIP10F-int|LTR|ERV1</td><td>580748</td><td>NR_040040</td><td>100287015</td><td>Hs.156928</td><td>NR_040040</td><td>ENSG00000246089</td><td>LOC100287015</td><td>ncRNA</td></tr>
<tr><td>chr2</td><td>149653429</td><td>149653451</td><td>+</td><td></td><td>560</td><td>NA</td><td>intron (NR_110240, intron 1 of 2)</td><td>intron (NR_110240, intron 1 of 2)</td><td>-65624</td><td>NM_015702</td><td>27249</td><td>Hs.5324</td><td>NM_015702</td><td>ENSG00000168288</td><td>MMADHC</td><td>protein-coding</td></tr>
<tr><td>chr20</td><td>55398553</td><td>55398575</td><td>+</td><td></td><td>560</td><td>NA</td><td>Intergenic</td><td>MLT1L|LTR|ERVL-MaLR</td><td>-24479</td><td>NR_110629</td><td>102723578</td><td>Hs.446269</td><td>NR_110629</td><td>ENSG00000235166</td><td>LINC01440</td><td>ncRNA</td></tr>
<tr><td>chr8</td><td>20430584</td><td>20430605</td><td>-</td><td></td><td>556</td><td>NA</td><td>Intergenic</td><td>Intergenic</td><td>154821</td><td>NR_047509</td><td>100874051</td><td>Hs.385799</td><td>NR_047509</td><td>ENSG00000253733</td><td>LZTS1-AS1</td><td>ncRNA</td></tr>
<tr><td>chr7</td><td>121594410</td><td>121594432</td><td>+</td><td></td><td>556</td><td>NA</td><td>Intergenic</td><td>LTR16B|LTR|ERVL</td><td>-198053</td><td>NM_014888</td><td>10447</td><td>Hs.434053</td><td>NM_014888</td><td>ENSG00000196937</td><td>FAM3C</td><td>protein-coding</td></tr>
<tr><td>chr21</td><td>20834105</td><td>20834127</td><td>-</td><td></td><td>552</td><td>NA</td><td>Intergenic</td><td>Intergenic</td><td>-31008</td><td>NR_024090</td><td>387486</td><td>Hs.576551</td><td>NR_024090</td><td>ENSG00000224924</td><td>LINC00320</td><td>ncRNA</td></tr>
<tr><td>chr10</td><td>87932580</td><td>87932602</td><td>-</td><td></td><td>530</td><td>NA</td><td>intron (NM_001304717, intron 5 of 9)</td><td>intron (NM_001304717, intron 5 of 9)</td><td>69153</td><td>NM_001304717</td><td>5728</td><td>Hs.500466</td><td>NM_000314</td><td>ENSG00000171862</td><td>PTEN</td><td>protein-coding</td></tr>
<tr><td>chr10</td><td>122301429</td><td>122301451</td><td>-</td><td></td><td>526</td><td>NA</td><td>intron (NM_144587, intron 9 of 15)</td><td>intron (NM_144587, intron 9 of 15)</td><td>30144</td><td>NM_144587</td><td>118663</td><td>Hs.422466</td><td>NM_144587</td><td>ENSG00000138152</td><td>BTBD16</td><td>protein-coding</td></tr>
<tr><td>chr13</td><td>22120675</td><td>22120697</td><td>-</td><td></td><td>524</td><td>NA</td><td>Intergenic</td><td>Intergenic</td><td>-89599</td><td>NR_103810</td><td>100506622</td><td>Hs.255773</td><td>NR_103810</td><td>ENSG00000276476</td><td>LINC00540</td><td>ncRNA</td></tr>
<tr><td>chr4</td><td>52052259</td><td>52052281</td><td>-</td><td></td><td>524</td><td>NA</td><td>intron (NM_001346103, intron 1 of 8)</td><td>intron (NM_001346103, intron 1 of 8)</td><td>939</td><td>NM_001346103</td><td>132671</td><td>Hs.527090</td><td>NM_145263</td><td>ENSG00000163071</td><td>SPATA18</td><td>protein-coding</td></tr>
<tr><td>chr1</td><td>196022319</td><td>196022341</td><td>-</td><td></td><td>524</td><td>NA</td><td>Intergenic</td><td>Intergenic</td><td>66385</td><td>NR_146895</td><td>105371673</td><td></td><td>NR_146895</td><td></td><td>LINC01724</td><td>ncRNA</td></tr>
<tr><td>chr17</td><td>82200448</td><td>82200470</td><td>+</td><td></td><td>518</td><td>NA</td><td>intron (NM_198082, intron 2 of 16)</td><td>intron (NM_198082, intron 2 of 16)</td><td>12419</td><td>NM_001316321</td><td>284001</td><td>Hs.631724</td><td>NM_152675</td><td>ENSG00000176155</td><td>CCDC57</td><td>protein-coding</td></tr>
<tr><td>chr8</td><td>81369918</td><td>81369940</td><td>+</td><td></td><td>512</td><td>NA</td><td>Intergenic</td><td>Intergenic</td><td>77594</td><td>NM_002677</td><td>5375</td><td>Hs.571512</td><td>NM_002677</td><td>ENSG00000147588</td><td>PMP2</td><td>protein-coding</td></tr>
<tr><td>chr3</td><td>29500655</td><td>29500676</td><td>-</td><td></td><td>508</td><td>NA</td><td>intron (NM_001003792, intron 3 of 13)</td><td>Tigger3a|DNA|TcMar-Tigger</td><td>219353</td><td>NM_001003792</td><td>27303</td><td>Hs.221436</td><td>NM_014483</td><td>ENSG00000144642</td><td>RBMS3</td><td>protein-coding</td></tr>
<tr><td>chr4</td><td>32261894</td><td>32261916</td><td>+</td><td></td><td>504</td><td>NA</td><td>Intergenic</td><td>Intergenic</td><td>-89133</td><td>NR_134672</td><td>101927363</td><td>Hs.559406</td><td>NR_134672</td><td>ENSG00000251329</td><td>LINC02353</td><td>ncRNA</td></tr>
<tr><td>chr7</td><td>126589477</td><td>126589498</td><td>-</td><td></td><td>502</td><td>NA</td><td>intron (NR_028041, intron 8 of 10)</td><td>L1MA8|LINE|L1</td><td>468697</td><td>NR_030323</td><td>693177</td><td></td><td>NR_030323</td><td>ENSG00000207692</td><td>MIR592</td><td>ncRNA</td></tr>
<tr><td>chrX</td><td>100249552</td><td>100249574</td><td>-</td><td></td><td>498</td><td>NA</td><td>Intergenic</td><td>MLT1L|LTR|ERVL-MaLR</td><td>160710</td><td>NM_020766</td><td>57526</td><td>Hs.4993</td><td>NM_020766</td><td>ENSG00000165194</td><td>PCDH19</td><td>protein-coding</td></tr>
<tr><td>chr11</td><td>36380525</td><td>36380547</td><td>+</td><td></td><td>496</td><td>NA</td><td>intron (NM_024841, intron 2 of 9)</td><td>intron (NM_024841, intron 2 of 9)</td><td>4551</td><td>NM_001160168</td><td>79899</td><td>Hs.19987</td><td>NM_024841</td><td>ENSG00000135362</td><td>PRR5L</td><td>protein-coding</td></tr>
<tr><td>chr16</td><td>80347415</td><td>80347437</td><td>-</td><td></td><td>496</td><td>NA</td><td>intron (NR_120307, intron 2 of 5)</td><td>L2c|LINE|L2</td><td>-193308</td><td>NM_001305017</td><td>83657</td><td>Hs.98849</td><td>NM_130897</td><td>ENSG00000168589</td><td>DYNLRB2</td><td>protein-coding</td></tr>
<tr><td>chr7</td><td>132268653</td><td>132268675</td><td>-</td><td></td><td>488</td><td>NA</td><td>intron (NM_020911, intron 4 of 31)</td><td>intron (NM_020911, intron 4 of 31)</td><td>7803</td><td>NR_134565</td><td>101928807</td><td>Hs.552131</td><td>NR_134565</td><td></td><td>LOC101928807</td><td>ncRNA</td></tr>
<tr><td>chr8</td><td>66418567</td><td>66418589</td><td>-</td><td></td><td>486</td><td>NA</td><td>Intergenic</td><td>L1MEd|LINE|L1</td><td>10399</td><td>NR_040434</td><td>100505676</td><td>Hs.121613</td><td>NM_001242759</td><td>ENSG00000246145</td><td>RRS1-AS1</td><td>ncRNA</td></tr>
<tr><td>chr15</td><td>94163544</td><td>94163566</td><td>-</td><td></td><td>484</td><td>NA</td><td>Intergenic</td><td>Intergenic</td><td>-55617</td><td>NR_120320</td><td>101927112</td><td>Hs.134687</td><td>NR_120320</td><td>ENSG00000259724</td><td>LINC01581</td><td>ncRNA</td></tr>
<tr><td>chr12</td><td>24946173</td><td>24946195</td><td>-</td><td></td><td>482</td><td>NA</td><td>intron (NM_001178091, intron 1 of 9)</td><td>intron (NM_001178091, intron 1 of 9)</td><td>3275</td><td>NM_001178092</td><td>586</td><td>Hs.438993</td><td>NM_005504</td><td>ENSG00000060982</td><td>BCAT1</td><td>protein-coding</td></tr>
<tr><td>chr4</td><td>188979583</td><td>188979605</td><td>+</td><td></td><td>474</td><td>NA</td><td>Intergenic</td><td>Intergenic</td><td>202937</td><td>NR_149102</td><td>105377610</td><td></td><td>NR_149102</td><td>ENSG00000251619</td><td>LINC02508</td><td>ncRNA</td></tr>
<tr><td>chr12</td><td>10363022</td><td>10363044</td><td>+</td><td></td><td>468</td><td>NA</td><td>promoter-TSS (NR_120430)</td><td>promoter-TSS (NR_120430)</td><td>-736</td><td>NR_120430</td><td>101928100</td><td>Hs.638425</td><td>NR_120430</td><td>ENSG00000245648</td><td>LOC101928100</td><td>ncRNA</td></tr>
<tr><td>chr2</td><td>213220681</td><td>213220703</td><td>+</td><td></td><td>464</td><td>NA</td><td>Intergenic</td><td>Intergenic</td><td>-52982</td><td>NR_146974</td><td>105373863</td><td>Hs.258045</td><td>NR_146974</td><td></td><td>LINC01953</td><td>ncRNA</td></tr>
<tr><td>chr20</td><td>38299436</td><td>38299458</td><td>+</td><td></td><td>462</td><td>NA</td><td>Intergenic</td><td>Intergenic</td><td>-4703</td><td>NM_001725</td><td>671</td><td>Hs.529019</td><td>NM_001725</td><td>ENSG00000101425</td><td>BPI</td><td>protein-coding</td></tr>
<tr><td>chr10</td><td>109832124</td><td>109832146</td><td>+</td><td></td><td>458</td><td>NA</td><td>Intergenic</td><td>Intergenic</td><td>91111</td><td>NM_001324136</td><td>7511</td><td>Hs.390623</td><td>NM_020383</td><td>ENSG00000108039</td><td>XPNPEP1</td><td>protein-coding</td></tr>
<tr><td>chr5</td><td>40821684</td><td>40821706</td><td>+</td><td></td><td>456</td><td>NA</td><td>Intergenic</td><td>MLT1A1|LTR|ERVL-MaLR</td><td>7447</td><td>NR_037665</td><td></td><td></td><td></td><td></td><td></td><td></td></tr>
<tr><td>chr3</td><td>139634833</td><td>139634854</td><td>+</td><td></td><td>452</td><td>NA</td><td>intron (NM_001320511, intron 2 of 6)</td><td>intron (NM_001320511, intron 2 of 6)</td><td>43207</td><td>NM_001320511</td><td>349565</td><td>Hs.208673</td><td>NM_178177</td><td>ENSG00000163864</td><td>NMNAT3</td><td>protein-coding</td></tr>
<tr><td>chr3</td><td>48426529</td><td>48426551</td><td>+</td><td></td><td>448</td><td>NA</td><td>intron (NM_001130082, intron 1 of 37)</td><td>intron (NM_001130082, intron 1 of 37)</td><td>2923</td><td>NM_002673</td><td>5364</td><td>Hs.476209</td><td>NM_002673</td><td>ENSG00000164050</td><td>PLXNB1</td><td>protein-coding</td></tr>
<tr><td>chr2</td><td>98356339</td><td>98356361</td><td>+</td><td></td><td>432</td><td>NA</td><td>intron (NM_001079878, intron 1 of 6)</td><td>intron (NM_001079878, intron 1 of 6)</td><td>10195</td><td>NM_001079878</td><td>1261</td><td>Hs.234785</td><td>NM_001298</td><td>ENSG00000144191</td><td>CNGA3</td><td>protein-coding</td></tr>
<tr><td>chr7</td><td>14730916</td><td>14730938</td><td>-</td><td></td><td>432</td><td>NA</td><td>intron (NM_145695, intron 4 of 23)</td><td>intron (NM_145695, intron 4 of 23)</td><td>110523</td><td>NM_145695</td><td>1607</td><td>Hs.567255</td><td>NM_004080</td><td>ENSG00000136267</td><td>DGKB</td><td>protein-coding</td></tr>
<tr><td>chr6</td><td>21698541</td><td>21698563</td><td>+</td><td></td><td>432</td><td>NA</td><td>intron (NR_015410, intron 2 of 11)</td><td>intron (NR_015410, intron 2 of 11)</td><td>32139</td><td>NR_015410</td><td>401237</td><td>Hs.712707</td><td>NR_015410</td><td></td><td>CASC15</td><td>ncRNA</td></tr>
<tr><td>chr4</td><td>75745177</td><td>75745199</td><td>+</td><td></td><td>420</td><td>NA</td><td>intron (NM_003715, intron 1 of 23)</td><td>intron (NM_003715, intron 1 of 23)</td><td>20666</td><td>NM_003715</td><td>8615</td><td>Hs.744877</td><td>NM_003715</td><td>ENSG00000138768</td><td>USO1</td><td>protein-coding</td></tr>
<tr><td>chr6</td><td>106784821</td><td>106784843</td><td>+</td><td></td><td>420</td><td>NA</td><td>TTS (NR_030314)</td><td>TTS (NR_030314)</td><td>707</td><td>NR_030314</td><td>693172</td><td></td><td>NR_030314</td><td>ENSG00000207577</td><td>MIR587</td><td>ncRNA</td></tr>
<tr><td>chr2</td><td>61385165</td><td>61385187</td><td>-</td><td></td><td>418</td><td>NA</td><td>intron (NM_014709, intron 5 of 79)</td><td>L1ME3B|LINE|L1</td><td>32201</td><td>NR_003707</td><td>100124537</td><td>Hs.675825</td><td>NR_003707</td><td>ENSG00000206937</td><td>SNORA70B</td><td>snoRNA</td></tr>
<tr><td>chr7</td><td>17260835</td><td>17260857</td><td>+</td><td></td><td>410</td><td>NA</td><td>Intergenic</td><td>Intergenic</td><td>-37806</td><td>NM_001621</td><td>196</td><td>Hs.171189</td><td>NM_001621</td><td>ENSG00000106546</td><td>AHR</td><td>protein-coding</td></tr>
<tr><td>chr6</td><td>50553837</td><td>50553859</td><td>+</td><td></td><td>408</td><td>NA</td><td>Intergenic</td><td>L2a|LINE|L2</td><td>-159696</td><td>NM_172238</td><td>83741</td><td>Hs.434107</td><td>NM_172238</td><td>ENSG00000008197</td><td>TFAP2D</td><td>protein-coding</td></tr>
<tr><td>chr4</td><td>176193527</td><td>176193549</td><td>-</td><td></td><td>402</td><td>NA</td><td>exon (NM_144644, exon 2 of 6)</td><td>exon (NM_144644, exon 2 of 6)</td><td>2133</td><td>NM_144644</td><td>132851</td><td>Hs.481235</td><td>NM_144644</td><td>ENSG00000150628</td><td>SPATA4</td><td>protein-coding</td></tr>
<tr><td>chr6</td><td>89377401</td><td>89377423</td><td>-</td><td></td><td>402</td><td>NA</td><td>intron (NM_021244, intron 5 of 6)</td><td>FLAM_A|SINE|Alu</td><td>-24512</td><td>NM_016021</td><td>51465</td><td>Hs.163776</td><td>NM_016021</td><td>ENSG00000198833</td><td>UBE2J1</td><td>protein-coding</td></tr>
<tr><td>chr2</td><td>98513989</td><td>98514011</td><td>+</td><td></td><td>398</td><td>NA</td><td>intron (NM_001351427, intron 1 of 25)</td><td>intron (NM_001351427, intron 1 of 25)</td><td>67990</td><td>NM_001351427</td><td>3631</td><td>Hs.469386</td><td>NM_001566</td><td>ENSG00000040933</td><td>INPP4A</td><td>protein-coding</td></tr>
<tr><td>chr20</td><td>47696378</td><td>47696400</td><td>+</td><td></td><td>394</td><td>NA</td><td>intron (NM_018837, intron 4 of 20)</td><td>intron (NM_018837, intron 4 of 20)</td><td>89675</td><td>NM_018837</td><td>55959</td><td>Hs.162016</td><td>NM_018837</td><td>ENSG00000196562</td><td>SULF2</td><td>protein-coding</td></tr>
<tr><td>chr8</td><td>53023386</td><td>53023408</td><td>+</td><td></td><td>388</td><td>NA</td><td>Intergenic</td><td>Intergenic</td><td>83489</td><td>NM_005285</td><td>2831</td><td>Hs.248117</td><td>NM_005285</td><td>ENSG00000183729</td><td>NPBWR1</td><td>protein-coding</td></tr>
<tr><td>chr19</td><td>13379964</td><td>13379986</td><td>-</td><td></td><td>384</td><td>NA</td><td>intron (NM_001127222, intron 3 of 46)</td><td>intron (NM_001127222, intron 3 of 46)</td><td>126485</td><td>NM_000068</td><td>773</td><td>Hs.501632</td><td>NM_000068</td><td>ENSG00000141837</td><td>CACNA1A</td><td>protein-coding</td></tr>
<tr><td>chr3</td><td>111775300</td><td>111775322</td><td>-</td><td></td><td>374</td><td>NA</td><td>intron (NM_001134437, intron 1 of 17)</td><td>intron (NM_001134437, intron 1 of 17)</td><td>42831</td><td>NM_001134437</td><td>90102</td><td>Hs.603252</td><td>NM_145753</td><td>ENSG00000144824</td><td>PHLDB2</td><td>protein-coding</td></tr>
<tr><td>chr4</td><td>137372205</td><td>137372227</td><td>+</td><td></td><td>372</td><td>NA</td><td>Intergenic</td><td>Intergenic</td><td>-159417</td><td>NR_149105</td><td>105377441</td><td>Hs.668386</td><td>NR_149105</td><td></td><td>LINC02511</td><td>ncRNA</td></tr>
<tr><td>chr6</td><td>129818166</td><td>129818188</td><td>-</td><td></td><td>372</td><td>NA</td><td>Intergenic</td><td>Intergenic</td><td>43094</td><td>NM_001010876</td><td>253582</td><td></td><td>NM_001010876</td><td>ENSG00000203756</td><td>TMEM244</td><td>protein-coding</td></tr>
<tr><td>chr1</td><td>23168498</td><td>23168520</td><td>-</td><td></td><td>362</td><td>NA</td><td>intron (NM_033631, intron 1 of 4)</td><td>CpG</td><td>349</td><td>NM_033631</td><td>7798</td><td>Hs.257900</td><td>NM_033631</td><td>ENSG00000169641</td><td>LUZP1</td><td>protein-coding</td></tr>
<tr><td>chr16</td><td>10258428</td><td>10258450</td><td>-</td><td></td><td>356</td><td>NA</td><td>Intergenic</td><td>Intergenic</td><td>-75685</td><td>NM_000833</td><td>2903</td><td>Hs.411472</td><td>NM_000833</td><td>ENSG00000183454</td><td>GRIN2A</td><td>protein-coding</td></tr>
<tr><td>chr16</td><td>76483341</td><td>76483363</td><td>+</td><td></td><td>354</td><td>NA</td><td>intron (NM_001322188, intron 12 of 24)</td><td>Tigger17a|DNA|TcMar-Tigger</td><td>-151646</td><td>NR_110934</td><td>101928203</td><td>Hs.737035</td><td>NR_110934</td><td>ENSG00000250514</td><td>LINC02125</td><td>ncRNA</td></tr>
<tr><td>chr14</td><td>62120490</td><td>62120512</td><td>-</td><td></td><td>354</td><td>NA</td><td>intron (NR_015358, intron 2 of 3)</td><td>intron (NR_015358, intron 2 of 3)</td><td>3144</td><td>NR_015358</td><td>646113</td><td>Hs.445241</td><td>NM_001039785</td><td></td><td>LINC00643</td><td>ncRNA</td></tr>
<tr><td>chr22</td><td>30628984</td><td>30629006</td><td>-</td><td></td><td>350</td><td>NA</td><td>Intergenic</td><td>Intergenic</td><td>-6820</td><td>NM_001001479</td><td>339665</td><td>Hs.660384</td><td>NM_001001479</td><td>ENSG00000100036</td><td>SLC35E4</td><td>protein-coding</td></tr>
<tr><td>chr9</td><td>27450050</td><td>27450072</td><td>+</td><td></td><td>344</td><td>NA</td><td>intron (NM_024761, intron 2 of 3)</td><td>intron (NM_024761, intron 2 of 3)</td><td>-74253</td><td>NM_020124</td><td>56832</td><td>Hs.591083</td><td>NM_020124</td><td>ENSG00000147896</td><td>IFNK</td><td>protein-coding</td></tr>
<tr><td>chr19</td><td>13079182</td><td>13079204</td><td>+</td><td></td><td>344</td><td>NA</td><td>intron (NM_001271043, intron 7 of 10)</td><td>intron (NM_001271043, intron 7 of 10)</td><td>23967</td><td>NM_005583</td><td>4066</td><td>Hs.46446</td><td>NM_005583</td><td>ENSG00000104903</td><td>LYL1</td><td>protein-coding</td></tr>
<tr><td>chr12</td><td>11572808</td><td>11572830</td><td>-</td><td></td><td>336</td><td>NA</td><td>Intergenic</td><td>Intergenic</td><td>24789</td><td>NR_033890</td><td>338817</td><td>Hs.733066</td><td>NR_033890</td><td>ENSG00000247157</td><td>LINC01252</td><td>ncRNA</td></tr>
<tr><td>chr14</td><td>20064334</td><td>20064356</td><td>+</td><td></td><td>336</td><td>NA</td><td>Intergenic</td><td>Intergenic</td><td>4300</td><td>NM_001004717</td><td>122742</td><td>Hs.553574</td><td>NM_001004717</td><td>ENSG00000176246</td><td>OR4L1</td><td>protein-coding</td></tr>
<tr><td>chr3</td><td>131478535</td><td>131478557</td><td>+</td><td></td><td>334</td><td>NA</td><td>TTS (NR_002985)</td><td>TTS (NR_002985)</td><td>687</td><td>NR_002985</td><td>677836</td><td>Hs.735762</td><td>NR_002985</td><td></td><td>SNORA58</td><td>snoRNA</td></tr>
<tr><td>chr2</td><td>128250946</td><td>128250968</td><td>-</td><td></td><td>332</td><td>NA</td><td>Intergenic</td><td>Intergenic</td><td>67640</td><td>NM_004807</td><td>9394</td><td>Hs.512841</td><td>NM_004807</td><td>ENSG00000136720</td><td>HS6ST1</td><td>protein-coding</td></tr>
<tr><td>chr1</td><td>20668626</td><td>20668648</td><td>-</td><td></td><td>326</td><td>NA</td><td>intron (NM_020816, intron 13 of 14)</td><td>intron (NM_020816, intron 13 of 14)</td><td>-7093</td><td>NM_005216</td><td>1650</td><td>Hs.523145</td><td>NM_005216</td><td>ENSG00000244038</td><td>DDOST</td><td>protein-coding</td></tr>
<tr><td>chr1</td><td>208310566</td><td>208310588</td><td>-</td><td></td><td>324</td><td>NA</td><td>Intergenic</td><td>Intergenic</td><td>-66257</td><td>NM_025179</td><td>5362</td><td>Hs.497626</td><td>NM_025179</td><td>ENSG00000076356</td><td>PLXNA2</td><td>protein-coding</td></tr>
<tr><td>chr5</td><td>91496332</td><td>91496354</td><td>-</td><td></td><td>320</td><td>NA</td><td>Intergenic</td><td>Intergenic</td><td>-112970</td><td>NR_138072</td><td>57561</td><td>Hs.24684</td><td>NM_020801</td><td>ENSG00000113369</td><td>ARRDC3</td><td>protein-coding</td></tr>
<tr><td>chr2</td><td>111912653</td><td>111912675</td><td>-</td><td></td><td>320</td><td>NA</td><td>intron (NM_006343, intron 1 of 18)</td><td>intron (NM_006343, intron 1 of 18)</td><td>14050</td><td>NM_006343</td><td>10461</td><td>Hs.306178</td><td>NM_006343</td><td>ENSG00000153208</td><td>MERTK</td><td>protein-coding</td></tr>
<tr><td>chr7</td><td>99380040</td><td>99380061</td><td>-</td><td></td><td>312</td><td>NA</td><td>intron (NM_005720, intron 1 of 9)</td><td>intron (NM_005720, intron 1 of 9)</td><td>5375</td><td>NM_005720</td><td>10095</td><td>Hs.489284</td><td>NM_005720</td><td>ENSG00000130429</td><td>ARPC1B</td><td>protein-coding</td></tr>
<tr><td>chr1</td><td>151710120</td><td>151710142</td><td>-</td><td></td><td>312</td><td>NA</td><td>intron (NM_001291107, intron 2 of 12)</td><td>intron (NM_001291107, intron 2 of 12)</td><td>6185</td><td>NM_001172648</td><td>11189</td><td>Hs.26047</td><td>NM_007185</td><td>ENSG00000159409</td><td>CELF3</td><td>protein-coding</td></tr>
<tr><td>chr10</td><td>78856714</td><td>78856736</td><td>-</td><td></td><td>310</td><td>NA</td><td>Intergenic</td><td>Intergenic</td><td>210723</td><td>NR_024431</td><td>283050</td><td>Hs.309176</td><td>NR_015429</td><td>ENSG00000224596</td><td>ZMIZ1-AS1</td><td>ncRNA</td></tr>
<tr><td>chr21</td><td>30310129</td><td>30310151</td><td>+</td><td></td><td>310</td><td>NA</td><td>Intergenic</td><td>Intergenic</td><td>10149</td><td>NM_203405</td><td>388818</td><td>Hs.28704</td><td>NM_203405</td><td>ENSG00000197683</td><td>KRTAP26-1</td><td>protein-coding</td></tr>
<tr><td>chr15</td><td>69072396</td><td>69072418</td><td>+</td><td></td><td>306</td><td>NA</td><td>Intergenic</td><td>MLT1J|LTR|ERVL-MaLR</td><td>-8443</td><td>NR_026949</td><td>283673</td><td>Hs.177927</td><td>NM_173610</td><td>ENSG00000212766</td><td>EWSAT1</td><td>ncRNA</td></tr>
<tr><td>chr11</td><td>58130840</td><td>58130862</td><td>+</td><td></td><td>304</td><td>NA</td><td>intron (NM_001005212, intron 2 of 2)</td><td>L1M5|LINE|L1</td><td>-11407</td><td>NM_001005211</td><td>219954</td><td>Hs.632122</td><td>NM_001005211</td><td>ENSG00000172377</td><td>OR9I1</td><td>protein-coding</td></tr>
<tr><td>chr2</td><td>85607917</td><td>85607939</td><td>+</td><td></td><td>304</td><td>NA</td><td>3' UTR (NM_001013649, exon 4 of 4)</td><td>3' UTR (NM_001013649, exon 4 of 4)</td><td>4128</td><td>NM_001013649</td><td>388969</td><td>Hs.516159</td><td>NM_001013649</td><td>ENSG00000168887</td><td>C2orf68</td><td>protein-coding</td></tr>
<tr><td>chr12</td><td>104076189</td><td>104076211</td><td>+</td><td></td><td>300</td><td>NA</td><td>intron (NM_013320, intron 3 of 14)</td><td>FLAM_A|SINE|Alu</td><td>11742</td><td>NM_013320</td><td>29915</td><td>Hs.506558</td><td>NM_013320</td><td>ENSG00000111727</td><td>HCFC2</td><td>protein-coding</td></tr>
<tr><td>chr4</td><td>84172290</td><td>84172312</td><td>+</td><td></td><td>298</td><td>NA</td><td>intron (NR_125909, intron 5 of 8)</td><td>MIR3|SINE|MIR</td><td>126868</td><td>NR_125909</td><td>101928978</td><td>Hs.647977</td><td>NR_125909</td><td>ENSG00000250546</td><td>LOC101928978</td><td>ncRNA</td></tr>
<tr><td>chr1</td><td>62391795</td><td>62391817</td><td>-</td><td></td><td>298</td><td>NA</td><td>Intergenic</td><td>Intergenic</td><td>-44498</td><td>NM_001017416</td><td>7398</td><td>Hs.35086</td><td>NM_003368</td><td>ENSG00000162607</td><td>USP1</td><td>protein-coding</td></tr>
<tr><td>chr15</td><td>101752655</td><td>101752678</td><td>-</td><td></td><td>296</td><td>NA</td><td>Intergenic</td><td>CpG</td><td>-5796</td><td>NM_001321551</td><td>100128108</td><td>Hs.104864</td><td>NM_001321551</td><td></td><td>LOC100128108</td><td>protein-coding</td></tr>
<tr><td>chr5</td><td>244291</td><td>244313</td><td>+</td><td></td><td>296</td><td>NA</td><td>intron (NM_001330758, intron 11 of 12)</td><td>HERVK9-int|LTR|ERVK</td><td>26079</td><td>NM_001294332</td><td>6389</td><td>Hs.440475</td><td>NM_004168</td><td>ENSG00000073578</td><td>SDHA</td><td>protein-coding</td></tr>
<tr><td>chr3</td><td>151873530</td><td>151873552</td><td>+</td><td></td><td>292</td><td>NA</td><td>promoter-TSS (NM_033050)</td><td>promoter-TSS (NM_033050)</td><td>-102</td><td>NM_033050</td><td>56670</td><td>Hs.133181</td><td>NM_033050</td><td>ENSG00000198829</td><td>SUCNR1</td><td>protein-coding</td></tr>
<tr><td>chrY</td><td>19970709</td><td>19970731</td><td>-</td><td></td><td>292</td><td>NA</td><td>Intergenic</td><td>L1PA8A|LINE|L1</td><td>-225781</td><td>NM_004653</td><td>8284</td><td>Hs.80358</td><td>NM_004653</td><td>ENSG00000012817</td><td>KDM5D</td><td>protein-coding</td></tr>
<tr><td>chr9</td><td>103677082</td><td>103677103</td><td>+</td><td></td><td>292</td><td>NA</td><td>Intergenic</td><td>Intergenic</td><td>-322617</td><td>NR_109803</td><td>101928523</td><td></td><td>NR_109803</td><td>ENSG00000285483</td><td>LOC101928523</td><td>ncRNA</td></tr>
<tr><td>chr10</td><td>105608207</td><td>105608229</td><td>-</td><td></td><td>292</td><td>NA</td><td>Intergenic</td><td>MER67D|LTR|ERV1</td><td>212115</td><td>NR_120625</td><td>101927549</td><td>Hs.556885</td><td>NR_120625</td><td></td><td>LINC02627</td><td>ncRNA</td></tr>
<tr><td>chr9</td><td>28413657</td><td>28413679</td><td>+</td><td></td><td>290</td><td>NA</td><td>intron (NM_152570, intron 3 of 6)</td><td>intron (NM_152570, intron 3 of 6)</td><td>305637</td><td>NM_152570</td><td>158038</td><td>Hs.444665</td><td>NM_152570</td><td>ENSG00000174482</td><td>LINGO2</td><td>protein-coding</td></tr>
<tr><td>chr10</td><td>115878525</td><td>115878547</td><td>+</td><td></td><td>290</td><td>NA</td><td>intron (NM_207303, intron 28 of 28)</td><td>intron (NM_207303, intron 28 of 28)</td><td>316253</td><td>NR_145811</td><td>109623470</td><td></td><td>NR_145811</td><td></td><td>SNORD158</td><td>snoRNA</td></tr>
<tr><td>chrX</td><td>126431052</td><td>126431074</td><td>+</td><td></td><td>290</td><td>NA</td><td>Intergenic</td><td>MSTD-int|LTR|ERVL-MaLR</td><td>121796</td><td>NM_178470</td><td>139170</td><td>Hs.120403</td><td>NM_178470</td><td>ENSG00000198889</td><td>DCAF12L1</td><td>protein-coding</td></tr>
<tr><td>chr8</td><td>101291420</td><td>101291442</td><td>+</td><td></td><td>288</td><td>NA</td><td>Intergenic</td><td>Intergenic</td><td>-77462</td><td>NR_002182</td><td>83955</td><td>Hs.567608</td><td>NM_032031</td><td>ENSG00000228224</td><td>NACA4P</td><td>pseudo</td></tr>
<tr><td>chr10</td><td>20198078</td><td>20198100</td><td>+</td><td></td><td>284</td><td>NA</td><td>intron (NM_032812, intron 9 of 13)</td><td>intron (NM_032812, intron 9 of 13)</td><td>-353881</td><td>NR_039822</td><td>100616383</td><td></td><td>NR_039822</td><td>ENSG00000265372</td><td>MIR4675</td><td>ncRNA</td></tr>
<tr><td>chr10</td><td>88377497</td><td>88377518</td><td>+</td><td></td><td>280</td><td>NA</td><td>intron (NM_018363, intron 4 of 6)</td><td>Tigger3b|DNA|TcMar-Tigger</td><td>205818</td><td>NM_018363</td><td>55328</td><td>Hs.149849</td><td>NM_018363</td><td>ENSG00000184719</td><td>RNLS</td><td>protein-coding</td></tr>
<tr><td>chr1</td><td>54479589</td><td>54479611</td><td>+</td><td></td><td>276</td><td>NA</td><td>Intergenic</td><td>Intergenic</td><td>-68534</td><td>NM_147161</td><td>26027</td><td>Hs.745173</td><td>NM_015547</td><td>ENSG00000162390</td><td>ACOT11</td><td>protein-coding</td></tr>
<tr><td>chr9</td><td>36140302</td><td>36140324</td><td>-</td><td></td><td>274</td><td>NA</td><td>intron (NM_001287014, intron 1 of 1)</td><td>intron (NM_001287014, intron 1 of 1)</td><td>3528</td><td>NM_001287013</td><td>152007</td><td>Hs.493819</td><td>NM_022343</td><td>ENSG00000122694</td><td>GLIPR2</td><td>protein-coding</td></tr>
<tr><td>chr13</td><td>35449637</td><td>35449659</td><td>+</td><td></td><td>268</td><td>NA</td><td>intron (NM_015678, intron 39 of 57)</td><td>L4_C_Mam|LINE|RTE-X</td><td>27047</td><td>NM_005584</td><td>4081</td><td>Hs.584776</td><td>NM_005584</td><td>ENSG00000180660</td><td>MAB21L1</td><td>protein-coding</td></tr>
<tr><td>chr14</td><td>29067071</td><td>29067093</td><td>+</td><td></td><td>266</td><td>NA</td><td>Intergenic</td><td>LTR16C|LTR|ERVL</td><td>-281295</td><td>NR_135255</td><td>105370424</td><td>Hs.375809</td><td>NR_135255</td><td>ENSG00000257056</td><td>LINC02282</td><td>ncRNA</td></tr>
<tr><td>chr4</td><td>174324044</td><td>174324066</td><td>+</td><td></td><td>264</td><td>NA</td><td>intron (NM_001145314, intron 10 of 10)</td><td>intron (NM_001145314, intron 10 of 10)</td><td>-39804</td><td>NM_012180</td><td>26269</td><td>Hs.76917</td><td>NM_012180</td><td>ENSG00000164117</td><td>FBXO8</td><td>protein-coding</td></tr>
<tr><td>chr8</td><td>67074392</td><td>67074414</td><td>+</td><td></td><td>262</td><td>NA</td><td>intron (NM_024790, intron 2 of 28)</td><td>intron (NM_024790, intron 2 of 28)</td><td>-9445</td><td>NM_001291339</td><td>79848</td><td>Hs.370147</td><td>NM_024790</td><td>ENSG00000104218</td><td>CSPP1</td><td>protein-coding</td></tr>
<tr><td>chr1</td><td>189451116</td><td>189451138</td><td>-</td><td></td><td>254</td><td>NA</td><td>Intergenic</td><td>MLT1E2|LTR|ERVL-MaLR</td><td>1024883</td><td>NM_001317188</td><td>339479</td><td>Hs.65765</td><td>NM_199051</td><td>ENSG00000162670</td><td>BRINP3</td><td>protein-coding</td></tr>
<tr><td>chr3</td><td>183115459</td><td>183115481</td><td>-</td><td></td><td>252</td><td>NA</td><td>Intergenic</td><td>ERV3-16A3_I-int|LTR|ERVL</td><td>-15883</td><td>NM_020166</td><td>56922</td><td>Hs.47649</td><td>NM_020166</td><td>ENSG00000078070</td><td>MCCC1</td><td>protein-coding</td></tr>
<tr><td>chr2</td><td>24024799</td><td>24024821</td><td>-</td><td></td><td>252</td><td>NA</td><td>intron (NM_001346880, intron 13 of 13)</td><td>L2b|LINE|L2</td><td>14723</td><td>NM_001346880</td><td>388931</td><td>Hs.407482</td><td>NM_001080473</td><td>ENSG00000205639</td><td>MFSD2B</td><td>protein-coding</td></tr>
<tr><td>chr11</td><td>95385187</td><td>95385209</td><td>-</td><td></td><td>252</td><td>NA</td><td>Intergenic</td><td>Intergenic</td><td>-152657</td><td>NM_001271594</td><td>143686</td><td>Hs.120633</td><td>NM_144665</td><td>ENSG00000149212</td><td>SESN3</td><td>protein-coding</td></tr>
<tr><td>chr18</td><td>9600227</td><td>9600248</td><td>-</td><td></td><td>250</td><td>NA</td><td>intron (NM_001042388, intron 2 of 19)</td><td>L1MEf|LINE|L1</td><td>14370</td><td>NM_005134</td><td>9989</td><td>Hs.743236</td><td>NM_005134</td><td>ENSG00000154845</td><td>PPP4R1</td><td>protein-coding</td></tr>
<tr><td>chr10</td><td>13519170</td><td>13519192</td><td>+</td><td></td><td>250</td><td>NA</td><td>Intergenic</td><td>Intergenic</td><td>-16205</td><td>NM_152751</td><td>222389</td><td>Hs.498740</td><td>NM_152751</td><td>ENSG00000165626</td><td>BEND7</td><td>protein-coding</td></tr>
<tr><td>chr11</td><td>58667575</td><td>58667597</td><td>+</td><td></td><td>246</td><td>NA</td><td>Intergenic</td><td>SATR1|Satellite|Satellite</td><td>44913</td><td>NM_000614</td><td>1270</td><td>Hs.524920</td><td>NM_000614</td><td>ENSG00000242689</td><td>CNTF</td><td>protein-coding</td></tr>
<tr><td>chr11</td><td>97711764</td><td>97711785</td><td>+</td><td></td><td>244</td><td>NA</td><td>Intergenic</td><td>MER110-int|LTR|ERV1</td><td>1203349</td><td>NR_135097</td><td>105369443</td><td>Hs.599377</td><td>NR_135097</td><td>ENSG00000256684</td><td>LINC02737</td><td>ncRNA</td></tr>
<tr><td>chr8</td><td>19729488</td><td>19729511</td><td>-</td><td></td><td>244</td><td>NA</td><td>intron (NM_001354475, intron 1 of 9)</td><td>L3|LINE|CR1</td><td>27584</td><td>NM_001354490</td><td>55790</td><td>Hs.613729</td><td>NM_018371</td><td>ENSG00000147408</td><td>CSGALNACT1</td><td>protein-coding</td></tr>
<tr><td>chrX</td><td>134156285</td><td>134156307</td><td>+</td><td></td><td>242</td><td>NA</td><td>Intergenic</td><td>Intergenic</td><td>13156</td><td>NR_029852</td><td>574031</td><td></td><td>NR_029852</td><td>ENSG00000284499</td><td>MIR363</td><td>ncRNA</td></tr>
<tr><td>chr1</td><td>204829726</td><td>204829748</td><td>+</td><td></td><td>242</td><td>NA</td><td>intron (NM_001005388, intron 1 of 29)</td><td>intron (NM_001005388, intron 1 of 29)</td><td>1083</td><td>NM_015090</td><td>23114</td><td>Hs.13349</td><td>NM_015090</td><td>ENSG00000163531</td><td>NFASC</td><td>protein-coding</td></tr>
<tr><td>chr1</td><td>202431661</td><td>202431683</td><td>+</td><td></td><td>242</td><td>NA</td><td>intron (NM_001167858, intron 8 of 8)</td><td>intron (NM_001167858, intron 8 of 8)</td><td>-31071</td><td>NM_032103</td><td>4660</td><td>Hs.444403</td><td>NM_002481</td><td>ENSG00000077157</td><td>PPP1R12B</td><td>protein-coding</td></tr>
<tr><td>chr4</td><td>9998679</td><td>9998701</td><td>-</td><td></td><td>240</td><td>NA</td><td>intron (NM_001001290, intron 3 of 12)</td><td>L1ME3B|LINE|L1</td><td>22800</td><td>NM_020041</td><td>56606</td><td>Hs.444612</td><td>NM_020041</td><td>ENSG00000109667</td><td>SLC2A9</td><td>protein-coding</td></tr>
<tr><td>chr17</td><td>15458489</td><td>15458511</td><td>+</td><td></td><td>236</td><td>NA</td><td>intron (NM_001204477, intron 1 of 3)</td><td>intron (NM_001204477, intron 1 of 3)</td><td>9111</td><td>NM_001204477</td><td>284040</td><td>Hs.164595</td><td>NM_173622</td><td>ENSG00000239704</td><td>CDRT4</td><td>protein-coding</td></tr>
<tr><td>chr2</td><td>216160902</td><td>216160924</td><td>+</td><td></td><td>234</td><td>NA</td><td>intron (NM_021141, intron 15 of 20)</td><td>AluJo|SINE|Alu</td><td>51616</td><td>NM_021141</td><td>7520</td><td>Hs.388739</td><td>NM_021141</td><td>ENSG00000079246</td><td>XRCC5</td><td>protein-coding</td></tr>
<tr><td>chr2</td><td>160340433</td><td>160340455</td><td>-</td><td></td><td>228</td><td>NA</td><td>intron (NM_016836, intron 2 of 13)</td><td>intron (NM_016836, intron 2 of 13)</td><td>67438</td><td>NR_039946</td><td>100616364</td><td></td><td>NR_039946</td><td>ENSG00000283734</td><td>MIR4785</td><td>ncRNA</td></tr>
<tr><td>chr3</td><td>165149249</td><td>165149271</td><td>-</td><td></td><td>228</td><td>NA</td><td>Intergenic</td><td>Intergenic</td><td>46742</td><td>NM_001318810</td><td>22865</td><td>Hs.101745</td><td>NM_014926</td><td>ENSG00000121871</td><td>SLITRK3</td><td>protein-coding</td></tr>
<tr><td>chr6</td><td>93734875</td><td>93734897</td><td>-</td><td></td><td>228</td><td>NA</td><td>intron (NR_152795, intron 3 of 6)</td><td>MLT1E|LTR|ERVL-MaLR</td><td>16583</td><td>NR_152795</td><td>643432</td><td>Hs.509936</td><td>NR_015362</td><td></td><td>TSG1</td><td>ncRNA</td></tr>
<tr><td>chr2</td><td>109210656</td><td>109210678</td><td>+</td><td></td><td>226</td><td>NA</td><td>intron (NM_001099289, intron 1 of 9)</td><td>intron (NM_001099289, intron 1 of 9)</td><td>-69079</td><td>NR_036223</td><td>100422863</td><td></td><td>NR_036223</td><td>ENSG00000264934</td><td>MIR4265</td><td>ncRNA</td></tr>
<tr><td>chr2</td><td>12647861</td><td>12647883</td><td>+</td><td></td><td>226</td><td>NA</td><td>Intergenic</td><td>Intergenic</td><td>-69000</td><td>NM_021643</td><td>28951</td><td>Hs.467751</td><td>NM_021643</td><td>ENSG00000071575</td><td>TRIB2</td><td>protein-coding</td></tr>
<tr><td>chr8</td><td>11494738</td><td>11494760</td><td>+</td><td></td><td>226</td><td>NA</td><td>intron (NM_001330465, intron 1 of 11)</td><td>intron (NM_001330465, intron 1 of 11)</td><td>737</td><td>NM_001715</td><td>640</td><td>Hs.146591</td><td>NM_001715</td><td>ENSG00000136573</td><td>BLK</td><td>protein-coding</td></tr>
<tr><td>chr1</td><td>225099686</td><td>225099708</td><td>-</td><td></td><td>226</td><td>NA</td><td>intron (NM_001373, intron 22 of 83)</td><td>intron (NM_001373, intron 22 of 83)</td><td>170043</td><td>NM_001145154</td><td>127602</td><td>Hs.133977</td><td>NM_001373</td><td>ENSG00000185842</td><td>DNAH14</td><td>protein-coding</td></tr>
<tr><td>chr11</td><td>109632317</td><td>109632339</td><td>+</td><td></td><td>226</td><td>NA</td><td>Intergenic</td><td>L1MA8|LINE|L1</td><td>210208</td><td>NM_207645</td><td>399947</td><td>Hs.172982</td><td>NM_207645</td><td>ENSG00000185742</td><td>C11orf87</td><td>protein-coding</td></tr>
<tr><td>chr20</td><td>19068845</td><td>19068866</td><td>-</td><td></td><td>222</td><td>NA</td><td>Intergenic</td><td>LTR33|LTR|ERVL</td><td>-143791</td><td>NM_020689</td><td>57419</td><td>Hs.654790</td><td>NM_020689</td><td>ENSG00000185052</td><td>SLC24A3</td><td>protein-coding</td></tr>
<tr><td>chr1</td><td>52568613</td><td>52568635</td><td>-</td><td></td><td>220</td><td>NA</td><td>Intergenic</td><td>Intergenic</td><td>-15534</td><td>NM_015269</td><td>23318</td><td>Hs.655407</td><td>NM_015269</td><td>ENSG00000134744</td><td>TUT4</td><td>protein-coding</td></tr>
<tr><td>chr2</td><td>240006057</td><td>240006079</td><td>-</td><td></td><td>220</td><td>NA</td><td>intron (NM_001322019, intron 7 of 9)</td><td>intron (NM_001322019, intron 7 of 9)</td><td>19334</td><td>NM_004544</td><td>4705</td><td>Hs.277677</td><td>NM_004544</td><td>ENSG00000130414</td><td>NDUFA10</td><td>protein-coding</td></tr>
<tr><td>chr10</td><td>102965463</td><td>102965485</td><td>-</td><td></td><td>220</td><td>NA</td><td>intron (NM_199076, intron 1 of 6)</td><td>intron (NM_199076, intron 1 of 6)</td><td>47156</td><td>NM_017649</td><td>54805</td><td>Hs.643509</td><td>NM_017649</td><td>ENSG00000148842</td><td>CNNM2</td><td>protein-coding</td></tr>
</table>

| | | | | | | | | | | | | | | | | |
|---|---|---|---|---|---|---|---|---|---|---|---|---|---|---|---|---|
| chr12 | 26478422 | 26478444 | - | | 218 | NA | intron (NM_002223, intron 43 of 56) | intron (NM_002223, intron 43 of 56) | 282860 | NM_005086 | 8082 | Hs.183428 | NM_005086 | ENSG00000123096 | SSPN | protein-coding |
| chr12 | 63713610 | 63713632 | - | | 218 | NA | Intergenic | Intergenic | -45047 | NM_173812 | 283417 | Hs.533644 | NM_173812 | ENSG00000177990 | DPY19L2 | protein-coding |
| chr11 | 90453328 | 90453350 | + | | 214 | NA | intron (NR_104190, intron 1 of 6) | MLT1C-int\|LTR\|ERVL-MaLR | 102518 | NR_039711 | 100616186 | | NR_039711 | ENSG00000266703 | MIR4490 | ncRNA |
| chr2 | 21790371 | 21790393 | - | | 208 | NA | Intergenic | Intergenic | -79730 | NR_038837 | 645949 | Hs.456782 | NR_038837 | ENSG00000229621 | LINC01822 | ncRNA |
| chr2 | 49966720 | 49966742 | - | | 204 | NA | intron (NM_001330096, intron 15 of 16) | intron (NM_001330096, intron 15 of 16) | 7472 | NM_001320157 | 9378 | Hs.637685 | NM_004801 | ENSG00000179915 | NRXN1 | protein-coding |
| chr7 | 31864795 | 31864817 | + | | 204 | NA | intron (NM_001191056, intron 7 of 16) | intron (NM_001191056, intron 7 of 16) | 177789 | NM_001145123 | 10842 | Hs.227011 | NM_006658 | ENSG00000106341 | PPP1R17 | protein-coding |
| chr18 | 8297934 | 8297957 | + | | 204 | NA | intron (NM_001105244, intron 20 of 32) | intron (NM_001105244, intron 20 of 32) | 69089 | NR_024419 | 100192426 | Hs.666655 | NR_024419 | ENSG00000266149 | LOC100192426 | ncRNA |
| chr2 | 80569264 | 80569286 | + | | 198 | NA | intron (NM_004389, intron 12 of 17) | intron (NM_004389, intron 12 of 17) | 255896 | NM_001282599 | 1496 | Hs.167368 | NM_004389 | ENSG00000066032 | CTNNA2 | protein-coding |
| chr17 | 18228280 | 18228302 | + | | 198 | NA | intron (NM_004140, intron 1 of 22) | L2a\|LINE\|L2 | 2669 | NM_004140 | 3996 | Hs.513983 | NM_004140 | ENSG00000131899 | LLGL1 | protein-coding |
| chr2 | 73766480 | 73766503 | + | | 194 | NA | exon (NM_003584, exon 8 of 9) | exon (NM_003584, exon 8 of 9) | 13666 | NM_003584 | 8446 | Hs.14611 | NM_003584 | ENSG00000144048 | DUSP11 | protein-coding |
| chr3 | 123546157 | 123546179 | - | | 192 | NA | intron (NM_001329786, intron 2 of 5) | intron (NM_001329786, intron 2 of 5) | 38449 | NM_001329787 | 201562 | Hs.705480 | NM_198402 | ENSG00000206527 | HACD2 | protein-coding |
| chr12 | 84994926 | 84994948 | + | | 190 | NA | Intergenic | Intergenic | 41340 | NM_001100917 | 144448 | Hs.156962 | NM_001100917 | ENSG00000231738 | TSPAN19 | protein-coding |
| chr9 | 90814950 | 90814972 | - | | 188 | NA | intron (NM_001174168, intron 1 of 12) | intron (NM_001174168, intron 1 of 12) | -12459 | NM_001174167 | 6850 | Hs.371720 | NM_003177 | ENSG00000165025 | SYK | protein-coding |
| chr8 | 83593590 | 83593612 | + | | 188 | NA | Intergenic | Intergenic | 189843 | NR_122034 | 103352670 | Hs.399852 | NR_122034 | ENSG00000253898 | LINC01419 | ncRNA |
| chr2 | 16753222 | 16753244 | - | | 186 | NA | Intergenic | HERVL-int\|LTR\|ERVL | -87367 | NM_030797 | 81553 | Hs.467769 | NM_030797 | ENSG00000197872 | FAM49A | protein-coding |
| chr6 | 9071559 | 9071580 | - | | 184 | NA | Intergenic | Tigger3a\|DNA\|TcMar-Tigger | 419360 | NR_004855 | 728655 | Hs.214343 | NR_004855 | | HULC | ncRNA |
| chr13 | 32228077 | 32228099 | - | | 182 | NA | intron (NM_023037, intron 39 of 60) | intron (NM_023037, intron 39 of 60) | 83866 | NM_001136571 | 646799 | Hs.569254 | NM_001136571 | ENSG00000189167 | ZAR1L | protein-coding |
| chr7 | 27077223 | 27077245 | - | | 180 | NA | Intergenic | MIR3\|SINE\|MIR | 18772 | NM_153620 | 3198 | Hs.67397 | NM_005522 | ENSG00000105991 | HOXA1 | protein-coding |
| chr10 | 62422785 | 62422807 | + | | 178 | NA | intron (NM_199451, intron 3 of 7) | ERVL-E-int\|LTR\|ERVL | -47669 | NR_104162 | 283045 | | NR_104162 | | LOC283045 | ncRNA |
| chr14 | 66383386 | 66383408 | - | | 178 | NA | Intergenic | Intergenic | -102974 | NR_110308 | 440184 | Hs.412138 | NM_001004331 | ENSG00000196553 | CCDC196 | protein-coding |
| chr9 | 90792361 | 90792383 | - | | 176 | NA | Intergenic | LTR8A\|LTR\|ERV1 | -9308 | NM_003177 | 6850 | Hs.371720 | NM_003177 | ENSG00000165025 | SYK | protein-coding |
| chr9 | 84059595 | 84059617 | - | | 174 | NA | Intergenic | Intergenic | -3837 | NR_110995 | 101927575 | Hs.459826 | NR_110995 | ENSG00000227463 | LOC101927575 | ncRNA |
| chr9 | 81092582 | 81092604 | - | | 170 | NA | Intergenic | Intergenic | 596942 | NM_005077 | 7088 | Hs.197320 | NM_005077 | ENSG00000196781 | TLE1 | protein-coding |
| chr11 | 7269464 | 7269486 | - | | 170 | NA | intron (NM_175733, intron 1 of 6) | MLT1J\|LTR\|ERVL-MaLR | 17525 | NM_175733 | 143425 | Hs.177193 | NM_175733 | ENSG00000170743 | SYT9 | protein-coding |
| chr1 | 117541711 | 117541733 | + | | 168 | NA | Intergenic | Intergenic | 64048 | NR_121626 | 100996263 | Hs.202533 | NR_121626 | ENSG00000236866 | LOC100996263 | ncRNA |
| chr13 | 71045378 | 71045400 | + | | 166 | NA | intron (NR_047699, intron 1 of 3) | intron (NR_047699, intron 1 of 3) | 30248 | NR_047699 | 100885781 | Hs.372660 | NR_047699 | ENSG00000226846 | LINC00348 | ncRNA |
| chr5 | 115709474 | 115709495 | - | | 166 | NA | Intergenic | Intergenic | -36832 | NR_104674 | 102467217 | Hs.147738 | NR_104674 | | LOC102467217 | ncRNA |
| chr12 | 69408151 | 69408173 | + | | 164 | NA | Intergenic | LTR89\|LTR\|ERVL? | 48452 | NM_001300950 | 8089 | Hs.4029 | NM_006530 | ENSG00000127337 | YEATS4 | protein-coding |
| chr1 | 21807806 | 21807828 | - | | 164 | NA | Intergenic | Arthur1\|DNA\|hAT-Tip100 | -4448 | NM_001013693 | 401944 | Hs.745158 | NM_001013693 | ENSG00000187942 | LDLRAD2 | protein-coding |
| chr7 | 15126823 | 15126845 | - | | 164 | NA | Intergenic | MER4E\|LTR\|ERV1 | -152057 | NM_001350722 | 1607 | Hs.567255 | NM_004080 | ENSG00000136267 | DGKB | protein-coding |
| chr21 | 40605398 | 40605420 | - | | 164 | NA | intron (NM_001271534, intron 3 of 32) | L1MC2\|LINE\|L1 | 25358 | NR_046774 | 100874326 | Hs.679746 | NR_046774 | ENSG00000233756 | DSCAM-IT1 | ncRNA |
| chr8 | 112856893 | 112856915 | - | | 162 | NA | intron (NM_198124, intron 12 of 71) | intron (NM_198124, intron 12 of 71) | 213411 | NR_031745 | 100302225 | | NR_031745 | ENSG00000238399 | MIR2053 | ncRNA |
| chr8 | 109009125 | 109009147 | - | | 162 | NA | Intergenic | Intergenic | -78288 | NM_003301 | 7201 | Hs.3022 | NM_003301 | ENSG00000174417 | TRHR | protein-coding |
| chr1 | 10532037 | 10532059 | + | | 160 | NA | intron (NM_004565, intron 2 of 8) | intron (NM_004565, intron 2 of 8) | 57102 | NM_004565 | 5195 | Hs.149983 | NM_004565 | ENSG00000142655 | PEX14 | protein-coding |
| chr4 | 29306678 | 29306700 | - | | 160 | NA | Intergenic | LTR102_Mam\|LTR\|ERVL | 92397 | NR_146996 | 105374561 | Hs.582013 | NR_146996 | ENSG00000248338 | LINC02472 | ncRNA |
| chr19 | 44475522 | 44475544 | - | | 160 | NA | 3' UTR (NM_001288762, exon 4 of 4) | 3' UTR (NM_001288762, exon 4 of 4) | 24990 | NM_001278509 | 7733 | Hs.22305 | NM_013256 | ENSG00000167384 | ZNF180 | protein-coding |
| chrX | 18129675 | 18129697 | + | | 160 | NA | Intergenic | Intergenic | -25042 | NR_133641 | 105373144 | | NR_133641 | ENSG00000225882 | LINC01456 | ncRNA |
| chr15 | 68037871 | 68037893 | - | | 158 | NA | Intergenic | L1M2\|LINE\|L1 | -16297 | NM_016166 | 8554 | Hs.162458 | NM_016166 | ENSG00000033800 | PIAS1 | protein-coding |
| chr20 | 5838945 | 5838967 | - | | 158 | NA | intron (NM_001303478, intron 3 of 3) | intron (NM_001303478, intron 3 of 3) | -72372 | NM_001819 | 1114 | Hs.516874 | NM_001819 | ENSG00000089199 | CHGB | protein-coding |
| chr2 | 217018446 | 217018468 | - | | 156 | NA | Intergenic | Intergenic | 147685 | NR_130782 | 101928327 | Hs.350698 | NR_130782 | ENSG00000223874 | LINC01921 | ncRNA |
| chr3 | 67725838 | 67725859 | + | | 154 | NA | intron (NR_109992, intron 3 of 6) | intron (NR_109992, intron 3 of 6) | 71151 | NR_109992 | 101927111 | Hs.518057 | NR_109992 | ENSG00000241316 | SUCLG2-AS1 | ncRNA |
| chr2 | 180618368 | 180618390 | - | | 154 | NA | Intergenic | Intergenic | -73725 | NR_104319 | 101669767 | | NR_104319 | ENSG00000281131 | SCHLAP1 | ncRNA |
| chr3 | 7974407 | 7974429 | - | | 154 | NA | intron (NR_110131, intron 2 of 3) | intron (NR_110131, intron 2 of 3) | 41889 | NR_110131 | 101927394 | Hs.517767 | NR_110131 | ENSG00000227110 | LOC101927394 | ncRNA |
| chr17 | 69306840 | 69306862 | + | | 152 | NA | exon (NM_172232, exon 6 of 39) | exon (NM_172232, exon 6 of 39) | 8782 | NM_018672 | 23461 | Hs.731824 | NM_018672 | ENSG00000154265 | ABCA5 | protein-coding |
| chr10 | 58691368 | 58691390 | + | | 150 | NA | intron (NM_001080512, intron 2 of 20) | intron (NM_001080512, intron 2 of 20) | -23636 | NR_027508 | 728640 | Hs.729209 | NR_027508 | | FAM133CP | pseudo |
| chr9 | 136833902 | 136833924 | + | | 150 | NA | exon (NM_001173989, exon 8 of 8) | exon (NM_001173989, exon 8 of 8) | 2956 | NR_036251 | 100422860 | | NR_036251 | ENSG00000265806 | MIR4292 | ncRNA |
| chr1 | 119882094 | 119882116 | - | | 148 | NA | Intergenic | Intergenic | 14419 | NM_021794 | 11085 | Hs.283011 | NM_021794 | ENSG00000134249 | ADAM30 | protein-coding |
| chr2 | 159924922 | 159924945 | + | | 148 | NA | Intergenic | Intergenic | -20177 | NM_001198760 | 100526664 | Hs.153563 | NM_001198759 | ENSG00000248672 | LY75-CD302 | protein-coding |
| chr1 | 7688847 | 7688869 | - | | 146 | NA | intron (NM_001349613, intron 1 of 11) | intron (NM_001349613, intron 1 of 11) | 8770 | NM_001349613 | 23261 | Hs.397705 | NM_015215 | ENSG00000171735 | CAMTA1 | protein-coding |
| chr6 | 3239501 | 3239523 | + | | 146 | NA | non-coding (NR_147505, exon 2 of 2) | non-coding (NR_147505, exon 2 of 2) | 8109 | NR_147505 | 100422781 | Hs.664539 | NR_147505 | | LOC100422781 | ncRNA |
| chr5 | 93399652 | 93399674 | + | | 146 | NA | Intergenic | Intergenic | 171680 | NR_109825 | 441094 | Hs.457407 | NR_015369 | ENSG00000237187 | NR2F1-AS1 | ncRNA |
| chr1 | 172528460 | 172528481 | - | | 146 | NA | Intergenic | Intergenic | -3879 | NM_016227 | 51430 | Hs.204559 | NM_014283 | ENSG00000094975 | SUCO | protein-coding |
| chr6 | 33360785 | 33360807 | - | | 142 | NA | Intergenic | Intergenic | -30740 | NM_002263 | 3833 | Hs.436912 | NM_002263 | ENSG00000237649 | KIFC1 | protein-coding |
| chr19 | 12772996 | 12773018 | - | | 142 | NA | exon (NM_013312, exon 4 of 23) | exon (NM_013312, exon 4 of 23) | 2613 | NM_013312 | 29911 | Hs.30792 | NM_013312 | ENSG00000095066 | HOOK2 | protein-coding |
| chr5 | 50805001 | 50805022 | + | | 142 | NA | intron (NM_001178055, intron 15 of 26) | L1MB5\|LINE\|L1 | 137465 | NM_001331028 | 79668 | Hs.369581 | NM_024615 | ENSG00000151883 | PARP8 | protein-coding |
| chr13 | 79231200 | 79231222 | - | | 140 | NA | Intergenic | Intergenic | 175010 | NM_001286632 | 64062 | Hs.558528 | NM_018605 | ENSG00000139746 | RBM26 | protein-coding |
| chr5 | 163589113 | 163589135 | + | | 140 | NA | Intergenic | L2b\|LINE\|L2 | 83596 | NM_013283 | 27430 | Hs.54642 | NM_013283 | ENSG00000038274 | MAT2B | protein-coding |
| chr10 | 130157603 | 130157625 | - | | 136 | NA | intron (NM_006541, intron 2 of 10) | MLT2B3\|LTR\|ERVL | 21205 | NM_001321980 | 10539 | Hs.42644 | NM_006541 | ENSG00000108010 | GLRX3 | protein-coding |
| chr7 | 23875975 | 23875997 | - | | 132 | NA | Intergenic | Intergenic | 165660 | NM_001260504 | 56164 | Hs.309767 | NM_031414 | ENSG00000196335 | STK31 | protein-coding |
| chr5 | 105684572 | 105684594 | - | | 130 | NA | Intergenic | L1M5\|LINE\|L1 | 585109 | NR_000039 | 9366 | Hs.158296 | NR_000039 | ENSG00000232159 | RAB9BP1 | pseudo |
| chr5 | 15961251 | 15961273 | - | | 128 | NA | Intergenic | MLT2B3\|LTR\|ERVL | 26080 | NR_030616 | 100126347 | | NR_030616 | ENSG00000216077 | MIR887 | ncRNA |
| chr5 | 20187984 | 20188006 | - | | 124 | NA | intron (NM_001349558, intron 3 of 15) | intron (NM_001349558, intron 3 of 15) | -117556 | NR_146519 | 102725105 | | NR_146519 | | CDH18-AS1 | ncRNA |
| chr8 | 68709283 | 68709305 | - | | 124 | NA | intron (NM_001349478, intron 8 of 12) | intron (NM_001349478, intron 8 of 12) | -377803 | NR_038877 | 286189 | Hs.593427 | NR_038877 | ENSG00000248801 | C8orf34-AS1 | ncRNA |
| chr12 | 91545345 | 91545367 | + | | 122 | NA | Intergenic | L1PA13\|LINE\|L1 | -362232 | NM_001920 | 1634 | Hs.156316 | NM_001920 | ENSG00000011465 | DCN | protein-coding |
| chr2 | 65527925 | 65527947 | - | | 122 | NA | Intergenic | Intergenic | -95414 | NM_181784 | 200734 | Hs.59332 | NM_181784 | ENSG00000198369 | SPRED2 | protein-coding |
| chr10 | 129921694 | 129921716 | + | | 122 | NA | intron (NM_001005463, intron 6 of 15) | intron (NM_001005463, intron 6 of 15) | 42122 | NM_001005463 | 253738 | Hs.591374 | NM_001005463 | ENSG00000108001 | EBF3 | protein-coding |
| chr2 | 97349324 | 97349346 | - | | 120 | NA | Intergenic | Intergenic | 65400 | NR_130704 | 100506123 | Hs.720604 | NR_040097 | | LOC100506123 | ncRNA |
| chr9 | 96384047 | 96384068 | - | | 120 | NA | promoter-TSS (NM_001286990) | promoter-TSS (NM_001286990) | -347 | NR_104627 | 11046 | Hs.494556 | NM_007001 | ENSG00000130958 | SLC35D2 | protein-coding |
| chr13 | 108028909 | 108028931 | - | | 120 | NA | Intergenic | Intergenic | -161808 | NM_001080396 | 728215 | Hs.535394 | NM_001080396 | ENSG00000204442 | FAM155A | protein-coding |
| chr14 | 54208894 | 54208916 | + | | 120 | NA | Intergenic | MLT1L\|LTR\|ERVL-MaLR | -187963 | NM_001330173 | 1033 | Hs.84113 | NM_005192 | ENSG00000100526 | CDKN3 | protein-coding |
| chr7 | 158446638 | 158446660 | - | | 118 | NA | intron (NM_130842, intron 1 of 21) | intron (NM_130842, intron 1 of 21) | 86164 | NR_030325 | 693180 | | NR_030325 | ENSG00000207637 | MIR595 | ncRNA |
| chr15 | 77018640 | 77018662 | + | | 118 | NA | intron (NM_001321137, intron 4 of 15) | intron (NM_001321137, intron 4 of 15) | 23566 | NM_001321136 | 9051 | Hs.129758 | NM_003978 | ENSG00000140368 | PSTPIP1 | protein-coding |
| chr5 | 100028305 | 100028327 | - | | 116 | NA | Intergenic | L1MA3\|LINE\|L1 | 359938 | NR_027503 | 100133050 | Hs.534796 | NR_027503 | | LOC100133050 | pseudo |
| chr11 | 82556867 | 82556889 | + | | 116 | NA | Intergenic | Intergenic | -152965 | NR_120571 | 101928989 | Hs.145117 | NR_120571 | ENSG00000245832 | MIR4300HG | ncRNA |
| chr10 | 252494 | 252516 | - | | 116 | NA | 3' UTR (NM_006624, exon 15 of 15) | 3' UTR (NM_006624, exon 15 of 15) | 29581 | NR_106988 | | | | | | |
| chr21 | 31462451 | 31462473 | - | | 116 | NA | intron (NM_001353691, intron 2 of 29) | intron (NM_001353691, intron 2 of 29) | 96625 | NM_001353689 | 7074 | Hs.517228 | NM_003253 | ENSG00000156299 | TIAM1 | protein-coding |
| chr7 | 141838297 | 141838319 | - | | 114 | NA | intron (NM_001008270, intron 2 of 4) | MIR\|SINE\|MIR | 3113 | NM_001171951 | 136242 | Hs.98947 | NM_001008270 | ENSG00000165076 | PRSS37 | protein-coding |
| chr7 | 124399416 | 124399437 | + | | 114 | NA | Intergenic | Tigger3a\|DNA\|TcMar-Tigger | -47330 | NR_110180 | 101928211 | Hs.131334 | NR_110180 | | LOC101928211 | ncRNA |
| chr7 | 86157290 | 86157312 | - | | 114 | NA | Intergenic | L1ME2z\|LINE\|L1 | -486613 | NM_000840 | 2913 | Hs.590575 | NM_000840 | ENSG00000198822 | GRM3 | protein-coding |
| chr1 | 178953139 | 178953161 | - | | 112 | NA | Intergenic | Intergenic | -72654 | NM_001324310 | 9917 | Hs.5737 | NM_014864 | ENSG00000116199 | FAM20B | protein-coding |
| chr10 | 89132853 | 89132875 | - | | 112 | NA | Intergenic | MER110A\|LTR\|ERV1 | -69453 | NR_039827 | 100616192 | | NR_039827 | ENSG00000265375 | MIR4679-2 | ncRNA |
| chr10 | 75706437 | 75706458 | + | | 112 | NA | intron (NR_131178, intron 1 of 7) | intron (NR_131178, intron 1 of 7) | -6008 | NR_132791 | 105378367 | Hs.464836 | NR_132791 | | LOC105378367 | ncRNA |
| chr5 | 137861769 | 137861791 | - | | 112 | NA | Intergenic | Intergenic | -6076 | NM_006790 | 9499 | Hs.84665 | NM_006790 | ENSG00000120729 | MYOT | protein-coding |
| chr7 | 70783312 | 70783334 | + | | 110 | NA | intron (NM_001127231, intron 14 of 17) | L3\|LINE\|CR1 | -349214 | NM_022479 | 64409 | Hs.488591 | NM_022479 | ENSG00000185274 | GALNT17 | protein-coding |
| chrX | 103268122 | 103268143 | - | | 110 | NA | Intergenic | L1M4b\|LINE\|L1 | 8737 | NM_001012979 | 340543 | Hs.403462 | NM_001012979 | ENSG00000204065 | TCEAL5 | protein-coding |
| chr19 | 29637160 | 29637182 | + | | 108 | NA | Intergenic | Eulor6B\|DNA?\|DNA? | -28249 | NM_024310 | 79156 | Hs.466383 | NM_024310 | ENSG00000166289 | PLEKHF1 | protein-coding |
| chrX | 45945430 | 45945452 | - | | 106 | NA | Intergenic | Intergenic | -93951 | NR_125365 | 401585 | Hs.633348 | NR_125365 | ENSG00000231566 | LINC02595 | ncRNA |
| chr18 | 55079103 | 55079125 | + | | 104 | NA | Intergenic | MLT1C\|LTR\|ERVL-MaLR | 45192 | NR_110743 | 101927229 | Hs.638474 | NR_110743 | ENSG00000267013 | LINC01929 | ncRNA |
| chr6 | 156594002 | 156594024 | - | | 104 | NA | Intergenic | MLT1E1A\|LTR\|ERVL-MaLR | -183842 | NM_001346813 | 57492 | Hs.291587 | NM_017519 | ENSG00000049618 | ARID1B | protein-coding |
| chr5 | 139585646 | 139585668 | - | | 102 | NA | intron (NM_181838, intron 2 of 7) | intron (NM_181838, intron 2 of 7) | 24491 | NM_003339 | 7322 | Hs.108332 | NM_003339 | ENSG00000131508 | UBE2D2 | protein-coding |
| chr20 | 17010458 | 17010480 | + | | 102 | NA | Intergenic | LTR48B\|LTR\|ERV1 | -215638 | NM_001201528 | 5126 | Hs.315186 | NM_002594 | ENSG00000125851 | PCSK2 | protein-coding |
| chr5 | 56556170 | 56556192 | + | | 100 | NA | 5' UTR (NM_001287053, exon 2 of 6) | 5' UTR (NM_001287053, exon 2 of 6) | 50051 | NM_001287053 | 101928448 | Hs.208052 | NM_001287053 | ENSG00000225940 | C5orf67 | ncRNA |
| chr10 | 29958549 | 29958571 | + | | 100 | NA | Intergenic | L2b\|LINE\|L2 | 101026 | NM_020848 | 57608 | Hs.533953 | NM_020848 | ENSG00000165757 | JCAD | protein-coding |
| chr5 | 56026937 | 56026959 | - | | 98 | NA | Intergenic | Intergenic | 31781 | NR_102755 | 441072 | Hs.482141 | NR_102755 | ENSG00000227908 | FLJ31104 | ncRNA |
| chr8 | 108256432 | 108256454 | + | | 98 | NA | Intergenic | L2c\|LINE\|L2 | -7713 | NM_001568 | 3646 | Hs.405590 | NM_001568 | ENSG00000104408 | EIF3E | protein-coding |
| chr16 | 18816443 | 18816465 | + | | 98 | NA | exon (NM_015092, exon 58 of 63) | exon (NM_015092, exon 58 of 63) | -14776 | NM_015161 | 23204 | Hs.634882 | NM_015161 | ENSG00000170540 | ARL6IP1 | protein-coding |
| chr1 | 52396414 | 52396435 | - | | 96 | NA | intron (NM_001190818, intron 4 of 16) | intron (NM_001190818, intron 4 of 16) | 8047 | NM_001190818 | 4998 | Hs.17908 | NM_004153 | ENSG00000085840 | ORC1 | protein-coding |
| chr12 | 5152594 | 5152616 | - | | 96 | NA | Intergenic | Charlie16a\|DNA\|hAT-Charlie | 90546 | NR_120477 | 101929584 | Hs.436529 | NR_120476 | ENSG00000256115 | LINC02443 | ncRNA |
| chr2 | 177121888 | 177121910 | + | | 94 | NA | Intergenic | Intergenic | -90795 | NM_001330251 | 220988 | Hs.516539 | NM_194247 | ENSG00000170144 | HNRNPA3 | protein-coding |
| chr4 | 53631255 | 53631277 | + | | 94 | NA | Intergenic | Intergenic | 38310 | NR_046827 | 100873955 | Hs.351074 | NR_046827 | | LNX1-AS2 | ncRNA |
| chr11 | 72107696 | 72107718 | - | | 92 | NA | intron (NM_001330321, intron 5 of 5) | intron (NM_001330321, intron 5 of 5) | -4320 | NM_017907 | 55004 | Hs.731528 | NM_017907 | ENSG00000149357 | LAMTOR1 | protein-coding |
| chr8 | 83692818 | 83692841 | + | | 92 | NA | Intergenic | Intergenic | 289071 | NR_122034 | 103352670 | Hs.399852 | NR_122034 | ENSG00000253898 | LINC01419 | ncRNA |
| chr3 | 117651545 | 117651567 | + | | 90 | NA | Intergenic | Intergenic | 39369 | NR_146652 | 105374057 | Hs.518179 | NR_146652 | ENSG00000241213 | LINC02024 | ncRNA |
| chr5 | 38052954 | 38052976 | + | | 88 | NA | intron (NR_147009, intron 2 of 2) | intron (NR_147009, intron 2 of 2) | 27268 | NR_147009 | 105374731 | Hs.570981 | NR_147009 | ENSG00000248461 | LINC02107 | ncRNA |
| chr11 | 95756604 | 95756626 | - | | 86 | NA | Intergenic | Intergenic | 33175 | NM_144664 | 143684 | Hs.288304 | NM_144664 | ENSG00000077458 | FAM76B | protein-coding |
| chr5 | 10583932 | 10583954 | + | | 86 | NA | intron (NM_001164440, intron 1 of 3) | intron (NM_001164440, intron 1 of 3) | 19620 | NM_001164440 | 651746 | Hs.26039 | NM_001164440 | ENSG00000164236 | ANKRD33B | protein-coding |
| chr4 | 17362022 | 17362044 | + | | 86 | NA | Intergenic | L2b\|LINE\|L2 | -41151 | NR_145770 | 109616994 | | NR_145770 | ENSG00000206780 | SNORA75B | snoRNA |
| chr2 | 38308953 | 38308975 | - | | 84 | NA | intron (NM_001330460, intron 8 of 12) | intron (NM_001330460, intron 8 of 12) | 67566 | NM_001308076 | 64225 | Hs.594950 | NM_022374 | ENSG00000119787 | ATL2 | protein-coding |
| chr5 | 26967403 | 26967425 | + | | 84 | NA | intron (NM_016279, intron 2 of 11) | intron (NM_016279, intron 2 of 11) | 71168 | NM_016279 | 1007 | Hs.272212 | NM_016279 | ENSG00000113100 | CDH9 | protein-coding |
| chr15 | 101764037 | 101764060 | - | | 82 | NA | Intergenic | CpG | -17178 | NM_001321551 | 100128108 | Hs.104864 | NM_001321551 | | LOC100128108 | protein-coding |
| chr12 | 42301305 | 42301327 | - | | 82 | NA | Intergenic | Intergenic | 22543 | NR_107005 | 102467003 | | NR_107005 | ENSG00000278520 | MIR7851 | ncRNA |
| chr19 | 8586068 | 8586090 | - | | 80 | NA | intron (NM_001282352, intron 9 of 12) | intron (NM_001282352, intron 9 of 12) | 6117 | NM_001282352 | 81794 | Hs.657508 | NM_030957 | ENSG00000142303 | ADAMTS10 | protein-coding |
| chr9 | 39002898 | 39002920 | + | | 80 | NA | Intergenic | Intergenic | 285394 | NM_033655 | 79937 | Hs.128474 | NM_033655 | ENSG00000106714 | CNTNAP3 | protein-coding |
| chr9 | 73959012 | 73959034 | - | | 80 | NA | Intergenic | MER102a\|DNA\|hAT-Charlie | -412508 | NR_121182 | 101927358 | Hs.559523 | NR_121182 | ENSG00000227809 | LOC101927358 | ncRNA |
| chr16 | 29860389 | 29860411 | - | | 78 | NA | intron (NM_001286586, intron 4 of 5) | intron (NM_001286586, intron 4 of 5) | 2888 | NM_001286586 | 10423 | Hs.121549 | NM_006319 | ENSG00000103502 | CDIPT | protein-coding |
| chr20 | 36903222 | 36903244 | - | | 76 | NA | intron (NM_015474, intron 13 of 15) | L2\|LINE\|L2 | 27112 | NM_080628 | 140711 | Hs.472630 | NM_080628 | ENSG00000101342 | TLDC2 | protein-coding |
| chr2 | 161568894 | 161568916 | + | | 76 | NA | Intergenic | Intergenic | -55430 | NM_001354441 | 57282 | Hs.333958 | NM_022058 | ENSG00000144290 | SLC4A10 | protein-coding |
| chr15 | 87919315 | 87919337 | - | | 70 | NA | intron (NM_001320135, intron 15 of 15) | intron (NM_001320135, intron 15 of 15) | 286648 | NM_001320135 | 4916 | Hs.185701 | NM_002530 | ENSG00000140538 | NTRK3 | protein-coding |
| chr7 | 134031565 | 134031587 | + | | 70 | NA | intron (NM_021807, intron 17 of 17) | intron (NM_021807, intron 17 of 17) | -95776 | NM_144648 | 136332 | Hs.149774 | NM_144648 | ENSG00000155530 | LRGUK | protein-coding |
| chrX | 147281516 | 147281538 | - | | 70 | NA | TTS (NR_030240) | TTS (NR_030240) | 503 | NR_030240 | 574518 | | NR_030240 | ENSG00000207867 | MIR514A3 | ncRNA |
| chr2 | 70791380 | 70791402 | + | | 70 | NA | promoter-TSS (NM_001004311) | promoter-TSS (NM_001004311) | -748 | NM_001004311 | 344018 | Hs.407636 | NM_001004311 | ENSG00000183733 | FIGLA | protein-coding |
| chrX | 126191670 | 126191692 | + | | 70 | NA | Intergenic | Intergenic | -25584 | NM_001013628 | 340578 | Hs.181867 | NM_001013628 | ENSG00000198354 | DCAF12L2 | protein-coding |
| chr10 | 86011612 | 86011634 | + | | 70 | NA | intron (NM_017551, intron 4 of 15) | intron (NM_017551, intron 4 of 15) | 253165 | NR_029907 | 442911 | | NR_029907 | ENSG00000199104 | MIR346 | ncRNA |
| chr8 | 20665220 | 20665242 | + | | 70 | NA | Intergenic | HAL1b\|LINE\|L1 | -50305 | NR_145737 | 109616983 | | NR_145737 | | SNORD3F | snoRNA |
| chr11 | 84864963 | 84864985 | - | | 70 | NA | intron (NM_001351274, intron 5 of 26) | intron (NM_001351274, intron 5 of 26) | 58447 | NM_001364 | 1740 | Hs.367656 | NM_001364 | ENSG00000150672 | DLG2 | protein-coding |
| chr2 | 135967814 | 135967836 | + | | 68 | NA | intron (NM_001349, intron 3 of 15) | intron (NM_001349, intron 3 of 15) | -17351 | NR_110200 | 101928243 | Hs.602178 | NR_110199 | ENSG00000231890 | DARS-AS1 | ncRNA |
| chr15 | 101757471 | 101757494 | - | | 68 | NA | Intergenic | CpG | -10612 | NM_001321551 | 100128108 | Hs.104864 | NM_001321551 | | LOC100128108 | protein-coding |

| | | | | | | | | | | | | | | | | |
|---|---|---|---|---|---|---|---|---|---|---|---|---|---|---|---|---|
| chr17 | 28624832 | 28624854 | - | | 66 | NA | intron (NM_014680, intron 24 of 38) | intron (NM_014680, intron 24 of 38) | -10650 | NM_001174103 | 124923 | Hs.729077 | NM_144610 | ENSG00000167524 | RSKR | protein-coding |
| chrX | 13237000 | 13237022 | - | | 66 | NA | Intergenic | Intergenic | 66441 | NR_146309 | 109729169 | Hs.348675 | NR_146309 | | LINC02154 | ncRNA |
| chr6 | 47146790 | 47146812 | + | | 66 | NA | Intergenic | Intergenic | -104438 | NM_025048 | 266977 | Hs.733762 | NM_025048 | ENSG00000153292 | ADGRF1 | protein-coding |
| chr10 | 61995111 | 61995133 | + | | 66 | NA | intron (NM_032199, intron 3 of 9) | intron (NM_032199, intron 3 of 9) | -54089 | NM_001244638 | 84159 | Hs.535297 | NM_032199 | ENSG00000150347 | ARID5B | protein-coding |
| chr6 | 40464854 | 40464876 | - | | 64 | NA | intron (NM_020737, intron 1 of 2) | LTR16C\|LTR\|ERVL | 86528 | NM_001359096 | 732253 | Hs.180197 | NM_001359094 | | TDRG1 | ncRNA |
| chr13 | 85120391 | 85120413 | + | | 64 | NA | intron (NR_126383, intron 1 of 5) | L1PA5\|LINE\|L1 | 27655 | NR_126383 | 104355140 | | NR_126383 | ENSG00000226370 | LINC00375 | ncRNA |
| chr1 | 169192579 | 169192601 | - | | 64 | NA | intron (NR_104229, intron 10 of 12) | intron (NR_104229, intron 10 of 12) | 85881 | NM_001677 | 481 | Hs.291196 | NM_001677 | ENSG00000143153 | ATP1B1 | protein-coding |
| chr5 | 167925195 | 167925217 | - | | 64 | NA | intron (NM_001122679, intron 3 of 28) | L1ME2\|LINE\|L1 | 170270 | NM_001080428 | 57451 | Hs.631957 | NM_001080428 | ENSG00000145934 | TENM2 | protein-coding |
| chr10 | 16170752 | 16170774 | - | | 64 | NA | Intergenic | HERVL-int\|LTR\|ERVL | -266180 | NM_001261838 | 9317 | Hs.444321 | NM_030664 | ENSG00000165983 | PTER | protein-coding |
| chr11 | 85008225 | 85008247 | + | | 62 | NA | intron (NM_001351274, intron 5 of 26) | intron (NM_001351274, intron 5 of 26) | -84815 | NM_001300983 | 1740 | Hs.367656 | NM_001364 | ENSG00000150672 | DLG2 | protein-coding |
| chr7 | 131095362 | 131095384 | - | | 62 | NA | intron (NR_109852, intron 2 of 5) | intron (NR_109852, intron 2 of 5) | 11657 | NR_024153 | 378805 | Hs.150556 | NM_001085379 | ENSG00000231721 | LINC-PINT | ncRNA |
| chr7 | 107452263 | 107452285 | + | | 60 | NA | intron (NM_181733, intron 6 of 20) | L2c\|LINE\|L2 | -17783 | NM_005295 | 2845 | Hs.657277 | NM_005295 | ENSG00000172209 | GPR22 | protein-coding |
| chr5 | 123803534 | 123803557 | - | | 60 | NA | Intergenic | Intergenic | 258128 | NM_004384 | 1456 | Hs.129206 | NM_004384 | ENSG00000151292 | CSNK1G3 | protein-coding |
| chr5 | 166246739 | 166246760 | + | | 60 | NA | Intergenic | Intergenic | 679621 | NR_108020 | 102557615 | | NR_108020 | ENSG00000254130 | LINC01947 | ncRNA |
| chr5 | 144181990 | 144182012 | + | | 58 | NA | intron (NM_020768, intron 2 of 3) | AluSq\|SINE\|Alu | 11128 | NM_020768 | 57528 | Hs.7093 | NM_020768 | ENSG00000183775 | KCTD16 | protein-coding |
| chr3 | 11891707 | 11891729 | - | | 58 | NA | Intergenic | Intergenic | -44799 | NR_104314 | 132001 | Hs.475472 | NM_138807 | ENSG00000144559 | TAMM41 | protein-coding |
| chr8 | 10957542 | 10957564 | - | | 56 | NA | intron (NM_173683, intron 1 of 2) | intron (NM_173683, intron 1 of 2) | 77749 | NR_030328 | 693183 | | NR_030328 | ENSG00000207600 | MIR598 | ncRNA |
| chr16 | 88020046 | 88020068 | + | | 56 | NA | intron (NM_001173542, intron 7 of 12) | intron (NM_001173542, intron 7 of 12) | 50039 | NM_001173543 | 54971 | Hs.461705 | NM_017869 | ENSG00000172530 | BANP | protein-coding |
| chr3 | 57546802 | 57546823 | + | | 56 | NA | Intergenic | MLT1E3\|LTR\|ERVL-MaLR | -2468 | NM_001291661 | 201625 | Hs.201378 | NM_178504 | ENSG00000174844 | DNAH12 | protein-coding |
| chr18 | 30733671 | 30733693 | + | | 56 | NA | Intergenic | L1PA8A\|LINE\|L1 | 309133 | NM_001941 | 1825 | Hs.41690 | NM_001941 | ENSG00000134762 | DSC3 | protein-coding |
| chr1 | 246607669 | 246607691 | - | | 54 | NA | intron (NM_152609, intron 2 of 10) | MER48\|LTR\|ERV1 | 41343 | NM_152609 | 163882 | Hs.368353 | NM_152609 | ENSG00000162852 | CNST | protein-coding |
| chr13 | 99969033 | 99969055 | - | | 52 | NA | non-coding (NR_146225, exon 1 of 2) | non-coding (NR_146225, exon 1 of 2) | 128 | NR_146225 | 85416 | Hs.508570 | NM_033132 | ENSG00000139800 | ZIC5 | protein-coding |
| chr1 | 110718446 | 110718468 | + | | 52 | NA | Intergenic | Intergenic | -43424 | NM_002232 | 3738 | Hs.169948 | NM_002232 | ENSG00000177272 | KCNA3 | protein-coding |
| chr3 | 39495064 | 39495085 | - | | 52 | NA | intron (NR_103506, intron 2 of 2) | L1M5\|LINE\|L1 | -6992 | NM_001278323 | 4336 | Hs.121333 | NM_182935 | ENSG00000168314 | MOBP | protein-coding |
| chr12 | 119283597 | 119283619 | - | | 50 | NA | TTS (NR_024246) | TTS (NR_024246) | 19772 | NR_024246 | 144742 | Hs.524782 | NR_024246 | ENSG00000281196 | LINC00934 | ncRNA |
| chr12 | 49636103 | 49636125 | - | | 50 | NA | intron (NM_001031698, intron 15 of 25) | intron (NM_001031698, intron 15 of 25) | 12494 | NM_012272 | 25766 | Hs.706827 | NM_012272 | ENSG00000110844 | PRPF40B | protein-coding |
| chr2 | 194650032 | 194650054 | - | | 50 | NA | Intergenic | L1ME3B\|LINE\|L1 | -80552 | NR_110223 | 101927431 | Hs.558215 | NR_110223 | ENSG00000230173 | LINC01790 | ncRNA |
| chr10 | 70147172 | 70147194 | + | | 50 | NA | promoter-TSS (NM_001040273) | promoter-TSS (NM_001040273) | -443 | NR_073580 | 219743 | Hs.533655 | NM_173555 | ENSG00000156521 | TYSND1 | protein-coding |
| chr22 | 33478731 | 33478753 | - | | 50 | NA | intron (NM_133642, intron 6 of 14) | MIRb\|SINE\|MIR | -42073 | NR_039921 | 100616295 | | NR_039921 | ENSG00000266012 | MIR4764 | ncRNA |
| chr2 | 50727678 | 50727700 | - | | 50 | NA | intron (NM_001330086, intron 4 of 22) | intron (NM_001330086, intron 4 of 22) | -31427 | NR_130470 | 103504737 | | NR_130470 | ENSG00000216191 | MIR8485 | ncRNA |
| chr3 | 42166632 | 42166654 | - | | 48 | NA | intron (NM_014965, intron 1 of 12) | intron (NM_014965, intron 1 of 12) | 6474 | NM_014965 | 22906 | Hs.535711 | NM_014965 | ENSG00000182606 | TRAK1 | protein-coding |
| chr10 | 33801073 | 33801095 | + | | 48 | NA | Intergenic | Intergenic | -28404 | NR_038932 | 100505583 | Hs.568804 | NR_038932 | ENSG00000261683 | LINC00838 | ncRNA |
| chr11 | 86478474 | 86478496 | + | | 48 | NA | intron (NM_001014811, intron 6 of 13) | intron (NM_001014811, intron 6 of 13) | 103749 | NM_001156474 | 60494 | Hs.144913 | NM_021827 | ENSG00000149201 | CCDC81 | protein-coding |
| chr20 | 44766977 | 44766999 | + | | 48 | NA | intron (NM_182970, intron 2 of 5) | LTR16C\|LTR\|ERVL | -20761 | NR_132377 | 106144538 | Hs.255479 | NR_132377 | | KCNK15-AS1 | ncRNA |
| chr7 | 27186242 | 27186264 | + | | 48 | NA | non-coding (NR_002795, exon 1 of 2) | non-coding (NR_002795, exon 1 of 2) | 845 | NR_002795 | 221883 | Hs.587427 | NR_002795 | ENSG00000240990 | HOXA11-AS | ncRNA |
| chr13 | 105990247 | 105990269 | + | | 46 | NA | Intergenic | Intergenic | 283428 | NR_046391 | 144920 | Hs.567700 | NR_046391 | | LINC00343 | ncRNA |
| chr1 | 241676993 | 241677015 | - | | 46 | NA | intron (NM_144625, intron 4 of 26) | intron (NM_144625, intron 4 of 26) | 24726 | NM_144625 | 128025 | Hs.97527 | NM_144625 | ENSG00000162843 | WDR64 | protein-coding |
| chr15 | 88007972 | 88007994 | - | | 46 | NA | intron (NM_001007156, intron 14 of 15) | intron (NM_001007156, intron 14 of 15) | 197991 | NM_001320135 | 4916 | Hs.185701 | NM_002530 | ENSG00000140538 | NTRK3 | protein-coding |
| chr2 | 228220471 | 228220493 | - | | 46 | NA | Intergenic | MLT1C\|LTR\|ERVL-MaLR | -38837 | NM_001142644 | 80309 | Hs.436306 | NM_030623 | ENSG00000153820 | SPHKAP | protein-coding |
| chr1 | 69828288 | 69828310 | - | | 46 | NA | intron (NM_020794, intron 3 of 24) | L1ME1\|LINE\|L1 | 68124 | NM_020794 | 57554 | Hs.479658 | NM_020794 | ENSG00000033122 | LRRC7 | protein-coding |
| chr7 | 57328083 | 57328105 | + | | 46 | NA | Intergenic | MIR\|SINE\|MIR | -76673 | NR_120505 | 100653233 | Hs.127379 | NR_120505 | ENSG00000260653 | LOC100653233 | ncRNA |
| chr6 | 152594471 | 152594493 | + | | 46 | NA | intron (NM_182961, intron 3 of 145) | intron (NM_182961, intron 3 of 145) | 42369 | NM_033071 | 23345 | Hs.12967 | NM_015293 | ENSG00000131018 | SYNE1 | protein-coding |
| chr10 | 118328893 | 118328915 | - | | 44 | NA | intron (NM_001134672, intron 5 of 7) | L2c\|LINE\|L2 | 13423 | NM_022063 | 63877 | Hs.105575 | NM_022063 | ENSG00000165669 | FAM204A | protein-coding |
| chr4 | 42592733 | 42592754 | + | | 44 | NA | intron (NM_006095, intron 6 of 36) | Tigger3a\|DNA\|TcMar-Tigger | 64362 | NM_006095 | 10396 | Hs.435052 | NM_006095 | ENSG00000124406 | ATP8A1 | protein-coding |
| chr6 | 82048790 | 82048812 | - | | 44 | NA | intron (NR_149135, intron 3 of 3) | MER5B\|DNA\|hAT-Charlie | 52964 | NR_149135 | 102724168 | Hs.485841 | NR_149134 | | LINC02542 | ncRNA |
| chr7 | 115777075 | 115777097 | - | | 44 | NA | Intergenic | Intergenic | 191227 | NM_001244583 | 22797 | Hs.125962 | NM_012252 | ENSG00000105967 | TFEC | protein-coding |
| chr16 | 81290343 | 81290365 | - | | 44 | NA | exon (NM_017429, exon 11 of 11) | exon (NM_017429, exon 11 of 11) | -24612 | NM_022041 | 8139 | Hs.112569 | NM_022041 | ENSG00000261609 | GAN | protein-coding |
| chrX | 123136947 | 123136969 | - | | 44 | NA | Intergenic | Intergenic | -47285 | NM_007325 | 2892 | Hs.377070 | NM_000828 | ENSG00000125675 | GRIA3 | protein-coding |
| chr8 | 117783060 | 117783082 | - | | 42 | NA | Intergenic | Intergenic | 137842 | NR_145799 | 109623457 | | NR_145799 | | SNORD168 | snoRNA |
| chr2 | 237446726 | 237446748 | - | | 42 | NA | Intergenic | Intergenic | -32530 | NM_057166 | 1293 | Hs.233240 | NM_004369 | ENSG00000163359 | COL6A3 | protein-coding |
| chr9 | 76958295 | 76958317 | - | | 42 | NA | Intergenic | L2b\|LINE\|L2 | -52219 | NM_001308048 | 158471 | Hs.262857 | NM_015225 | ENSG00000106772 | PRUNE2 | protein-coding |
| chr2 | 224715349 | 224715371 | - | | 40 | NA | Intergenic | Intergenic | -129963 | NM_001257197 | 8452 | Hs.372286 | NM_003590 | ENSG00000036257 | CUL3 | protein-coding |
| chr20 | 4179992 | 4180014 | + | | 40 | NA | intron (NM_175841, intron 3 of 3) | intron (NM_175841, intron 3 of 3) | -13087 | NR_033917 | 728228 | Hs.636379 | NR_033917 | ENSG00000230176 | LINC01433 | ncRNA |
| chr2 | 13256290 | 13256312 | + | | 40 | NA | Intergenic | L3\|LINE\|CR1 | -249288 | NR_038434 | 100506474 | Hs.242196 | NR_038434 | ENSG00000225649 | LOC100506474 | ncRNA |
| chr11 | 16042189 | 16042211 | - | | 40 | NA | intron (NM_001145811, intron 11 of 14) | MIRb\|SINE\|MIR | -72579 | NR_106721 | 102464826 | | NR_106721 | ENSG00000284094 | MIR6073 | ncRNA |
| chr2 | 168285120 | 168285141 | - | | 38 | NA | Intergenic | BLACKJACK\|DNA\|hAT-Blackjack | -37535 | NM_013233 | 27347 | Hs.276271 | NM_013233 | ENSG00000198648 | STK39 | protein-coding |
| chr8 | 115507414 | 115507436 | + | | 38 | NA | intron (NM_014112, intron 5 of 6) | intron (NM_014112, intron 5 of 6) | 160587 | NM_001282902 | 7227 | Hs.657018 | NM_014112 | ENSG00000104447 | TRPS1 | protein-coding |
| chr11 | 42706767 | 42706789 | - | | 38 | NA | Intergenic | Intergenic | -453088 | NR_038309 | 100507205 | Hs.99310 | NR_038309 | | LINC02740 | ncRNA |
| chr5 | 126726275 | 126726297 | - | | 38 | NA | Intergenic | L1M5\|LINE\|L1 | 50200 | NR_134485 | 102723557 | Hs.582207 | NR_134485 | ENSG00000251072 | LMNB1-DT | ncRNA |
| chr3 | 72793768 | 72793790 | - | | 36 | NA | intron (NM_018130, intron 9 of 10) | intron (NM_018130, intron 9 of 10) | 54668 | NM_018130 | 55164 | Hs.606584 | NM_018130 | ENSG00000144736 | SHQ1 | protein-coding |
| chr3 | 81069593 | 81069615 | - | | 36 | NA | intron (NR_132411, intron 1 of 1) | LTR19-int\|LTR\|ERV1 | 75736 | NR_132411 | 728290 | Hs.563132 | NR_132411 | ENSG00000243694 | LINC02027 | ncRNA |
| chr4 | 185547480 | 185547502 | - | | 36 | NA | Intergenic | Intergenic | -11933 | NM_001114107 | 27295 | Hs.701364 | NM_014476 | ENSG00000154553 | PDLIM3 | protein-coding |
| chr3 | 53825307 | 53825329 | - | | 36 | NA | intron (NM_018397, intron 2 of 8) | L1MC1\|LINE\|L1 | 21075 | NM_018397 | 55349 | Hs.126688 | NM_018397 | ENSG00000016391 | CHDH | protein-coding |
| chr9 | 11802796 | 11802818 | + | | 36 | NA | Intergenic | L1MA8\|LINE\|L1 | -890579 | NM_000550 | 7306 | Hs.270279 | NM_000550 | ENSG00000107165 | TYRP1 | protein-coding |
| chr3 | 53586311 | 53586333 | + | | 34 | NA | intron (NM_001128840, intron 3 of 47) | intron (NM_001128840, intron 3 of 47) | 91273 | NM_001128839 | 776 | Hs.476358 | NM_000720 | ENSG00000157388 | CACNA1D | protein-coding |
| chr6 | 158754201 | 158754223 | + | | 34 | NA | intron (NM_001009991, intron 12 of 16) | Charlie2a\|DNA\|hAT-Charlie | 10541 | NR_037482 | 100500851 | | NR_037482 | ENSG00000265558 | MIR3918 | ncRNA |
| chr7 | 55065252 | 55065274 | + | | 34 | NA | intron (NM_001346941, intron 1 of 21) | intron (NM_001346941, intron 1 of 21) | -44554 | NM_001346900 | 1956 | Hs.488293 | NM_005228 | ENSG00000146648 | EGFR | protein-coding |
| chr6 | 28873313 | 28873335 | + | | 34 | NA | Intergenic | LTR57-int\|LTR\|ERVL | -9647 | NR_033379 | 401242 | Hs.448285 | NR_033379 | ENSG00000225595 | LINC01623 | ncRNA |
| chr21 | 31226802 | 31226824 | - | | 34 | NA | intron (NM_003253, intron 7 of 28) | intron (NM_003253, intron 7 of 28) | -65873 | NM_001353684 | 7074 | Hs.517228 | NM_003253 | ENSG00000156299 | TIAM1 | protein-coding |
| chr18 | 35263657 | 35263679 | - | | 34 | NA | intron (NM_001166012, intron 3 of 4) | intron (NM_001166012, intron 3 of 4) | 22638 | NM_032347 | 84307 | Hs.591061 | NM_032347 | ENSG00000186812 | ZNF397 | protein-coding |
| chr12 | 66796286 | 66796308 | + | | 32 | NA | Intergenic | Intergenic | -117152 | NM_001178074 | 23426 | Hs.505946 | NM_021150 | ENSG00000155974 | GRIP1 | protein-coding |
| chr9 | 134598961 | 134598983 | + | | 32 | NA | Intergenic | Intergenic | -42833 | NM_001278074 | 1289 | Hs.210283 | NM_000093 | ENSG00000130635 | COL5A1 | protein-coding |
| chr2 | 21615898 | 21615920 | - | | 32 | NA | Intergenic | L1MA1\|LINE\|L1 | 94743 | NR_038837 | 645949 | Hs.456782 | NR_038837 | ENSG00000229621 | LINC01822 | ncRNA |
| chr8 | 131937375 | 131937397 | + | | 32 | NA | intron (NM_001323557, intron 1 of 22) | intron (NM_001323557, intron 1 of 22) | 20632 | NM_001323557 | 23167 | Hs.204564 | NM_015137 | ENSG00000132294 | EFR3A | protein-coding |
| chrX | 68635525 | 68635547 | - | | 30 | NA | Intergenic | Intergenic | -12133 | NM_001142503 | 9754 | Hs.95140 | NM_014725 | ENSG00000130052 | STARD8 | protein-coding |
| chr1 | 247437358 | 247437380 | - | | 30 | NA | intron (NM_001079821, intron 8 of 10) | L3\|LINE\|CR1 | 14613 | NM_001004492 | 127623 | Hs.446999 | NM_001004492 | ENSG00000177535 | OR2B11 | protein-coding |
| chr20 | 7078003 | 7078025 | + | | 30 | NA | Intergenic | L1MC3\|LINE\|L1 | 176188 | NR_110609 | 101929265 | Hs.637730 | NR_110609 | ENSG00000228888 | LINC01428 | ncRNA |
| chr13 | 38174447 | 38174469 | - | | 30 | NA | Intergenic | Intergenic | -31226 | NR_047500 | 100874188 | Hs.569249 | NR_047500 | ENSG00000223685 | LINC00571 | ncRNA |
| chr2 | 29807366 | 29807388 | + | | 30 | NA | intron (NM_004304, intron 1 of 28) | intron (NM_004304, intron 1 of 28) | 114234 | NM_004304 | 238 | Hs.654469 | NM_004304 | ENSG00000171094 | ALK | protein-coding |
| chr2 | 150835257 | 150835279 | - | | 30 | NA | Intergenic | HERVL-int\|LTR\|ERVL | 206371 | NR_110247 | 101929282 | Hs.126866 | NR_110244 | ENSG00000224048 | LINC02612 | ncRNA |
| chr4 | 174976343 | 174976365 | + | | 30 | NA | exon (NM_014269, exon 5 of 5) | exon (NM_014269, exon 5 of 5) | 57996 | NM_001278125 | 11086 | Hs.126838 | NM_014269 | ENSG00000168594 | ADAM29 | protein-coding |
| chr13 | 109548171 | 109548193 | - | | 30 | NA | Intergenic | Intergenic | 147486 | NR_126361 | 104326054 | Hs.569298 | NR_126361 | ENSG00000229792 | LINC00399 | ncRNA |
| chr15 | 43444009 | 43444031 | - | | 28 | NA | intron (NM_001141979, intron 14 of 27) | MIRb\|SINE\|MIR | 49151 | NM_001141979 | 7158 | Hs.440968 | NM_005657 | ENSG00000067369 | TP53BP1 | protein-coding |
| chr14 | 85792463 | 85792485 | - | | 28 | NA | Intergenic | L1PA16\|LINE\|L1 | -142204 | NR_110155 | 101928767 | Hs.536380 | NR_110155 | | LINC02328 | ncRNA |
| chr4 | 139360926 | 139360948 | + | | 28 | NA | intron (NM_057175, intron 13 of 19) | intron (NM_057175, intron 13 of 19) | -58386 | NM_001184990 | 4717 | Hs.84549 | NM_002494 | ENSG00000109390 | NDUFC1 | protein-coding |
| chr2 | 230827354 | 230827376 | - | | 28 | NA | Intergenic | MIR\|SINE\|MIR | -37233 | NM_001287241 | 81618 | Hs.111577 | NM_030926 | ENSG00000135916 | ITM2C | protein-coding |
| chr1 | 242574213 | 242574235 | - | | 28 | NA | Intergenic | Intergenic | -49528 | NM_152666 | 200150 | Hs.672452 | NM_152666 | ENSG00000180287 | PLD5 | protein-coding |
| chr17 | 9972299 | 9972320 | - | | 28 | NA | intron (NM_201432, intron 3 of 13) | intron (NM_201432, intron 3 of 13) | -12860 | NM_003644 | 8522 | Hs.462214 | NM_003644 | ENSG00000007237 | GAS7 | protein-coding |
| chr15 | 101112356 | 101112378 | - | | 26 | NA | Intergenic | Intergenic | 139565 | NM_014918 | 22856 | Hs.110488 | NM_014918 | ENSG00000131873 | CHSY1 | protein-coding |
| chr1 | 180570426 | 180570448 | - | | 26 | NA | Intergenic | L1MC4\|LINE\|L1 | 11463 | NR_125716 | 148756 | | NR_125716 | ENSG00000236719 | OVAAL | ncRNA |
| chr1 | 33279984 | 33280006 | + | | 26 | NA | intron (NM_152493, intron 4 of 8) | intron (NM_152493, intron 4 of 8) | 23422 | NM_152493 | 149076 | Hs.743358 | NM_152493 | ENSG00000160094 | ZNF362 | protein-coding |
| chr7 | 113687210 | 113687232 | - | | 26 | NA | Intergenic | Intergenic | 231806 | NM_002711 | 5506 | Hs.458309 | NM_002711 | ENSG00000154415 | PPP1R3A | protein-coding |
| chr6 | 89414238 | 89414260 | - | | 24 | NA | Intergenic | Intergenic | -1973 | NM_021244 | 58528 | Hs.31712 | NM_021244 | ENSG00000025039 | RRAGD | protein-coding |
| chr21 | 34618960 | 34618982 | - | | 24 | NA | Intergenic | Intergenic | -3829 | NM_004414 | 1827 | Hs.282326 | NM_004414 | ENSG00000159200 | RCAN1 | protein-coding |
| chr6 | 102139854 | 102139876 | + | | 24 | NA | Intergenic | MIR3\|SINE\|MIR | 740880 | NM_021956 | 2898 | Hs.98262 | NM_021956 | ENSG00000164418 | GRIK2 | protein-coding |
| chr2 | 15059612 | 15059634 | + | | 24 | NA | Intergenic | Intergenic | 426940 | NR_144632 | 151354 | Hs.260855 | NM_145175 | ENSG00000162981 | LRATD1 | protein-coding |
| chr8 | 57505492 | 57505514 | + | | 24 | NA | Intergenic | MLT1B-int\|LTR\|ERVL-MaLR | 161716 | NR_125818 | 101929488 | Hs.407557 | NR_125818 | ENSG00000253871 | LOC101929488 | ncRNA |
| chr2 | 45031168 | 45031190 | + | | 24 | NA | Intergenic | Intergenic | -21776 | NM_016932 | 10736 | Hs.101937 | NM_016932 | ENSG00000170577 | SIX2 | protein-coding |
| chr12 | 116186478 | 116186500 | - | | 24 | NA | intron (NM_015335, intron 2 of 30) | intron (NM_015335, intron 2 of 30) | -37835 | NR_030351 | 693205 | | NR_030351 | ENSG00000207967 | MIR620 | ncRNA |
| chr1 | 164499061 | 164499083 | - | | 24 | NA | Intergenic | Intergenic | -60288 | NM_001204961 | 5087 | Hs.557097 | NM_002585 | ENSG00000185630 | PBX1 | protein-coding |
| chr2 | 89016903 | 89016925 | - | | 24 | NA | Intergenic | MER70-int\|LTR\|ERVL | 204544 | NR_039635 | 100616399 | | NR_039635 | ENSG00000265510 | MIR4436A | ncRNA |
| chr10 | 103969690 | 103969712 | + | | 24 | NA | intron (NM_014720, intron 1 of 18) | MIRc\|SINE\|MIR | 2516 | NM_014720 | 9748 | Hs.591922 | NM_014720 | ENSG00000065613 | SLK | protein-coding |
| chr6 | 21900999 | 21901020 | - | | 22 | NA | intron (NR_015410, intron 4 of 11) | intron (NR_015410, intron 4 of 11) | 234596 | NR_015410 | 401237 | Hs.712707 | NR_015410 | | CASC15 | ncRNA |
| chr10 | 60492113 | 60492134 | - | | 22 | NA | intron (NM_001204403, intron 2 of 43) | Tigger3a\|DNA\|TcMar-Tigger | 80833 | NM_001204404 | 288 | Hs.499725 | NM_001149 | ENSG00000151150 | ANK3 | protein-coding |
| chr5 | 124588619 | 124588641 | + | | 22 | NA | Intergenic | Intergenic | -150110 | NR_125774 | 103724389 | Hs.582211 | NR_125774 | | LINC01170 | ncRNA |
| chr1 | 231829049 | 231829071 | - | | 22 | NA | intron (NM_001164541, intron 9 of 9) | intron (NM_001164541, intron 9 of 9) | -10543 | NR_002227 | 27184 | Hs.720568 | NR_002227 | | DISC2 | ncRNA |
| chr10 | 15093248 | 15093270 | + | | 20 | NA | Intergenic | Intergenic | -3921 | NM_001097590 | 10557 | Hs.94986 | NM_006414 | ENSG00000152464 | RPP38 | protein-coding |
| chr3 | 5872472 | 5872493 | - | | 20 | NA | Intergenic | Tigger1\|DNA\|TcMar-Tigger | -622227 | NR_039953 | 100616334 | | NR_039953 | ENSG00000265180 | MIR4790 | ncRNA |
| chr7 | 97159243 | 97159265 | - | | 20 | NA | intron (NM_020186, intron 1 of 1) | intron (NM_020186, intron 1 of 1) | 42661 | NM_020186 | 57001 | Hs.592269 | NM_020186 | ENSG00000196636 | SDHAF3 | protein-coding |
| chr11 | 126091225 | 126091247 | - | | 20 | NA | Intergenic | HAL1\|LINE\|L1 | -27944 | NM_001243597 | 50937 | Hs.38034 | NM_016952 | ENSG00000064309 | CDON | protein-coding |
| chr8 | 41053555 | 41053577 | - | | 20 | NA | Intergenic | Intergenic | -155742 | NM_001135731 | 79698 | Hs.591850 | NM_024645 | ENSG00000165061 | ZMAT4 | protein-coding |
| chr11 | 123426792 | 123426814 | - | | 20 | NA | Intergenic | Intergenic | -3660 | NM_001354837 | 57476 | Hs.144725 | NM_020716 | ENSG00000023171 | GRAMD1B | protein-coding |
| chr10 | 89858160 | 89858182 | + | | 20 | NA | intron (NR_149080, intron 1 of 1) | intron (NR_149080, intron 1 of 1) | 4265 | NR_149080 | 105378425 | Hs.559891 | NR_149079 | ENSG00000280560 | LINC01374 | ncRNA |
| chr2 | 222303116 | 222303138 | + | | 18 | NA | intron (NM_153038, intron 1 of 1) | intron (NM_153038, intron 1 of 1) | -4131 | NM_181458 | 5077 | Hs.42146 | NM_000438 | ENSG00000135903 | PAX3 | protein-coding |
| chr10 | 102445447 | 102445469 | + | | 18 | NA | Intergenic | L2b\|LINE\|L2 | -4359 | NR_038938 | 100505761 | Hs.656457 | NR_038937 | ENSG00000269609 | RPARP-AS1 | ncRNA |
| chr17 | 12147557 | 12147579 | - | | 18 | NA | Intergenic | MIRb\|SINE\|MIR | 65669 | NR_030613 | 100126313 | | NR_030613 | ENSG00000266297 | MIR744 | ncRNA |
| chr6 | 109238197 | 109238219 | + | | 18 | NA | intron (NR_152435, intron 14 of 45) | intron (NR_152435, intron 14 of 45) | 72377 | NR_152435 | 221262 | Hs.373914 | NM_173671 | ENSG00000203799 | CCDC162P | pseudo |
| chr2 | 95724166 | 95724188 | + | | 18 | NA | Intergenic | Intergenic | 102804 | NR_103734 | 150759 | Hs.503463 | NR_103734 | | LINC00342 | ncRNA |
| chr3 | 123057239 | 123057261 | - | | 18 | NA | Intergenic | Intergenic | -9759 | NM_006810 | 10954 | Hs.477352 | NM_006810 | ENSG00000065485 | PDIA5 | protein-coding |
| chr11 | 68589937 | 68589959 | - | | 18 | NA | intron (NM_001164160, intron 16 of 24) | intron (NM_001164160, intron 16 of 24) | -94527 | NM_015973 | 51083 | Hs.278959 | NM_015973 | ENSG00000069482 | GAL | protein-coding |
| chr9 | 83186253 | 83186275 | - | | 18 | NA | Intergenic | Intergenic | 81023 | NM_001244962 | 257019 | Hs.127535 | NM_174938 | ENSG00000172159 | FRMD3 | protein-coding |
| chr2 | 75531194 | 75531216 | - | | 18 | NA | intron (NR_110281, intron 1 of 2) | intron (NR_110281, intron 1 of 2) | 7137 | NR_110281 | 101927884 | Hs.671110 | NR_110281 | ENSG00000231172 | LOC101927884 | ncRNA |
| chr10 | 50885790 | 50885812 | - | | 18 | NA | promoter-TSS (NM_001198819) | promoter-TSS (NM_001198819) | -126 | NM_001198820 | 29974 | Hs.282795 | NM_014576 | ENSG00000148584 | A1CF | protein-coding |
| chr14 | 21536563 | 21536585 | + | | 18 | NA | intron (NM_005407, intron 1 of 1) | intron (NM_005407, intron 1 of 1) | 642 | NM_005407 | 6297 | Hs.416358 | NM_005407 | ENSG00000165821 | SALL2 | protein-coding |
| chr1 | 45948370 | 45948392 | - | | 18 | NA | intron (NM_001324321, intron 3 of 28) | L1PREC2\|LINE\|L1 | 84033 | NM_001324321 | 23139 | Hs.319481 | NM_015112 | ENSG00000086015 | MAST2 | protein-coding |
| chr13 | 106686685 | 106686707 | + | | 18 | NA | Intergenic | Intergenic | 32816 | NR_047026 | 100874173 | Hs.569297 | NR_047026 | ENSG00000230156 | LINC00443 | ncRNA |
| chr17 | 59941512 | 59941534 | + | | 18 | NA | intron (NM_001272044, intron 14 of 15) | AluJb\|SINE\|Alu | 23233 | NM_016125 | 51136 | Hs.531701 | NM_016125 | ENSG00000189050 | RNFT1 | protein-coding |

| | | | | | | | | | | | | | | | | |
|---|---|---|---|---|---|---|---|---|---|---|---|---|---|---|---|---|
| chr19 | 13380324 | 13380346 | - | | 18 | NA | intron (NM_001127222, intron 3 of 46) | MER33\|DNA\|hAT-Charlie | 126125 | NM_000068 | 773 | Hs.501632 | NM_000068 | ENSG00000141837 | CACNA1A | protein-coding |
| chr10 | 112729797 | 112729819 | - | | 18 | NA | intron (NM_001318205, intron 7 of 7) | intron (NM_001318205, intron 7 of 7) | -93688 | NR_120684 | 103344931 | Hs.458447 | NR_120684 | ENSG00000260917 | LOC103344931 | ncRNA |
| chr3 | 176716332 | 176716354 | + | | 18 | NA | Intergenic | Intergenic | -80811 | NR_109968 | 100505547 | Hs.253350 | NR_109968 | | LINC01208 | ncRNA |
| chr2 | 223696211 | 223696233 | - | | 16 | NA | Intergenic | Intergenic | -93723 | NM_003469 | 7857 | Hs.516726 | NM_003469 | ENSG00000171951 | SCG2 | protein-coding |
| chr7 | 94342746 | 94342768 | + | | 16 | NA | Intergenic | Intergenic | 49435 | NR_147206 | 101927525 | Hs.571263 | NR_147206 | | COL1A2-AS1 | ncRNA |
| chr7 | 38755649 | 38755671 | + | | 16 | NA | intron (NM_014396, intron 19 of 28) | MIR3\|SINE\|MIR | -68571 | NR_028347 | 340286 | Hs.144075 | NM_001105282 | ENSG00000164556 | FAM183BP | pseudo |
| chr10 | 131883770 | 131883792 | - | | 16 | NA | Intergenic | Intergenic | -16863 | NM_018461 | 55844 | Hs.380372 | NM_018461 | ENSG00000175470 | PPP2R2D | protein-coding |
| chr8 | 137439771 | 137439793 | - | | 16 | NA | Intergenic | L1MEc\|LINE\|L1 | -26194 | NR_125428 | 101927915 | Hs.695851 | NR_125428 | | LOC101927915 | ncRNA |
| chr10 | 120657743 | 120657765 | + | | 16 | NA | Intergenic | Intergenic | 59805 | NR_103717 | 404216 | Hs.196578 | NM_001012711 | ENSG00000177234 | LINC01561 | ncRNA |
| chr16 | 51002340 | 51002361 | - | | 16 | NA | intron (NR_039672, intron 2 of 6) | Tigger3a\|DNA\|TcMar-Tigger | -15408 | NR_110913 | 101927334 | Hs.555184 | NR_110913 | ENSG00000261637 | LINC02127 | ncRNA |
| chr6 | 70256953 | 70256974 | - | | 16 | NA | intron (NM_001851, intron 20 of 37) | intron (NM_001851, intron 20 of 37) | 26245 | NM_078485 | 1297 | Hs.590892 | NM_001851 | ENSG00000112280 | COL9A1 | protein-coding |
| chr3 | 115945315 | 115945338 | - | | 16 | NA | intron (NM_001318915, intron 3 of 8) | intron (NM_001318915, intron 3 of 8) | -40200 | NR_145779 | 109623450 | | NR_145779 | | SNORD155 | snoRNA |
| chr8 | 2591790 | 2591812 | - | | 16 | NA | intron (NR_125423, intron 4 of 4) | intron (NR_125423, intron 4 of 4) | 31564 | NR_125423 | 101927815 | Hs.571467 | NR_125423 | ENSG00000254319 | LOC101927815 | ncRNA |
| chr6 | 82833813 | 82833835 | - | | 14 | NA | Intergenic | MLT1A0-int\|LTR\|ERVL-MaLR | 232017 | NM_001350604 | 90025 | Hs.148609 | NM_198920 | ENSG00000118420 | UBE3D | protein-coding |
| chr3 | 4442576 | 4442598 | - | | 14 | NA | intron (NM_001164674, intron 2 of 7) | HAL1M8\|LINE\|L1 | 24695 | NM_001164674 | 285362 | Hs.350475 | NM_182760 | ENSG00000144455 | SUMF1 | protein-coding |
| chr14 | 56246090 | 56246112 | - | | 14 | NA | intron (NM_021255, intron 2 of 5) | L3\|LINE\|CR1 | 127726 | NM_021255 | 57161 | Hs.657926 | NM_021255 | ENSG00000139946 | PELI2 | protein-coding |
| chr7 | 16050467 | 16050489 | - | | 14 | NA | Intergenic | Intergenic | -160008 | NR_038947 | 100506025 | Hs.649029 | NR_038946 | ENSG00000229688 | CRPPA-AS1 | ncRNA |
| chr12 | 128946087 | 128946109 | + | | 14 | NA | intron (NR_133646, intron 8 of 10) | intron (NR_133646, intron 8 of 10) | 92664 | NR_133646 | 144423 | Hs.655668 | NM_144669 | ENSG00000151948 | GLT1D1 | protein-coding |
| chr11 | 120321962 | 120321984 | + | | 14 | NA | Intergenic | Intergenic | -3156 | NM_001198670 | 219902 | Hs.643516 | NM_174926 | ENSG00000181264 | TLCD5 | protein-coding |
| chr2 | 159765068 | 159765090 | + | | 14 | NA | intron (NM_001282805, intron 11 of 11) | intron (NM_001282805, intron 11 of 11) | 31188 | NM_001282807 | 64844 | Hs.529272 | NM_022826 | ENSG00000136536 | 7-Mar | protein-coding |
| chr18 | 33086493 | 33086515 | - | | 14 | NA | intron (NM_001308126, intron 20 of 22) | MER103C\|DNA\|hAT-Charlie | -313493 | NM_020805 | 57565 | Hs.446164 | NM_020805 | ENSG00000197705 | KLHL14 | protein-coding |
| chr4 | 4896409 | 4896431 | - | | 14 | NA | Intergenic | MLT1D\|LTR\|ERVL-MaLR | -24388 | NR_125893 | 101928306 | Hs.543743 | NR_125893 | | LOC101928306 | ncRNA |
| chr1 | 208628929 | 208628951 | - | | 14 | NA | Intergenic | Intergenic | -99748 | NR_146914 | 105372893 | Hs.568720 | NR_146914 | ENSG00000232812 | LINC01717 | ncRNA |
| chr10 | 82485501 | 82485523 | - | | 14 | NA | intron (NM_001010848, intron 2 of 8) | intron (NM_001010848, intron 2 of 8) | -252592 | NR_120666 | 101929590 | Hs.733268 | NR_120666 | ENSG00000225738 | NRG3-AS1 | ncRNA |
| chr4 | 58939438 | 58939460 | - | | 14 | NA | Intergenic | Intergenic | -44833 | NR_133941 | 105377247 | Hs.298228 | NR_133941 | ENSG00000251266 | LINC02429 | ncRNA |
| chr14 | 21107333 | 21107355 | - | | 12 | NA | Intergenic | Intergenic | -2640 | NM_001102454 | 51222 | Hs.250493 | NM_016423 | ENSG00000165804 | ZNF219 | protein-coding |
| chr10 | 53813505 | 53813527 | + | | 12 | NA | intron (NM_001142770, intron 32 of 34) | L1ME3A\|LINE\|L1 | -672714 | NR_134503 | 105378311 | Hs.623892 | NR_134503 | ENSG00000234173 | LOC105378311 | ncRNA |
| chr3 | 79973032 | 79973053 | + | | 12 | NA | Intergenic | Intergenic | -205133 | NM_002941 | 6091 | Hs.744218 | NM_002941 | ENSG00000169855 | ROBO1 | protein-coding |
| chrX | 101639755 | 101639777 | - | | 12 | NA | Intergenic | Intergenic | 16405 | NM_177948 | 51566 | Hs.592225 | NM_016607 | ENSG00000102401 | ARMCX3 | protein-coding |
| chr1 | 77293426 | 77293448 | + | | 12 | NA | intron (NM_012093, intron 2 of 13) | intron (NM_012093, intron 2 of 13) | 10835 | NM_012093 | 26289 | Hs.559718 | NM_012093 | ENSG00000154027 | AK5 | protein-coding |
| chr6 | 153989584 | 153989606 | + | | 12 | NA | Intergenic | Intergenic | -20901 | NM_001145279 | 4988 | Hs.2353 | NM_000914 | ENSG00000112038 | OPRM1 | protein-coding |
| chr14 | 73338732 | 73338754 | + | | 12 | NA | intron (NM_001005743, intron 4 of 12) | MER20B\|DNA\|hAT-Charlie | -92742 | NR_135248 | 101928123 | Hs.660593 | NR_135248 | | LOC101928123 | ncRNA |
| chr2 | 90183723 | 90183745 | + | | 12 | NA | Intergenic | MER70-int\|LTR\|ERVL | -545827 | NR_136329 | 101927050 | Hs.578075 | NR_136329 | | LOC101927050 | pseudo |
| chr8 | 98346631 | 98346653 | + | | 12 | NA | Intergenic | L1MC1\|LINE\|L1 | -52249 | NR_135747 | 79815 | Hs.309489 | NM_024759 | ENSG00000104361 | NIPAL2 | protein-coding |
| chr2 | 138689550 | 138689572 | - | | 12 | NA | intron (NM_007226, intron 1 of 1) | intron (NM_007226, intron 1 of 1) | 90680 | NM_007226 | 11249 | Hs.435019 | NM_007226 | ENSG00000144227 | NXPH2 | protein-coding |
| chr9 | 7245101 | 7245123 | - | | 12 | NA | Intergenic | Intergenic | 487762 | NR_148679 | 23081 | Hs.709425 | NM_015061 | ENSG00000107077 | KDM4C | protein-coding |
| chr5 | 30348979 | 30349001 | - | | 12 | NA | Intergenic | Intergenic | 468430 | NR_134264 | 105374704 | Hs.151157 | NR_134264 | | LOC105374704 | ncRNA |
| chr2 | 210094865 | 210094887 | - | | 12 | NA | intron (NM_152519, intron 5 of 14) | intron (NM_152519, intron 5 of 14) | 64954 | NR_110291 | 101928020 | Hs.635556 | NR_110291 | | LOC101928020 | ncRNA |
| chr8 | 115042055 | 115042077 | - | | 12 | NA | Intergenic | L2c\|LINE\|L2 | 625946 | NM_001282902 | 7227 | Hs.657018 | NM_014112 | ENSG00000104447 | TRPS1 | protein-coding |
| chr14 | 70357125 | 70357147 | + | | 10 | NA | intron (NM_001202548, intron 3 of 5) | L1M4\|LINE\|L1 | 2595 | NM_001204090 | 51241 | Hs.709581 | NM_016468 | ENSG00000133983 | COX16 | protein-coding |
| chr6 | 98050836 | 98050858 | + | | 10 | NA | Intergenic | Intergenic | 26316 | NR_031579 | 100302164 | | NR_031579 | ENSG00000238367 | MIR2113 | ncRNA |
| chr12 | 21531681 | 21531703 | - | | 10 | NA | non-coding (NR_135188, exon 3 of 3) | non-coding (NR_135188, exon 3 of 3) | 3767 | NR_135188 | 80763 | Hs.130692 | NM_030572 | ENSG00000134548 | SPX | protein-coding |
| chr15 | 97169283 | 97169305 | - | | 10 | NA | Intergenic | Intergenic | 352517 | NR_120324 | 101927286 | Hs.611279 | NR_120324 | ENSG00000259664 | LINC02254 | ncRNA |
| chr2 | 102598782 | 102598804 | - | | 10 | NA | Intergenic | L1ME2\|LINE\|L1 | -20741 | NM_003048 | 6549 | Hs.250083 | NM_003048 | ENSG00000115616 | SLC9A2 | protein-coding |
| chrX | 103513427 | 103513449 | - | | 10 | NA | intron (NM_080879, intron 2 of 2) | intron (NM_080879, intron 2 of 2) | 6051 | NM_080879 | 142684 | Hs.706904 | NM_080879 | ENSG00000172476 | RAB40A | protein-coding |
| chr7 | 101641374 | 101641396 | - | | 10 | NA | Intergenic | MLT1I\|LTR\|ERVL-MaLR | -12089 | NM_138403 | 93408 | Hs.247831 | NM_138403 | ENSG00000106436 | MYL10 | protein-coding |
| chr8 | 81839718 | 81839739 | + | | 10 | NA | exon (NM_152837, exon 2 of 7) | exon (NM_152837, exon 2 of 7) | 2558 | NM_152836 | 64089 | Hs.492121 | NM_022133 | ENSG00000104497 | SNX16 | protein-coding |
| chr17 | 38460492 | 38460514 | - | | 10 | NA | intron (NM_001199417, intron 2 of 23).2 | intron (NM_001199417, intron 2 of 23).2 | 32027 | NM_001199417 | 57636 | Hs.374446 | NM_020876 | ENSG00000275832 | ARHGAP23 | protein-coding |
| chr7 | 14450639 | 14450661 | - | | 10 | NA | intron (NM_145695, intron 20 of 23) | intron (NM_145695, intron 20 of 23) | 390800 | NM_145695 | 1607 | Hs.567255 | NM_004080 | ENSG00000136267 | DGKB | protein-coding |
| chr15 | 101759732 | 101759755 | - | | 10 | NA | Intergenic | CpG | -12873 | NM_001321551 | 100128108 | Hs.104864 | NM_001321551 | | LOC100128108 | protein-coding |
| chr15 | 101753207 | 101753230 | - | | 10 | NA | Intergenic | CpG | -6348 | NM_001321551 | 100128108 | Hs.104864 | NM_001321551 | | LOC100128108 | protein-coding |
| chr17 | 21743687 | 21743709 | + | | 10 | NA | Intergenic | Intergenic | 51175 | NM_001194958 | 100134444 | Hs.200629 | NM_001194958 | ENSG00000260458 | KCNJ18 | protein-coding |
| chr10 | 133028226 | 133028248 | + | | 10 | NA | Intergenic | Intergenic | 58460 | NR_111905 | 100128127 | Hs.694139 | NR_111905 | | ADGRA1-AS1 | ncRNA |
| chr3 | 141594994 | 141595016 | + | | 10 | NA | intron (NM_001303246, intron 20 of 24) | L1MB8\|LINE\|L1 | -68194 | NR_136190 | 646730 | Hs.570677 | NR_136190 | | LINC02618 | ncRNA |
| chr9 | 8729609 | 8729631 | - | | 10 | NA | intron (NM_001040712, intron 1 of 29) | MIR\|SINE\|MIR | 4326 | NM_130393 | 5789 | Hs.446083 | NM_002839 | ENSG00000153707 | PTPRD | protein-coding |
| chr17 | 52416837 | 52416859 | + | | 10 | NA | intron (NR_146898, intron 1 of 3) | intron (NR_146898, intron 1 of 3) | 26326 | NR_146898 | 105371830 | | NR_146898 | ENSG00000263317 | LINC01982 | ncRNA |
| chrX | 104707815 | 104707837 | + | | 8 | NA | intron (NM_017416, intron 2 of 10) | intron (NM_017416, intron 2 of 10) | 141511 | NM_017416 | 26280 | Hs.675519 | NM_017416 | ENSG00000189108 | IL1RAPL2 | protein-coding |
| chr11 | 88210326 | 88210348 | - | | 8 | NA | Intergenic | MIR\|SINE\|MIR | 33835 | NR_036124 | 100423040 | | NR_036124 | ENSG00000266581 | MIR3166 | ncRNA |
| chr16 | 10700074 | 10700096 | - | | 8 | NA | Intergenic | SATR2\|Satellite\|Satellite | -5140 | NM_144674 | 146279 | Hs.143519 | NM_144674 | ENSG00000153060 | TEKT5 | protein-coding |
| chr10 | 128449123 | 128449145 | + | | 8 | NA | Intergenic | L1ME3Cz\|LINE\|L1 | 163184 | NR_120619 | 101927381 | Hs.523342 | NR_120619 | ENSG00000280953 | LINC01163 | ncRNA |
| chr4 | 85789156 | 85789178 | + | | 8 | NA | intron (NM_001042669, intron 1 of 7) | MER21B\|LTR\|ERVL | 10469 | NM_001042669 | 83478 | Hs.444229 | NM_031305 | ENSG00000138639 | ARHGAP24 | protein-coding |
| chr8 | 15891661 | 15891683 | - | | 8 | NA | Intergenic | Intergenic | 301119 | NM_138715 | 4481 | Hs.147635 | NM_002445 | ENSG00000038945 | MSR1 | protein-coding |
| chr3 | 55351904 | 55351926 | - | | 8 | NA | Intergenic | Intergenic | 129483 | NM_001256105 | 7474 | Hs.643085 | NM_003392 | ENSG00000114251 | WNT5A | protein-coding |
| chr10 | 124206314 | 124206336 | - | | 8 | NA | Intergenic | Intergenic | -112718 | NM_001270765 | 51363 | Hs.287537 | NM_014863 | ENSG00000182022 | CHST15 | protein-coding |
| chr7 | 149530259 | 149530281 | - | | 8 | NA | Intergenic | LTR25-int\|LTR\|ERV1 | -32463 | NM_001163474 | 155061 | Hs.24643 | NM_152557 | ENSG00000181220 | ZNF746 | protein-coding |
| chr6 | 39439539 | 39439561 | - | | 8 | NA | intron (NM_001289021, intron 12 of 21) | intron (NM_001289021, intron 12 of 21) | -8176 | NM_001351566 | 221458 | Hs.588202 | NM_145027 | ENSG00000164627 | KIF6 | protein-coding |
| chrX | 145095175 | 145095197 | - | | 8 | NA | Intergenic | Intergenic | -152401 | NM_001009614 | 494118 | | NM_001009614 | ENSG00000203923 | SPANXN1 | protein-coding |
| chr22 | 33076347 | 33076369 | + | | 8 | NA | Intergenic | Intergenic | -17967 | NM_001135774 | 8224 | Hs.608750 | NM_003490 | ENSG00000185666 | SYN3 | protein-coding |
| chr8 | 20857535 | 20857557 | - | | 8 | NA | Intergenic | L1M4\|LINE\|L1 | -96087 | NR_130773 | 102467222 | Hs.444226 | NR_130773 | ENSG00000253147 | LOC102467222 | ncRNA |
| chr9 | 85291897 | 85291919 | + | | 8 | NA | Intergenic | Intergenic | 450121 | NM_001330701 | 23287 | Hs.719980 | NM_015239 | ENSG00000135049 | AGTPBP1 | protein-coding |
| chr19 | 6479626 | 6479648 | + | | 8 | NA | intron (NM_024898, intron 4 of 22) | intron (NM_024898, intron 4 of 22) | 2171 | NM_001290331 | 79958 | Hs.236449 | NM_024898 | ENSG00000205744 | DENND1C | protein-coding |
| chr14 | 77048263 | 77048285 | - | | 6 | NA | intron (NR_110554, intron 1 of 3) | intron (NR_110554, intron 1 of 3) | 7225 | NR_110554 | 283575 | Hs.651736 | NR_110554 | ENSG00000246548 | LINC02288 | ncRNA |
| chr4 | 188178661 | 188178683 | - | | 6 | NA | Intergenic | Intergenic | 39231 | NM_178556 | 339976 | Hs.348618 | NM_178556 | ENSG00000184108 | TRIML1 | protein-coding |
| chr1 | 14541814 | 14541836 | + | | 6 | NA | Intergenic | Intergenic | -56892 | NM_015209 | 23254 | Hs.368823 | NM_015209 | ENSG00000189337 | KAZN | protein-coding |
| chr10 | 95838283 | 95838305 | + | | 6 | NA | intron (NM_001164183, intron 2 of 7) | L1ME3\|LINE\|L1 | -69671 | NM_001130446 | 387707 | Hs.652240 | NM_001001732 | ENSG00000188649 | CC2D2B | protein-coding |
| chr4 | 61639973 | 61639995 | + | | 6 | NA | intron (NM_001322246, intron 4 of 25) | intron (NM_001322246, intron 4 of 25) | 438728 | NM_001322246 | 23284 | Hs.28391 | NM_015236 | ENSG00000150471 | ADGRL3 | protein-coding |
| chr14 | 77356327 | 77356349 | + | | 6 | NA | intron (NM_001346131, intron 1 of 5) | intron (NM_001346131, intron 1 of 5) | 17571 | NM_001346134 | 283578 | Hs.26403 | NM_213601 | ENSG00000100580 | TMED8 | protein-coding |
| chr5 | 39385753 | 39385775 | - | | 6 | NA | intron (NM_001244871, intron 8 of 13) | intron (NM_001244871, intron 8 of 13) | -21204 | NM_001737 | 735 | Hs.654443 | NM_001737 | ENSG00000113600 | C9 | protein-coding |
| chr3 | 11734281 | 11734303 | + | | 6 | NA | Intergenic | L2a\|LINE\|L2 | -13524 | NM_014667 | 9686 | Hs.740389 | NM_014667 | ENSG00000144560 | VGLL4 | protein-coding |
| chr20 | 26098176 | 26098198 | - | | 6 | NA | Intergenic | Intergenic | 15854 | NR_003678 | 149934 | Hs.711274 | NM_001039379 | ENSG00000240108 | NCOR1P1 | pseudo |
| chr3 | 139023456 | 139023478 | + | | 6 | NA | Intergenic | L1M5\|LINE\|L1 | -2541 | NM_001013650 | 389151 | Hs.531376 | NM_001013650 | ENSG00000184814 | PRR23B | protein-coding |
| chr9 | 91409334 | 91409356 | - | | 6 | NA | 3' UTR (NM_005384, exon 2 of 2) | 3' UTR (NM_005384, exon 2 of 2) | 14517 | NM_005384 | 4783 | Hs.79334 | NM_005384 | ENSG00000165030 | NFIL3 | protein-coding |
| chr11 | 61739029 | 61739051 | + | | 6 | NA | intron (NM_006133, intron 16 of 19) | intron (NM_006133, intron 16 of 19) | -13597 | NM_001127392 | 745 | Hs.473109 | NM_013279 | ENSG00000124920 | MYRF | protein-coding |
| chr3 | 42744287 | 42744309 | + | | 6 | NA | intron (NR_046732, intron 3 of 3) | intron (NR_046732, intron 3 of 3) | 11723 | NR_046732 | 100874114 | Hs.669654 | NR_046732 | ENSG00000173811 | CCDC13-AS1 | ncRNA |
| chr15 | 62404707 | 62404729 | + | | 6 | NA | Intergenic | Intergenic | 61689 | NR_106733 | 102464834 | | NR_106733 | ENSG00000275451 | MIR6085 | ncRNA |
| chr15 | 101521204 | 101521226 | + | | 6 | NA | Intergenic | Intergenic | -31231 | NM_002570 | 5046 | Hs.498494 | NM_002570 | ENSG00000140479 | PCSK6 | protein-coding |
| chr1 | 163698112 | 163698134 | - | | 6 | NA | Intergenic | Intergenic | -274932 | NR_104294 | 100422212 | Hs.556898 | NR_104294 | | LOC100422212 | pseudo |
| chr14 | 47259135 | 47259157 | - | | 6 | NA | intron (NM_001113498, intron 2 of 16) | Charlie2a\|DNA\|hAT-Charlie | 84100 | NR_103766 | 161357 | Hs.436380 | NM_182830 | ENSG00000139915 | MDGA2 | protein-coding |
| chr7 | 109760082 | 109760104 | + | | 6 | NA | Intergenic | L1PA15\|LINE\|L1 | 200120 | NR_003024 | 442720 | Hs.530096 | NR_003024 | ENSG00000237064 | EIF3IP1 | pseudo |
| chr9 | 34520783 | 34520805 | + | | 6 | NA | TTS (NM_198573) | TTS (NM_198573) | 2245 | NM_198573 | 375704 | Hs.522085 | NM_198573 | ENSG00000168913 | ENHO | protein-coding |
| chr3 | 110919086 | 110919108 | - | | 6 | NA | Intergenic | Intergenic | 150862 | NR_045114 | 100506555 | Hs.535689 | NR_045114 | | NECTIN3-AS1 | ncRNA |
| chr2 | 204644667 | 204644689 | + | | 6 | NA | intron (NM_205863, intron 1 of 21) | intron (NM_205863, intron 1 of 21) | 98885 | NM_205863 | 117583 | Hs.657382 | NM_057177 | ENSG00000116117 | PARD3B | protein-coding |
| chr8 | 63936549 | 63936571 | + | | 6 | NA | intron (NR_125826, intron 4 of 4) | L1MD2\|LINE\|L1 | 167130 | NR_038875 | 286184 | Hs.586402 | NR_038875 | ENSG00000253734 | LINC01289 | ncRNA |
| chr11 | 27699822 | 27699844 | + | | 6 | NA | promoter-TSS (NM_170734) | promoter-TSS (NM_170734) | -166 | NM_170734 | 627 | Hs.502182 | NM_001709 | ENSG00000176697 | BDNF | protein-coding |
| chr3 | 102455503 | 102455525 | + | | 6 | NA | intron (NM_001329788, intron 4 of 11) | intron (NM_001329788, intron 4 of 11) | 20499 | NM_001329788 | 131368 | Hs.352213 | NM_175056 | ENSG00000170044 | ZPLD1 | protein-coding |
| chr13 | 80666067 | 80666089 | - | | 4 | NA | Intergenic | L1MDa\|LINE\|L1 | -324963 | NM_005842 | 10253 | Hs.18676 | NM_005842 | ENSG00000136158 | SPRY2 | protein-coding |
| chr2 | 54015785 | 54015807 | - | | 4 | NA | intron (NM_001320587, intron 2 of 5) | intron (NM_001320587, intron 2 of 5) | 44693 | NM_001320587 | 98 | Hs.516173 | NM_138448 | ENSG00000170634 | ACYP2 | protein-coding |
| chr6 | 57484981 | 57485003 | - | | 4 | NA | intron (NM_000947, intron 7 of 13) | L2a\|LINE\|L2 | 94860 | NR_036097 | 100422884 | | NR_036097 | ENSG00000212017 | MIR548U | ncRNA |
| chr2 | 11212731 | 11212753 | - | | 4 | NA | intron (NM_001321643, intron 17 of 32) | intron (NM_001321643, intron 17 of 32) | 57370 | NM_001282712 | 130814 | Hs.274415 | NM_152391 | ENSG00000162976 | SLC66A3 | protein-coding |
| chr16 | 52285428 | 52285450 | - | | 4 | NA | Intergenic | Intergenic | 26875 | NR_135281 | 283854 | Hs.266768 | NR_135281 | ENSG00000260887 | CASC22 | ncRNA |
| chr6 | 36595059 | 36595080 | + | | 4 | NA | intron (NM_003017, intron 1 of 5) | intron (NM_003017, intron 1 of 5) | 756 | NM_003017 | 6428 | Hs.405144 | NM_003017 | ENSG00000112081 | SRSF3 | protein-coding |
| chr15 | 63701620 | 63701642 | - | | 4 | NA | intron (NM_003922, intron 25 of 77) | MER58A\|DNA\|hAT-Charlie | -100804 | NR_034080 | 100130855 | Hs.631163 | NR_034080 | ENSG00000259248 | USP3-AS1 | ncRNA |
| chr12 | 131084759 | 131084781 | + | | 4 | NA | intron (NM_198827, intron 15 of 24) | intron (NM_198827, intron 15 of 24) | -80241 | NR_026670 | 116437 | Hs.675497 | NR_026670 | ENSG00000204603 | LINC01257 | ncRNA |
| chr2 | 152859799 | 152859821 | + | | 4 | NA | Intergenic | Intergenic | 141173 | NR_146431 | 151188 | Hs.643580 | NM_152522 | ENSG00000177917 | ARL6IP6 | protein-coding |
| chr3 | 1521277 | 1521299 | - | | 4 | NA | Intergenic | Intergenic | 428630 | NM_001349361 | 27255 | Hs.387300 | NM_014461 | ENSG00000134115 | CNTN6 | protein-coding |
| chr16 | 71374193 | 71374215 | + | | 4 | NA | intron (NR_027910, intron 2 of 9) | intron (NR_027910, intron 2 of 9) | 15491 | NM_007088 | 794 | Hs.106857 | NM_001740 | ENSG00000172137 | CALB2 | protein-coding |
| chr13 | 106820578 | 106820600 | - | | 4 | NA | Intergenic | Intergenic | 166709 | NR_047026 | 100874173 | Hs.569297 | NR_047026 | ENSG00000230156 | LINC00443 | ncRNA |
| chr1 | 168484662 | 168484684 | + | | 4 | NA | intron (NR_125956, intron 4 of 4) | intron (NR_125956, intron 4 of 4) | 10971 | NR_125956 | 101928565 | Hs.563806 | NR_125956 | ENSG00000228697 | LOC101928565 | ncRNA |
| chr9 | 8803427 | 8803449 | - | | 4 | NA | intron (NM_002839, intron 11 of 45) | intron (NM_002839, intron 11 of 45) | -54580 | NR_121599 | 101929407 | Hs.649971 | NR_121599 | | PTPRD-AS1 | ncRNA |
| chr6 | 165642962 | 165642984 | - | | 4 | NA | intron (NM_006661, intron 1 of 22) | Charlie2a\|DNA\|hAT-Charlie | 19127 | NM_001130690 | 10846 | Hs.348762 | NM_006661 | ENSG00000112541 | PDE10A | protein-coding |
| chr13 | 89476501 | 89476523 | - | | 4 | NA | Intergenic | L1PA13\|LINE\|L1 | 23997 | NR_131228 | 105370308 | Hs.599406 | NR_131228 | ENSG00000226037 | LINC01040 | ncRNA |
| chr10 | 28126640 | 28126662 | + | | 4 | NA | intron (NM_001318170, intron 6 of 16) | intron (NM_001318170, intron 6 of 16) | -53273 | NR_132973 | 106635548 | | NR_132973 | | SNORD130 | snoRNA |
| chr6 | 65787998 | 65788020 | + | | 4 | NA | non-coding (NR_026540, exon 1 of 1) | non-coding (NR_026540, exon 1 of 1) | 130 | NR_026540 | 442229 | Hs.390414 | NR_026540 | | SLC25A51P1 | pseudo |
| chr2 | 235601979 | 235602001 | + | | 4 | NA | intron (NM_001037131, intron 1 of 17) | L2a\|LINE\|L2 | -84069 | NR_146582 | 27321 | | NR_146582 | | TNRC17 | ncRNA |
| chr9 | 118060221 | 118060243 | - | | 4 | NA | Intergenic | Intergenic | 356057 | NM_138557 | 7099 | Hs.174312 | NM_003266 | ENSG00000136869 | TLR4 | protein-coding |
| chr9 | 35367627 | 35367649 | + | | 4 | NA | intron (NM_001330653, intron 11 of 39) | intron (NM_001330653, intron 11 of 39) | -39117 | NR_110320 | 158381 | Hs.575661 | NR_003581 | ENSG00000179766 | ATP8B5P | pseudo |
| chr22 | 32196407 | 32196429 | + | | 4 | NA | intron (NM_001098527, intron 2 of 4) | intron (NM_001098527, intron 2 of 4) | -2939 | NM_006605 | 10739 | Hs.157427 | NM_006605 | ENSG00000128253 | RFPL2 | protein-coding |
| chr6 | 97777821 | 97777843 | + | | 4 | NA | Intergenic | L1M7\|LINE\|L1 | -246699 | NR_031579 | 100302164 | | NR_031579 | ENSG00000238367 | MIR2113 | ncRNA |
| chr3 | 180221523 | 180221545 | - | | 4 | NA | Intergenic | L1M3\|LINE\|L1 | -184481 | NM_001349401 | 51555 | Hs.235331 | NM_016559 | ENSG00000114757 | PEX5L | protein-coding |
| chr8 | 31346768 | 31346790 | + | | 4 | NA | Intergenic | Intergenic | -292973 | NM_013962 | 3084 | Hs.453951 | NM_004495 | ENSG00000157168 | NRG1 | protein-coding |
| chr8 | 119666370 | 119666392 | + | | 4 | NA | intron (NM_001330600, intron 1 of 25) | intron (NM_001330600, intron 1 of 25) | 7195 | NM_001330600 | 5168 | Hs.190977 | NM_006209 | ENSG00000136960 | ENPP2 | protein-coding |
| chr13 | 110309014 | 110309036 | + | | 4 | NA | intron (NM_001846, intron 3 of 47) | intron (NM_001846, intron 3 of 47) | 1758 | NM_001846 | 1284 | Hs.508716 | NM_001846 | ENSG00000134871 | COL4A2 | protein-coding |
| chr12 | 98630861 | 98630883 | + | | 4 | NA | intron (NM_153687, intron 2 of 2) | intron (NM_153687, intron 2 of 2) | 14179 | NM_153687 | 121457 | Hs.252543 | NM_153687 | ENSG00000166130 | IKBIP | protein-coding |
| chr4 | 169340102 | 169340124 | - | | 4 | NA | Intergenic | HERVL40-int\|LTR\|ERVL | -69015 | NM_020870 | 57630 | Hs.301804 | NM_020870 | ENSG00000154447 | SH3RF1 | protein-coding |
| chr8 | 141703217 | 141703239 | - | | 4 | NA | Intergenic | Intergenic | 83085 | NR_031636 | 100302147 | | NR_031636 | ENSG00000221768 | MIR1302-7 | ncRNA |
| chr3 | 98627383 | 98627405 | - | | 4 | NA | Intergenic | Intergenic | -33783 | NM_000097 | 1371 | Hs.476982 | NM_000097 | ENSG00000080819 | CPOX | protein-coding |
| chr3 | 161415971 | 161415993 | - | | 4 | NA | Intergenic | Intergenic | -10445 | NR_102265 | 101243545 | Hs.737526 | NR_102265 | ENSG00000240567 | LINC02067 | ncRNA |

| | | | | | | | | | | | | | | | |
|---|---|---|---|---|---|---|---|---|---|---|---|---|---|---|---|
| chr9 | 35248667 | 35248689 | + | 4 | NA | intron (NM_001330653, intron 6 of 39) | L1PA5\|LINE\|L1 | 86686 | NM_006377 | 10497 | Hs.493791 | NM_006377 | ENSG00000198722 | UNC13B | protein-coding |
| chr9 | 93030728 | 93030750 | + | 4 | NA | intron (NM_001083536, intron 15 of 17) | intron (NM_001083536, intron 15 of 17) | -15795 | NR_121565 | 101927954 | Hs.673794 | NR_121565 | | LOC101927954 | ncRNA |
| chr11 | 40534521 | 40534543 | + | 4 | NA | intron (NM_001258419, intron 3 of 6) | MIR\|SINE\|MIR | -241402 | NR_047673 | 57689 | Hs.745123 | NM_020929 | ENSG00000148948 | LRRC4C | protein-coding |
